# Tracks, Maps and Gaps: A Testable Research Definition for Critical Chondrichthyan Areas, European Atlantic Insights

**DOI:** 10.64898/2026.05.01.722225

**Authors:** C Renn, B.J. Ciotti, D.W. Sims, A. Edwards, R.A. Turner, P. Hosegood, E.V. Sheehan

## Abstract

Designing effective spatial management for chondrichthyans (sharks, skates, rays and chimaeras) requires incorporating critical areas, sites essential for population maintenance, such as reproductive and feeding areas. Yet most area-based measures have been developed without consideration of chondrichthyan habitat use. The Important Shark and Ray Area (ISRA) initiative has been pivotal in designating priority areas through a rigorous, consultative process. To complement this, our study offers researchers a testable definition for generating robust evidence to strengthen future critical area delineations and related management decisions. We define critical areas using three criteria: 1) relative frequency of use, (2) extended within-year occupancy and (3) repeated use across years. This framework enables objective comparison among candidate sites and is generalisable across different critical area types. The definition builds upon established early-life-stage habitat concepts and applies these to broader life-history functions. The utility of this framework is then demonstrated through a systematic review of contemporary peer-reviewed literature of critical chondrichthyan areas in the European Atlantic. The review highlighted 62 critical areas with *Strong* evidence and 41 areas of *Moderate* strength evidence, which informed the European Atlantic ISRA selection process. Research effort was concentrated in inshore areas, particularly around the British Isles and Portugal, with biases towards large, threatened and commercially valuable species, whilst chimaeras were notably underrepresented. Early-life stage areas were most frequently identified, whereas resting areas were rarely documented. Evidence patterns and biases are examined in the context of evolving critical area concepts to advance their development and improve the quality and breadth of future research. By outlining a testable definition, identifying key knowledge gaps, and proposing research and reporting guidelines, this work enhances the consistency, comparability, and spatial coverage of future chondrichthyan habitat research to support its application to conservation planning.

## 1. Introduction

After surviving all five of the major extinction events (Bazzi *et al*., 2021), fishes in the Chondrichthyes class (sharks, skates, rays and chimaeras) are now amongst the most threatened taxa due to human-driven pressures (Lascelles *et al*., 2014; Luedtke *et al*., 2023). Over a third of all chondrichthyans are currently at risk of extinction, predominantly due to overfishing; assuming data-deficient species are threatened at the same rate as assessed species (Dulvy *et al*., 2021; Jabado *et al*., 2024). This is concerning considering their key ecological and social functions, including food provision, nutrient transport and capacity for food web structuring (Williams *et al*., 2018; Barbosa-Filho *et al*., 2019; Dedman *et al*., 2024; Hammerschlag *et al*., 2025), and their cultural value to a wide range of groups, including anglers and Indigenous communities (Skubel, Shriver-Rice & Maranto, 2019; Scotts *et al*., 2023).

To curb such declines, multiple conservation strategies are available, spanning the remit of both nature conservation and fisheries management (Booth, Squires & Milner-Gulland, 2020). Whilst the relative efficacy of different management strategies remains subject to debate (Shipp, 2003; Shiffman & Hueter, 2017; Porcher, Darvell & Cuny, 2019; Jorgensen *et al*., 2022), a foundational understanding of chondrichthyan habitat use is a common prerequisite for most interventions. For instance, fishing measures, such as gear restrictions and seasonal closures, are often assigned regionally, particularly in inshore regions (e.g. local byelaws are established regionally in English coastal waters (Solandt, Clark & Coulthard, 2025)). Furthermore, habitat use information is beneficial for achieving ecosystem-based fishery management, whereby ecological benefits are extended beyond individual target species to the wider ecosystem (Pikitch *et al*., 2004; Holsman *et al*., 2020). However, the need for habitat use information is most apparent when designing new and/or improving existing area-based management (Kelleher, 1999; Escalle *et al*., 2015).

Various factors influence the efficacy of spatial management for chondrichthyans; these include social drivers, such as engagement with and resultant support from the local community, and whether successful fishing interventions are introduced (White *et al*., 2015; MacKeracher, Diedrich & Simpfendorfer, 2019; Leurs *et al*., 2021). Such social drivers should not be neglected as they often represent the key limiting factors in many implemented marine protected areas (MPAs); however, biological and physical factors limit the theoretical maximum efficacy of an optimally designed MPA with unlimited management resources. These biophysical factors include both habitat characteristics, such as inherent sensitivity to and overlap with ongoing fishing pressure (Murawski *et al*., 2000), and species characteristics, including species’ mobility and tendency to form close habitat associations (Breen, Posen & Righton, 2015; Oh *et al*., 2017).

When a species demonstrates strong associations with an area (Thorburn *et al*., 2015; Dodd *et al*., 2022), protection of that site may yield considerable population benefits, even for species with wide-ranging movement patterns, otherwise presumed unsuitable for spatial management (Doherty *et al*., 2017a; Gallagher *et al*., 2021). Protecting any area in which a species spends a significant portion of time reduces mortality risk, assuming appropriate protective measures are implemented (Sala *et al*., 2018). However, these benefits are amplified when the habitat is associated with a critical life stage or behaviour, as protection not only increases survival rates but supports demographic processes essential for population maintenance and growth (Beck *et al*., 2001; Da Silva *et al*., 2021). Accordingly, area-based management is considered most effective when it encompasses *critical areas,* meaning sites disproportionately supporting population maintenance through crucial life-history functions such as feeding and reproduction (Le Port and Lavery, 2012; Camaclang *et al*., 2015; Da Silva *et al*., 2021; Hyde *et al*., 2022).

Unfortunately, chondrichthyans are rarely included in spatial management considerations, meaning few MPAs, including those established specifically for chondrichthyan protection, have incorporated prior knowledge on critical areas into their design (Ward-Paige, 2017; Gallagher *et al*., 2021). As a result, area-based management size and placement rarely align effectively with chondrichthyan habitat use (Mouton *et al*., 2025; García-Rodríguez *et al*., 2025). Meanwhile, the lack of meaningful fishing interventions in the vast majority of existing MPAs would provide little benefit to threatened chondrichthyans (Sala *et al*., 2018; Bridier, Wauchope & Vad, 2025). Furthermore, whilst generic information on species’ geographic range is widely available, species-specific abundance estimates at different life-history stages (e.g. neonates vs adults) and during vital functions (e.g. feeding, mating, migration) remain limited (Rooper *et al*., 2019). When critical area information does exist, coverage is typically greater for larger, flagship species [such as those listed on the Convention on Migratory Species (CMS)(García-Rodríguez *et al*., 2025)].

To address this evidence deficit, this review aims to support researchers in designing studies of maximal value to spatial management frameworks while identifying and guiding future work toward key data gaps. Specifically, the review seeks to:

1. Offer a testable working definition to support the identification of critical chondrichthyan areas in future research;
2. Demonstrate the framework by identifying potential critical areas from peer-reviewed literature in the European Atlantic;
3. Highlight key knowledge gaps and limitations in current research practices.

The framework is informed by key critical chondrichthyan area concepts which are presented through a timeline of the field’s development and connections to essential fish habitat literature. Critical areas are then identified through a systematic review of peer-reviewed literature for a selection of UK-priority chondrichthyan fishes in the European Atlantic. This region was selected as a model system due to its location in the most-studied ocean basin (Potter & Pearson, 2023) and to extract useful data for the European Atlantic ISRA delineation process (Hyde *et al*., 2022a; Jabado *et al*., 2025). Standardised reporting and research guidelines are then proposed to promote consistent, high-quality future research.

### 1.1 Key Terminology

*Critical*, *important*, and *essential* are often used interchangeably to describe ecologically significant areas for chondrichthyans. Here, we adopt *critical* as the more widely used term in the encompassed literature for this review, and to distinguish this review’s findings from the *Important Shark & Ray Areas,* which are exclusively designated to inform conservation planning through the expert-consultative ISRA process (Hyde *et al*., 2022). A further distinction is needed between *critical habitat* and *critical area*. *Habitat* definitions describe the observed locations where organisms live, encompassing both descriptive ecosystem categories (e.g., seagrass, mangrove) and the measurable spatially and temporally defined abiotic and biotic variables that support a species’ occurrence (Hayes, Ferreri & Taylor, 1996; Hall, Krausman & Morrison, 1997; Sass, Rypel & Stafford, 2017).

*Critical habitats* represent a subsample of these conditions which support vital life-history functions and behaviours, where demographic rates are disproportionately influenced (U.S. Congress, 1973; Camaclang *et al*., 2015; Rylander *et al*., 2020). Here, we use *critical area* to denote a three-dimensional portion of habitat in a specific geographic location that may be suitable for area-based management [such as ISRAs (Hyde et al., 2022)], therefore representing a subset of critical habitats. The two concepts are closely linked: identifying critical habitat preferences can help predict the location of yet-undiscovered critical areas, while known critical areas are likely to share common habitat characteristics; however, such characteristics may vary contextually, reflecting the species’ realised niche (Hutchinson, 1957; Rodriguez-Cabal, Barrios-Garcia & Nunez, 2012) within that area. This review focuses primarily on identifying critical areas; species’ critical habitat preferences are discussed only briefly for supplementary context.

## 2. Field development

Significant advancements in critical area research for fishes have been made over the past 50 years. A major cornerstone was the 1996 amendment of the Magnuson-Stevens Fishery Conservation and Management Act (U.S. Department of Commerce, 1996; Benaka, 1999). The amendment required fishery management plans to include the identification of Essential Fish Habitat (EFH), defined as “*those waters and substrata necessary to fish for spawning, breeding, feeding, or growth to maturity*” (U.S. Department of Commerce, 1996; p. 6). The act also required EFH threats and potential protective measures to be identified, signalling a shift towards ecosystem-based fishery management (U.S. Department of Commerce, 1996; Pikitch *et al*., 2004).

A four-level information framework: (1) presence/absence, (2) density, (3) growth, reproduction, and survival, and (4) production, was proposed in the EFH guidelines under 50 CFR 600 (NOAA Fisheries, 2002) to support identification of EFH under the Magnuson–Stevens Act. Although this hierarchy offered a useful starting point, EFH designations in practice often relied on the lower tiers of evidence, falling short of the higher-level metrics (Levels 3–4) needed to demonstrate contributions to population productivity, which remains an issue today (NOAA Fisheries, 2002; Grüss *et al.,* 2017; Zu Ermgassen *et al.,* 2021). Likewise, early nursery area examinations were constrained by vague definitions and assumptions, often equating juvenile presence or density with habitat importance (Levin & Stunz, 2005; Heupel *et al*., 2019). This undermined the motivation of EFH identification to direct limited conservation resources towards the most important fish habitats (Beck *et al*., 2001; Levin & Stunz, 2005). Researchers, therefore, sought clearer conceptual frameworks to enable more rigorous, quantitative identification of the habitats most critical to population maintenance.

A major advance in identifying nursery habitat was the testable Nursery-Role Hypothesis by Beck *et al*. (2001, p. 635) *“A habitat is a nursery for juveniles of a particular species if its contribution per unit area to the production of individuals that recruit to adult populations is greater, on average, than production from other habitats in which juveniles occur”.* This characterised nursery areas as a subset of overall juvenile habitats, distinguishable for their disproportionate contribution to the adult population (Ciotti *et al*., 2025). Further modifications improved usability and management relevancy. The concept of Effective Juvenile Habitat emphasised total site contribution to recruitment, over per-unit-area contribution, identifying the most important areas to population maintenance regardless of efficiency (Dahlgren *et al*., 2006). A further conceptual shift, “seascape nursery concept,”, acknowledged nurseries as dynamic mosaics of functionally connected habitat patches rather than discrete, isolated areas [(Nagelkerken *et al*., 2015), see also Ciotti *et al. (*2025) for a more detailed discussion of this field].

Until 2007, research on chondrichthyan nursery areas largely developed separately from teleost literature and typically exhibited similar flaws: definitions were ambiguous, methods inconsistent, lacked testable hypotheses and often inferred habitat importance merely from juvenile abundance or occurrence (e.g., Olsen, 1953; Blackburn, McCandless & Pratt, 2007; see Heupel *et al.,* 2007 for further detail). To address this, the shark nursery area definition was formalised by Heupel *et al*. (2007), taking influence from key teleost nursery literature (Beck *et al*., 2001). Recognising the difficulty of quantifying juvenile-to-adult recruitment in slow-growing, threatened and highly mobile chondrichthyans, Heupel *et al*. (2007) proposed three practical criteria based on a early juvenile density, tendency to remain or return, and interannual habitat use [residency meaning the largely uninterrupted occupancy of an area (Chapman *et al*., 2015)]. These criteria serve as proxies for the relatively higher production of nursery areas compared with other non-nursery early juvenile habitats, whilst offering feasible metrics suited to chondrichthyan life histories. Although this framework improved chondrichthyan nursery delineation, biases toward tropical and coastal species have been recognised (Heupel *et al*., 2019).

To better encompass oviparous chondrichthyan EFHs, the term *egg case nursery* was coined (Hoff, 2016) to differentiate sites with high densities of developing eggs from important early juvenile habitats, following observations of spatial partitioning between these two life stages (Hoff, 2010, 2016; Hunt, Lindsay & Shahalemi, 2011). Martins *et al*. (2018) further streamlined the *egg nursery area* concept to mean areas with: (1) high densities of eggs attached to benthic structures [effectively combining criteria 1 and 2 from Hoff *et al*. (2016)], (2) repeated annual use by adults for oviposition, and (3) departure of newborns or young-of-the-year soon after hatching.

While early life stage concepts remain comparatively more developed than other critical chondrichthyan areas, research advancements and theory developments have enhanced our understanding of habitat use during key reproductive events, such as gestation (Jirik & Lowe, 2012) and mating (Pratt & Carrier, 2001). Pregnant individuals often behaviourally thermoregulate to reduce gestation time, increase offspring size, or avoid males through sexual habitat segregation, explaining frequent use of warm, shallow waters during gestation (Wallman & Bennett, 2006; Hight & Lowe, 2007; Wearmouth & Sims, 2008; Jirik & Lowe, 2012).

Broader advances were made with the International Union for Conservation of Nature Species Survival Commission (IUCN SSC) Shark Specialist Group, ISRA framework, which identifies habitats of importance anywhere chondrichthyans naturally occur (Hyde *et al*., 2022). The criteria define critical areas based on: A) species vulnerability (whether a species is threatened with extinction) B) range restriction, C) life-history function (reproduction, feeding, resting, movement, aggregations), D) special attributes [areas supporting chondrichthyans with distinct characteristics or elevated chondrichthyan diversity, see Hyde *et al*. (2022) for further detail]. These evidence-based criteria are largely descriptive rather than hypothesis-driven, providing necessary flexibility within the participatory, consultative ISRA designation process; however, each proposed ISRA must undergo rigorous peer-review to demonstrate regular or predictable use (IUCN SSC Shark Specialist Group, 2024). Criterion C (Life-History) encompasses all EFH categories (Benaka, 1999), and incorporates the nursery (Heupel, Carlson & Simpfendorfer, 2007) and egg nursery (Martins *et al*., 2018) area concepts. However, an ISRA designated under Sub-criterion C1 (Reproductive Area) need not satisfy all nursery or egg nursery area criteria.

Recent work has also clarified chondrichthyan aggregation areas, addressing inconsistencies in earlier usage (Jacoby, Croft & Sims, 2012; McInturf *et al*., 2023). Under this framework, aggregations must: 1) contain at least two individuals, 2) occur at a scale where individuals can simultaneously detect and respond to the same driver, 3) demonstrate deliberate use of that common driver (McInturf *et al*., 2023). Whilst this has resolved much of the ambiguity surrounding this term, some researchers recommend a minimum of three individuals to avoid conflating incidental co-occurrence with true aggregation, a threshold adopted by the ISRA framework (IUCN SSC Shark Specialist Group, 2024). Likewise, ISRA aggregation designations do not require identification of a common driver (Hyde *et al*., 2022).

### 2.1 A testable definition for identifying critical areas through research

To help researchers generate consistent and comparable evidence for spatial planning processes, in line with the concepts outlined, we propose a testable, and generalisable definition for critical chondrichthyan areas. Building on the nursery framework proposed by Heupel *et al*. (2007), we show that, with minor adaptation, the evaluation of an area’s importance through density, occupancy, and interannual use can be generalised to habitats beyond nursery areas (Table 1). Relative density or frequency of use (R1) reflects the number of individuals exhibiting a preference for the site; meanwhile, within-year occupancy (R2) and repeated interannual use (R3) indicate the relative time an area is occupied. Since direct measurement of recruitment is challenging for slow-growing or rare species with relatively inaccessible archival growth tissues, high relative density or frequency of use combined with elevated occupancy can serve as a proxy for higher productivity [i.e., adult recruitment (Heupel *et al.,* 2007)].

**Table 1.** Testable critical area definition proposed, including three key requirements (R1-R3), building on the Nursery concept (Heupel *et al*., 2007). All requirements must be associated with the key life-stage or behaviour, relating to critical area function.

| Nursery Criteria (Heupel *et al.*, 2007) |  | Critical Area Definition (Proposed by this review) |  |
| --- | --- | --- | --- |
| **C1** | Neonates or young-of-the-year (YOY) are more commonly encountered in the area than in other areas | **R1** | The key life-stage or behaviour is **more frequently encountered in this area** compared to other areas, as demonstrated by effort-standardised observations |
| **C2** | Neonates or YOY tend to remain for extended periods | **R2** | **Extended within-year occupancy** during the key life-stage or behaviour is observed |
| **C2 & C3** | Habitat is repeatedly used across years by Neonates or YOY (C3); **or** Sharks tend to return (C2) | **R3** | **Repeated interannual occurrence** during the key life-stage or behaviour is observed (not necessarily in successive years) |

To avoid ambiguity, we explicitly separate habitat use within a single year (within-year) from repeated habitat use over multiple years (interannual). For within-year habitat use, we opt for the term *extended occupancy* rather than *residency* (Chapman *et al*., 2015), to include both continuous and intermittent use, at both species– and individual-level, since all variations are relevant to management decisions. In this review, individual-level use is considered marginally more significant, as it confirms that individuals occupy the area for extended periods, whereas species-level intra– or interannual occupancy may simply reflect sequential use by different individuals, each spending only limited time in the area. Further, individual-level interannual use may be met either by *site fidelity,* where individuals leave and return after a prolonged period of absence (Speed *et al*., 2011; Chapman *et al*., 2015), or by continuous presence in the area across years.

To capture the diversity of critical area functions, the first nursery criterion of Heupel *et al*. (2007) was modified to reflect either: (1) greater relative densities of individuals at a key life stage (e.g., reproductively active adults in mating areas, pregnant females in gestation areas) or (2) more frequent observations of key behaviours (e.g., feeding, mating, resting). Critically, to meet this requirement, observation data must be standardised by effort to ensure higher counts reflect biological patterns rather than sampling intensity. The proposed requirements (Table 1) are compatible with the ISRA framework, which designates ISRAs based on regular or predictable habitat use, and increasingly requires a spatial comparison with adjacent areas to signify elevated importance (IUCN SSC Shark Specialist Group, 2024).

The requirements are readily testable using common research methods and survey designs and are applicable across taxa, with potential suitability to non-chondrichthyan taxa with similar life histories. To maintain broad applicability across critical area types, the definition intentionally avoids habitat-specific qualifiers [e.g. neonates or Young-of-the-year (YOY) or egg contact with benthos]. This more inclusive approach has the advantage of capturing all critical areas of potential relevance to management (e.g. all critical immature areas, rather than just nursery areas). Habitat-specific frameworks can then be to further classify critical areas according to more specific functions [e.g., egg nurseries: (Martins *et al*., 2018); aggregations: (McInturf *et al*., 2023)]. In this review, such habitat-specific frameworks were incorporated into the quality assessment process (Section 3.7).

### 2.3. Thresholds of importance applied in this review

There is no universal level of habitat use that defines importance, as this varies between species and populations. For some species, a critical reproductive event may occur in a single day; whilst others may regularly gather in large numbers for months across multiple areas (Pratt & Carrier, 2001; Espino *et al*., 2022; Pratt *et al*., 2022). Future research should establish region– and species-specific baselines to define thresholds of normal *vs.* important habitat use. Each requirement could then be assessed relative such baselines.

For this review, we defined generic thresholds of extended occupancy to enable rapid, objective comparison among candidate sites. To account for some interspecific differences in mobility, we applied two thresholds: ≥ two days for highly mobile pelagic sharks and ≥ two months for less mobile benthic and benthopelagic species. Inclusive limits, signifying the lower bounds of observed residency, were applied to both taxonomic groups to facilitate a comprehensive search for all potential critical areas. Telemetry studies in the northeast Atlantic show that benthic and benthopelagic elasmobranchs often exhibit multi-month residency, whereas transient use typically occurs over days to weeks (Thorburn *et al*., 2015; Simpson, Humphries & Sims, 2021; Papadopoulo *et al*., 2023). Thus, a two-month threshold provides a pragmatic and conservative distinction between transient visits and sustained use that may indicate critical habitat function. It also aligns with the lower bound for which chondrichthyan seasonal habitat use is typically defined and seasonal management is recommended (Saïdi, Bradaï & Bouaïn, 2008; Narayanakumar *et al*., 2017; Pratt *et al*., 2022; Crespo *et al*., 2024).

Pelagic species typically show residency over shorter periods of days to weeks (Biais *et al*., 2017; Sims *et al*., 2022; Thorburn *et al*., 2024), though occupancy can extend into months at broader spatial scales (Biais *et al*., 2017; Doherty *et al*., 2017a; Bortoluzzi *et al*., 2024; Jung *et al*., 2024a). We therefore adopted a two-day (48-hour) lower bound as an inclusive, evidence-based threshold for identifying potentially important areas for the highly mobile species assessed here (*Cetorhinus maximus, Lamna nasus, Alopias vulpinus*). Limited available data for these species in the Northeast Atlantic indicate that *C. maximus* has an average occupancy of 0.94 days per area (Thorburn *et al*., 2024), making a two-day threshold sufficient for filtering out routine transience. Multi-day occupancy also indicates a pause in the long-distance movements typical of migrating pelagic sharks (e.g., daily travel in *L. nasus:* mean ≈ 28 km, max ≈ 200 km; Biais *et al*., 2017; Bortoluzzi *et al*., 2024) whilst remaining inclusive enough to capture brief but meaningful behaviours such as multi-day courtship aggregations in *C. maximus* (Sims *et al*., 2022).

## 3. Systematic review methods

### 3.1 Species selection

This systematic review provides an overview of critical chondrichthyan areas in the European Atlantic, focusing on a subset of priority UK species to illustrate the application of our definition and examine general trends, rather than provide an exhaustive review of all areas [for which consult the recent European ISRA designations (Jabado *et al*., 2025)]. Focal species for this review include the top 21 priority elasmobranchs from the British Isles (McCully Phillips, 2020), highlighted based on several factors including biological vulnerability, conservation interest, commercial importance, and the importance of UK waters to the species range; to capturing the diverse vulnerabilities and contributions of chondrichthyans to social-ecological systems. To increase attention to the Chimaeriformes order, all chimaeras listed in the UK National Plan of Action were also included (Fowler, Mogensen & Blasdale, 2004). The complete list of 27 focal species is in Table 2.

**Table 2.**
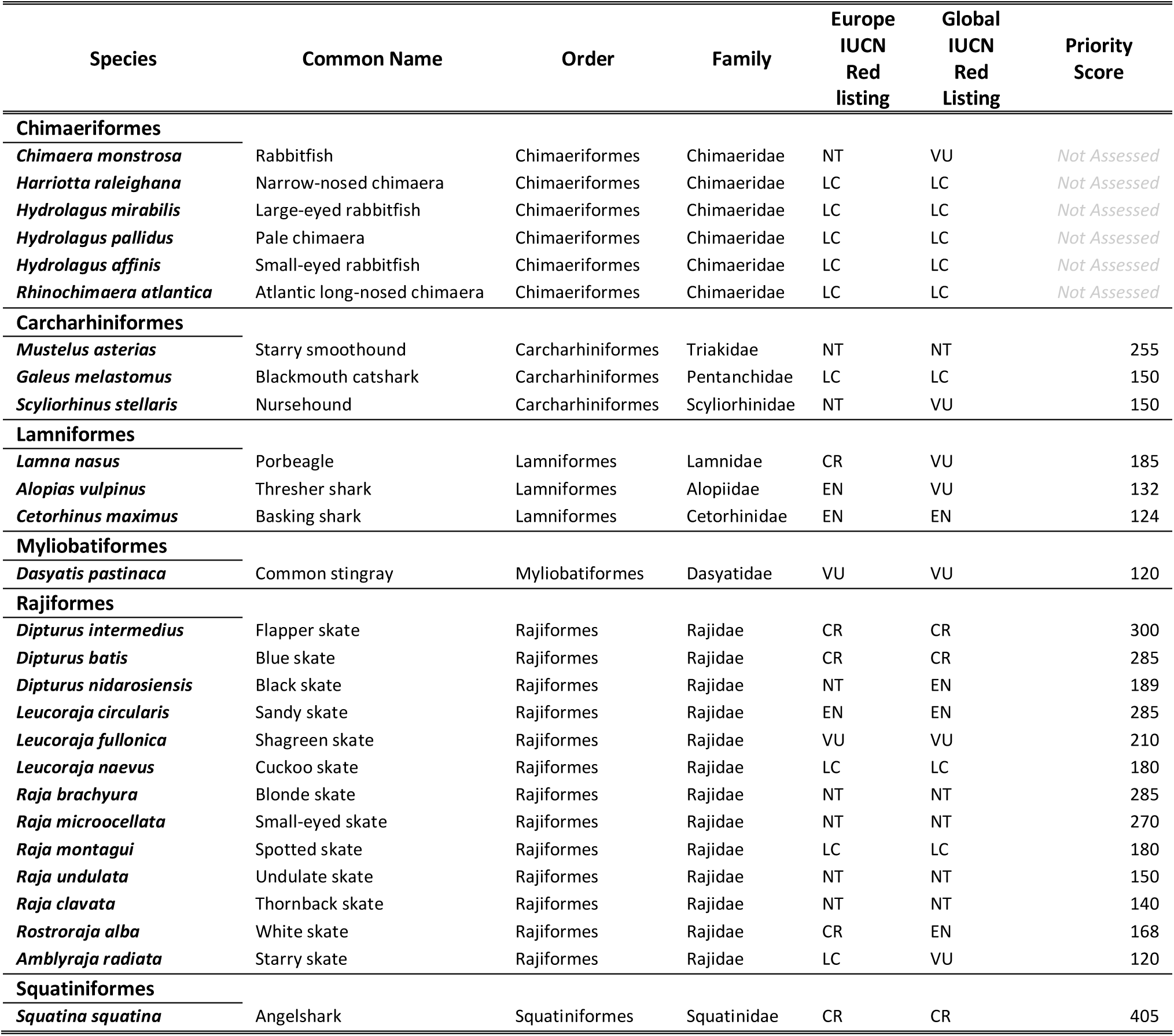
Selected study species, including the top 21 UK priority elasmobranch species (McCully Phillips, 2020) and six chimaeras listed in the UK NPOA (Fowler *et al*., 2004) including their European and global IUCN Red List of Threated Species statuses (IUCN, 2025; See Table S1 for species-specific assessments used). The priority scores represent each species’ overall conservation priority, with higher scores indicating greater priority (McCully Phillips, 2020).

### 3.2 Critical areas

The critical area categories encompassed by this review are outlined in Table 3. These span the ISRA Criteria C (Life History) and D [Special Attributes (Hyde *et al*., 2022)]. Areas of importance for threatened species [ISRA Criterion A (Vulnerability)] were examined indirectly via the distribution of critical areas per IUCN Red List category, using the global-level assessments, as the most current assessments (IUCN, 2025; Table S1). However, none of the focal species are considered range-restricted (ISRA Criterion B), whereby distribution is limited to one or two adjacent Large Marine Ecosystems (Sherman and Alexander, 1986; Hyde *et al*. 2022).

**Table 3.** Critical area categories studied, their related ISRA criteria and additional habitat-specific frameworks.

| Related ISRA Criterion | Habitat Categories Studied | Additional Useful Existing Frameworks |
| --- | --- | --- |
| **C1 Reproductive Areas** | High-use Area (Undefined Function) | / |
|  | Immature Area | Heupel et al., 2007 |
|  | Egg-laying Area | Martins et al., 2018; Hoff et al., 2016 |
|  | Viviparous Gestation or Parturition Area | / |
|  | Mating or Courtship Area | / |
| **C2 Feeding Areas** | Feeding Area | / |
| **C3 Resting Areas** | Resting Area | / |
| **C4 Movement** | Movement Area | / |
| **C5 Undefined Aggregations** | Undefined Aggregation | McInturf et al., 2023 |
| **D2 Diversity** | Area of Diversity | / |

Reproductive habitats (ISRA Sub-criterion C1) were subdivided into mating/courtship areas, egg-laying areas, immature (free-swimming stage) areas, and viviparous gestation/parturition areas to aid closer examination, though some overlap is likely, particularly between viviparous gestation/parturition and immature areas. Oviparous gestation areas were not examined independently from egg nurseries, as egg-bearing females must often remain near egg-laying areas for prolonged oviposition periods (Ellis *et al*., 2005; Thorburn *et al*., 2021a). Thus, observations of large, mature, egg-bearing females were instead classified as evidence of egg-laying areas. Areas of high relative use (meeting requirements R1 – R3) without a clear function were also recorded to capture potentially important areas of unknown use. Both immature areas and egg-laying areas were further subdivided in the discussion to identify potential nursery areas and egg nursery areas respectively. Potentially important Movement Areas (ISRA Sub-criterion C4), such as migratory corridors, were not examined in depth because: their typically large size limits relevance to area-based management and their identification requires analyses beyond the scope of this review, owing to high individual variability in shark movements (Cameron *et al*., 2019; Bortoluzzi *et al*., 2024; Sequeira *et al*., 2025). However, relevant movement studies are signposted in Table S2.

### 3.3 Search strategy

The search strategy followed PRISMA guidelines (Page *et al*., 2021; Figure 1) and targeted Scopus and Web of Science from January 2010 to January 2025 to capture the previous 15 years of research, in line with ISRA guidance (IUCN SSC Shark Specialist Group, 2024). Searches were restricted to English-language publications and designed to cover all priority species (Table 2) and critical area categories (Table 3). Final search strings (Table S3) combined each species’ accepted Latin name with all habitat terms, using an asterisk to capture relevant word variants (e.g., migrate, migratory, migration). Additional detail on the justification for search string decisions (e.g. exclusion of common names) and the handling of taxonomic ambiguity is found in the supplementary materials (Table S3; accompanying text).

**Figure 1.**
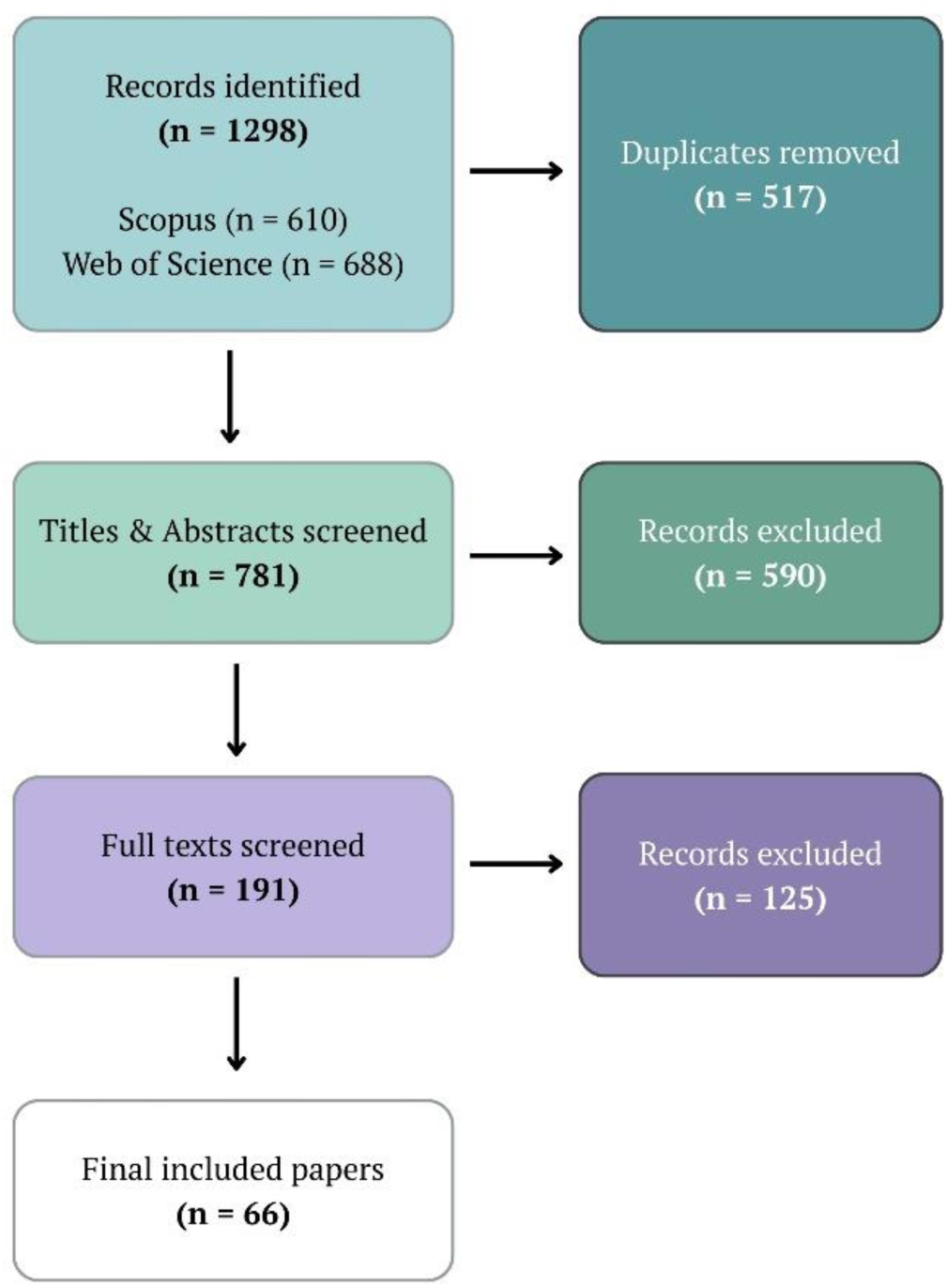
Screening and selection process of included studies. Studies were subject to dual screening.

### 3.4 Inclusion and Exclusion criteria

The Population, Concept, Context (PCC) framework, suitable for scoping reviews, was used to refine the research question [Table S4 (Peters *et al*., 2020)]. Titles and abstracts were independently screened using inclusion and exclusion criteria designed to for compatibility with the critical area definition (Table 1) and ISRA framework (IUCN SSC Shark Specialist Group, 2024), and to capture studies identifying potential critical areas, both explicitly and incidentally. Full-text screening followed the same criteria, and the process was repeated after two months to ensure repeatability (Cherry, Boland & Dickson, 2024). Final included studies are listed in Table S5.

### 3.5 Data extraction

For each included study (and associated supplementary materials), all areas in which a key life-stage or behaviour related to any of the critical area types was observed were recorded, along with the specific requirements (Table 1) that each observation satisfied. To assess the spatial extent of research effort, the ICES subareas (FAO Subdivision 27) and CECAF areas (FAO Subdivision 34) of focal chondrichthyan observations were noted. Evidence was only extracted when species-specific, unless explicitly noted as important for diversity. To acknowledge the potential importance of all juvenile stages to population recruitment (Kinney & Simpfendorfer, 2009), evidence for all free-swimming immature life stage areas was extracted. The specific immature stage [neonate, YOY, juvenile or immature (no lower size bound) stage; Figure 2] of the smallest observed individuals was then recorded using size-at-birth, size-at-one-year or first observable cohort (Ellis *et al*., 2024) and size at maturity (Ebert & Dando, 2021) to aid further area categorisation (Figure 3; Table S6). Juvenile is used here to indicate the life stage following YOY until maturity, which here also encompasses subadult stages (Hoyos-Padilla *et al*., 2016), as subadult size thresholds are not widely defined. These finer categorisations allowed immature areas to be further broken down into their constituent area types (Figure 3), including areas meeting strict nursery and egg nursery area definitions (Heupel, Carlson & Simpfendorfer, 2007; Martins *et al*., 2018).

**Figure 2.**
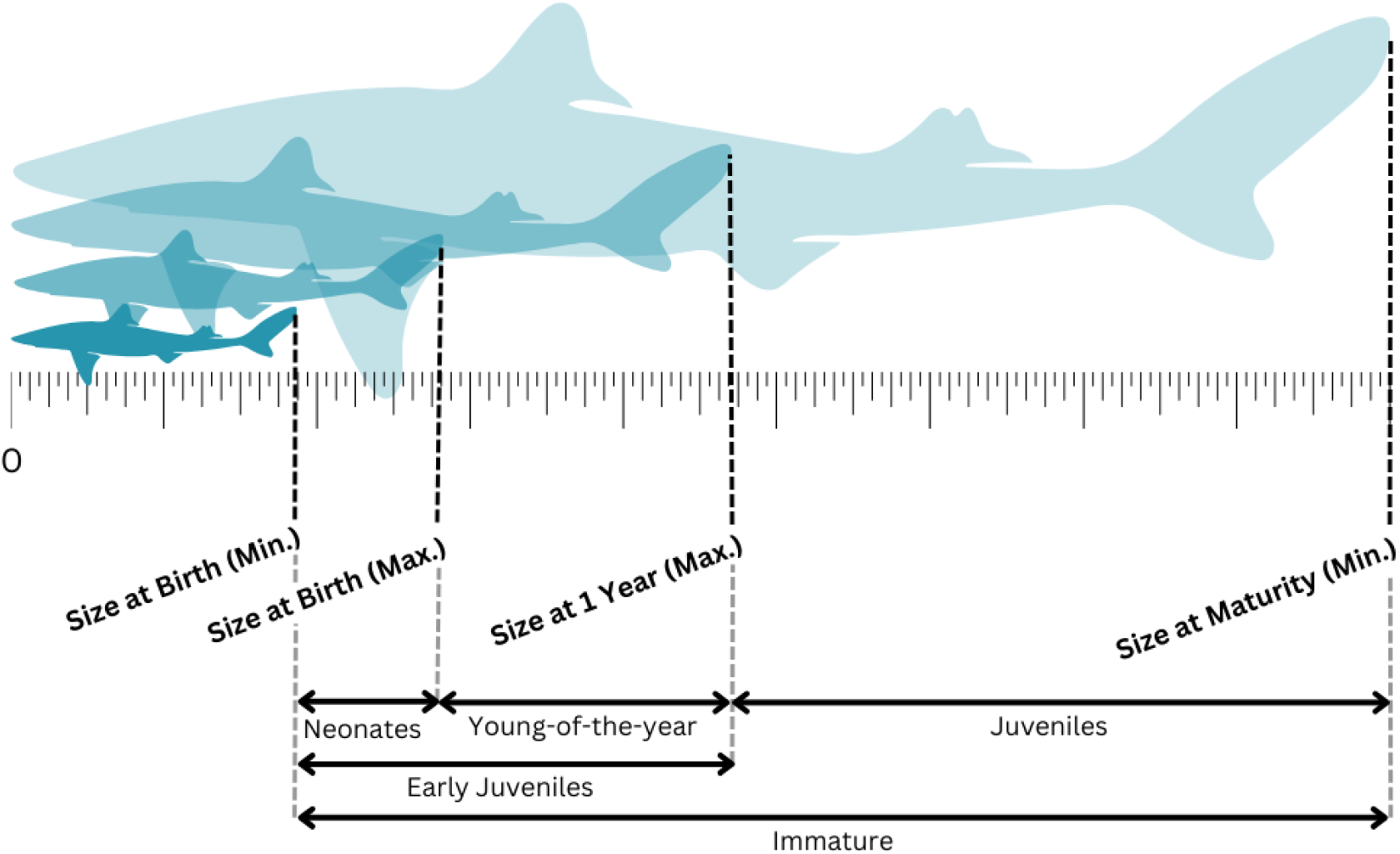
Free-swimming immature life stage categories used in this review.

**Figure 3.**
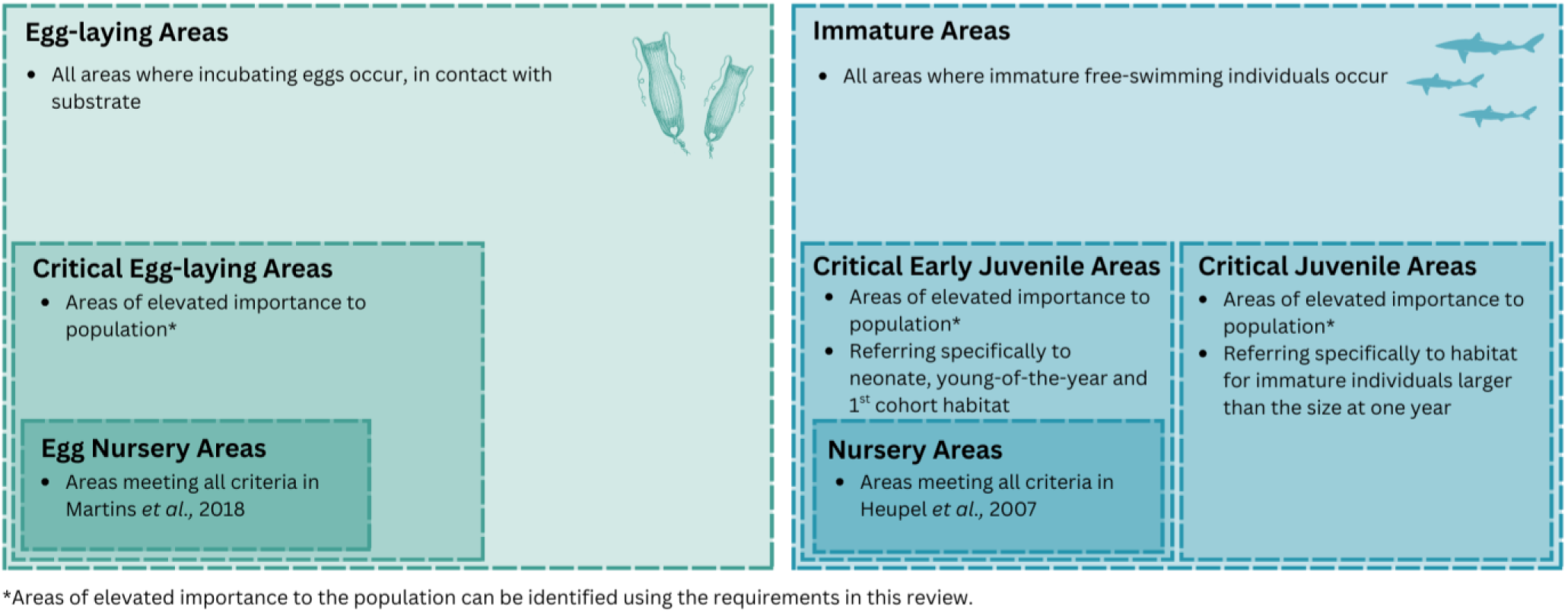
Conceptualisation of the nested early life stage area definitions used in this review for both egg-laying areas and free-swimming immature areas and the increasing evidence requirements needed.

### 3.6 Evidence Evaluation

Because study selection and evidence extraction were designed to maximise inclusivity, evidence strength for each potential critical area was assessed by the number of requirements met (Table 4). Scores of ≥ 2 were considered *Strong* evidence, 1.5 was considered *Moderate* (meaning consistent use by individuals was confirmed), and < 1.5 was classified as *Weak/ Cannot Assess*. Requirements were only satisfied when habitat use evidence was clearly linked to the key behaviour (e.g. aggregating, feeding, resting) or life stage (e.g. eggs, neonates, subadults, pregnant females). To ensure this, direct observations of each behaviour or life-stage were necessary for evidence to be classified as *Strong*; otherwise, evidence was capped at *Moderate* (Figure S1).

**Table 4.**
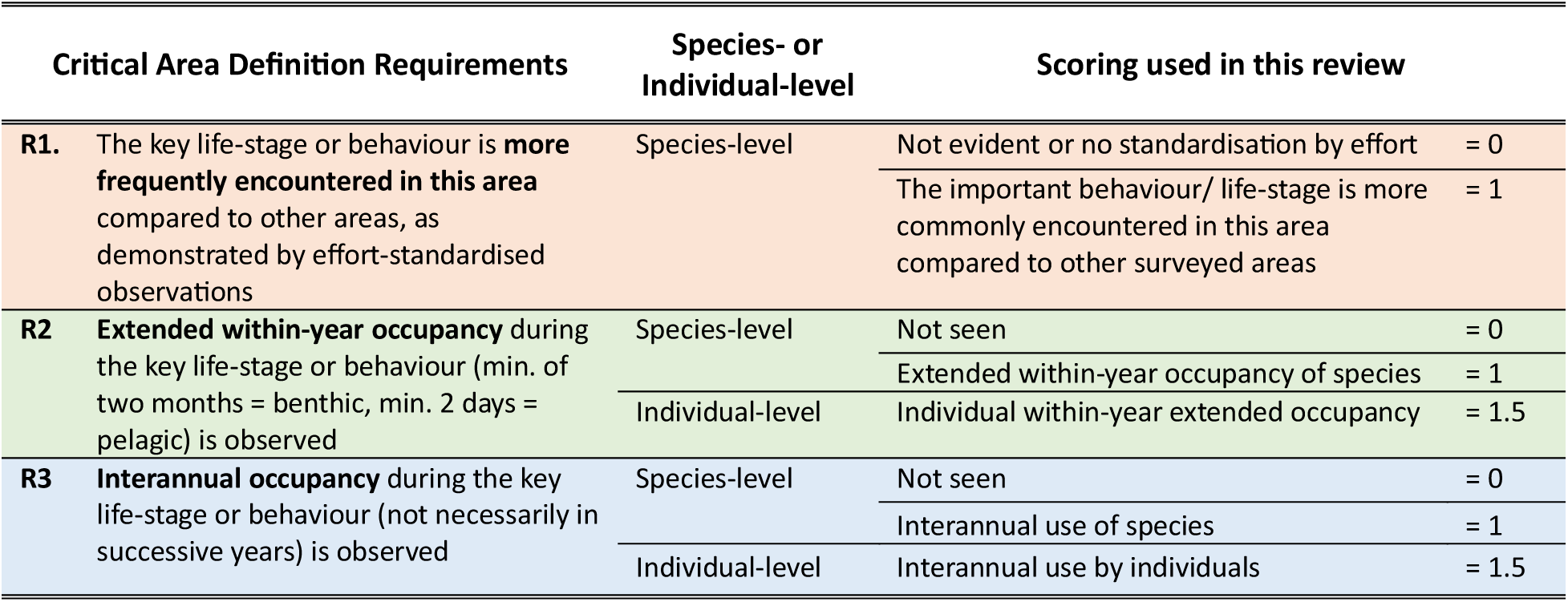
Critical area definition requirements and the scoring system applied by this review.

### 3.7 Risk of bias

To comply with PRISMA standards, a quality assessment tool was adapted based on the Cochrane risk-of-bias tool (Higgins *et al*., 2011; Page *et al*., 2021; Cherry *et al*., 2024), incorporating both generic and critical-area-specific criteria informed by existing habitat frameworks (Heupel *et al*., 2007; Martins *et al*., 2018; Hyde *et al*., 2022; McInturf *et al*., 2023). Evidence was scored against these criteria and converted into a percentage of the total possible score. Generic criteria included evidence type (empirical *vs*. anecdotal), survey effort (high or low), representation of the focal behaviour or life stage in the sample, and whether behaviours/ life history functions were directly observed or inferred (Figure S1).

Additional critical-area-specific criteria were introduced where necessary to ensure correct interpretation of area function. When the type of evidence supplied could not unequivocally confirm area function, overall evidence score was capped at *Moderate.* For instance, for aggregations, two individuals were considered adequate evidence (McInturf *et al*., 2023), however ≥3 individuals were required for *Strong* evidence, since this review did not explicitly examine aggregation drivers (aligning with the ISRA process; Hyde *et al*. 2022). To standardise decisions and minimise bias, an initial subset of areas was independently dual-assessed and the agreed thresholds were then applied to all remaining studies (see Table S7 and Figure S1 for more detail).

### 3.8 Overall Strength of Evidence

To assess the overall strength of evidence for each potential critical area, the total requirement score was combined with the risk of bias assessment (Table 5). The evidence was restricted to *Weak/ migraCannot Assess* when the risk of bias was high (score < 45 %). However, such areas meeting ≥ 1.5 requirements were labelled *Priority* areas for further research due to potential high use.

**Table 5.**
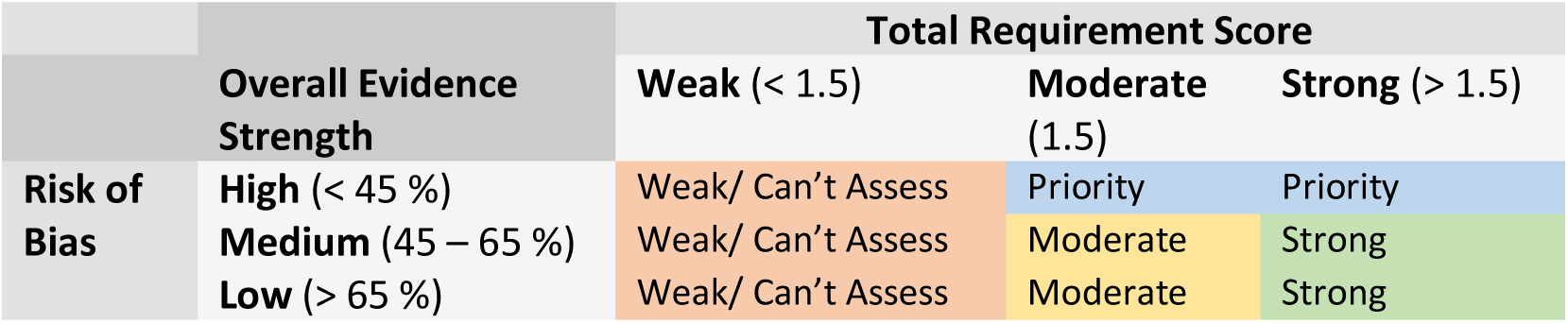
Matrix demonstrating how *Risk of Bias* and *Total Requirements Met* are combined to determine *Overall Evidence Strength. Priority* areas are those which could be easily upgraded to *Strong* or *Moderate* and therefore should be prioritised by future research.

|  | Overall Evidence Strength | Total Requirement Score |  |  |
| --- | --- | --- | --- | --- |
|  |  | Weak (< 1.5) | Moderate (1.5) | Strong (> 1.5) |
| Risk of Bias | High (< 45 %) | Weak/ Can't Assess | Priority | Priority |
|  | Medium (45 – 65 %) | Weak/ Can't Assess | Moderate | Strong |
|  | Low (> 65 %) | Weak/ Can't Assess | Moderate | Strong |

When study areas overlapped completely, their evidence was pooled, generating a single overall evidence score based on the mean Risk of Bias score and the combined requirements met. In cases of only partial overlap evidence was kept separate. These results were then visualised in a matrix plot illustrating the distribution of research effort across species and critical area types.

### 3.9 Mapping areas

All potential critical areas scoring ≥ 1 requirement were mapped using QGIS (QGIS.org, 2025) at a 1° × 0.5° resolution using an extended ICES statistical rectangle system. The grid was expanded from FAO (Food and Agriculture Organisation of the United Nations) region 27 into FAO Area 34 (Eastern Central Atlantic) and labelled following a continuation of the ICES rectangle alphanumeric code to provide consistent mapping resolution across the European Atlantic region.

During mapping, each species was represented only once per ICES rectangle for each critical area type (e.g., Aggregation, Feeding Area) to assess the total number of qualifying species within each rectangle and to identify multispecies critical areas, while avoiding artificial inflation of importance caused by multiple smaller, same-species areas occurring within the same rectangle. To achieve this, multiple species records per rectangle were ranked by evidence strength, and only the highest-ranked record for each combination of species, habitat, and evidence strength was kept. However, to provide a more detailed understanding of critical area boundaries, all areas are also described and presented at their extracted spatial resolution in Table S8. Differences between map and matrix spatial resolutions mean that the total number of identified areas does not match exactly between outputs; likewise, each ICES rectangle may contain multiple finer-resolution habitat features for a given species.

## 4. Results & Discussion

### 4.1. Research Effort Trends

The literature search and selection process retrieved 66 peer-reviewed articles (Figure 1) from the two searched databases. One article represented a correction to an included study (Thorburn *et al*., 2021b) and was therefore not subject to full data extraction. While the search period included articles from 2010 – 2024, studies satisfying all selection inclusion criteria were limited to 2012 – 2024 (Figure 4). The selection process excluded articles with data older than 15 years to align with the ISRA requirements to assess contemporary habitat use (Hyde *et al*., 2022), which disproportionately affected earlier publications. Despite this, there was a clear increase in publications identifying potential critical chondrichthyan areas over time, especially beyond 2019, likely signifying a growing interest in this field.

**Figure 4.**
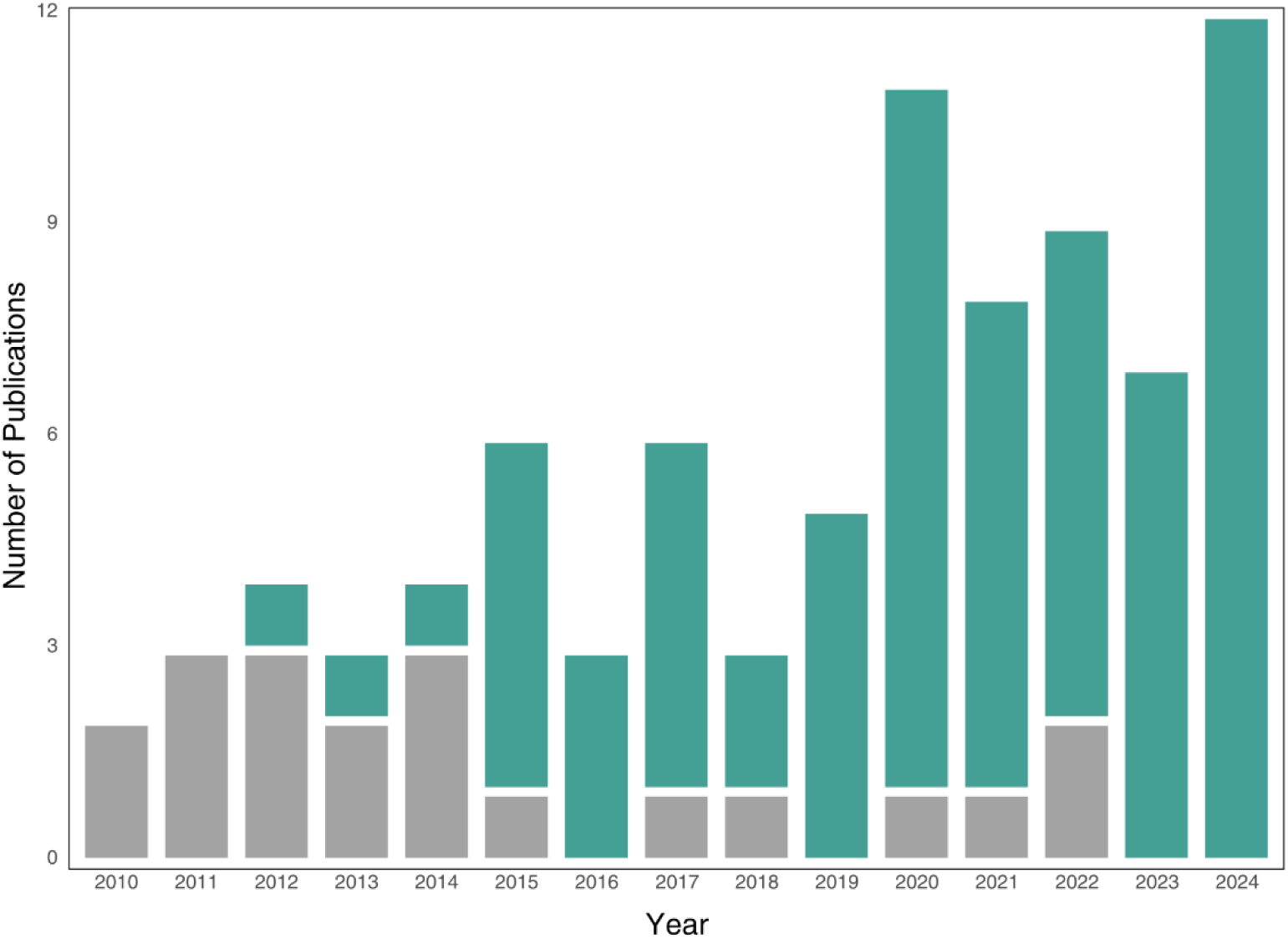
Temporal trend in final included publications (blue bars), identifying critical chondrichthyan areas in the European Atlantic and studies excluded solely due to non-contemporary data (grey bars) from 2010 – 2024.

#### Research by taxonomic group

Overall, 20 of the 27 focal species were studied in the final included studies (Figure 5). Seven species (*Chimaera monstrosa, Harriotta raleighana, Hydrolagus affinis*, *Hydrolagus mirabilis, Hydrolagus pallidus*, *Rhinochimaera atlantica* and *Scyliorhinus stellaris*) were entirely unrepresented and should therefore be prioritised for future research on critical areas. Conversely, *C. maximus* and *L. nasus* were the most studied species, together attracting over 20 % of the total research effort. The order Chimaeriformes appears to have been systematically overlooked; despite constituting 22 % of the focal species included in this study, they were not represented at all in the research effort. This likely reflects their comparative inaccessibility as deepwater fishes and aligns with the wider trend whereby species residing in deeper waters (e.g. *Leucoraja circularis, Leucoraja naevus, A. vulpinus*) were typically less studied than inshore species (e.g. *Raja montagui, Raja brachyura, Mustelus asterias).* Resultantly, deepwater critical habitat use remains a blind spot, consistent with findings from the Central and South American Pacific (García-Rodríguez *et al*., 2025). Generally, well-studied species were either severely threatened according to IUCN Red List (Jabado *et al*., 2024) or commercially important (e.g. *Raja clavata*), whilst those falling into neither category tended to be less studied (e.g. *Dasyatis pastinaca)*.

**Figure 5.**
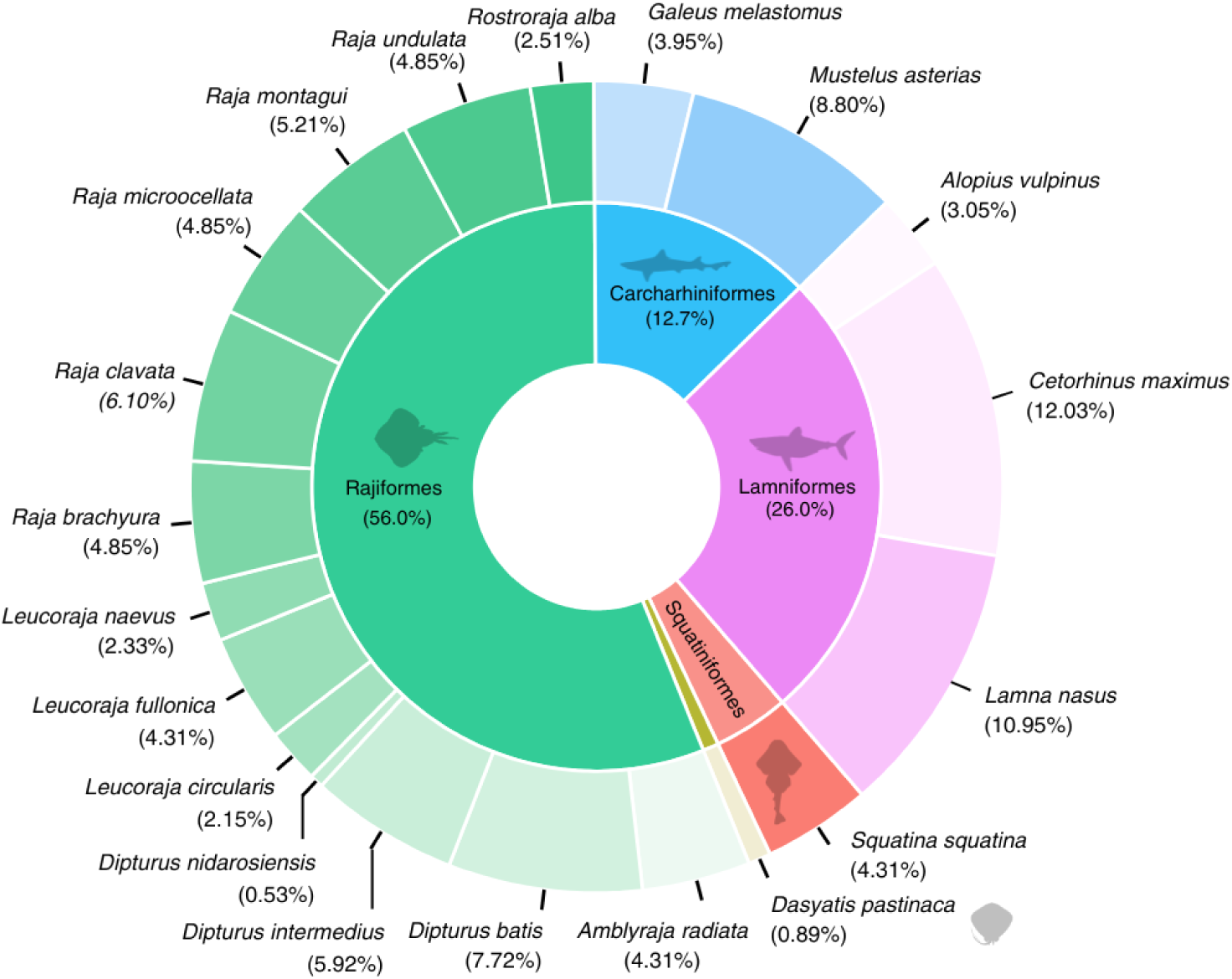
Frequency of species featured in the included research. Elasmobranch silhouettes are from PhyloPic (Keesey, 2023) and were selected based on general body form rather than species identity. Contributors: M. Kolhmann (2019; <u>CC0</u> <u>1.0</u>); I. Contreras (2022; <u>CC BY 3.0</u>); G. Dera (2023; <u>CC0 1.0)</u>.

#### Research by IUCN Red List category

The focal species selected for this study spanned five global IUCN Red List categories (Figure 6b). Despite representing a minority of focal species (n = 7, 25.9 %), the most threatened species [Critically Endangered (CR), and Endangered (EN)] were disproportionately represented in the selected literature (Figure 6a). Meanwhile, Vulnerable (VU) and Least Concern (LC) species were under-represented. Research effort was also uneven within IUCN categories. For instance, within the EN category, *C. maximus* and *L. nasus* attracted a disproportionate amount of research relative to other species such as *D. nidarosiensis* and *Rostroraja alba* (Figure 5).

**Figure 6.**
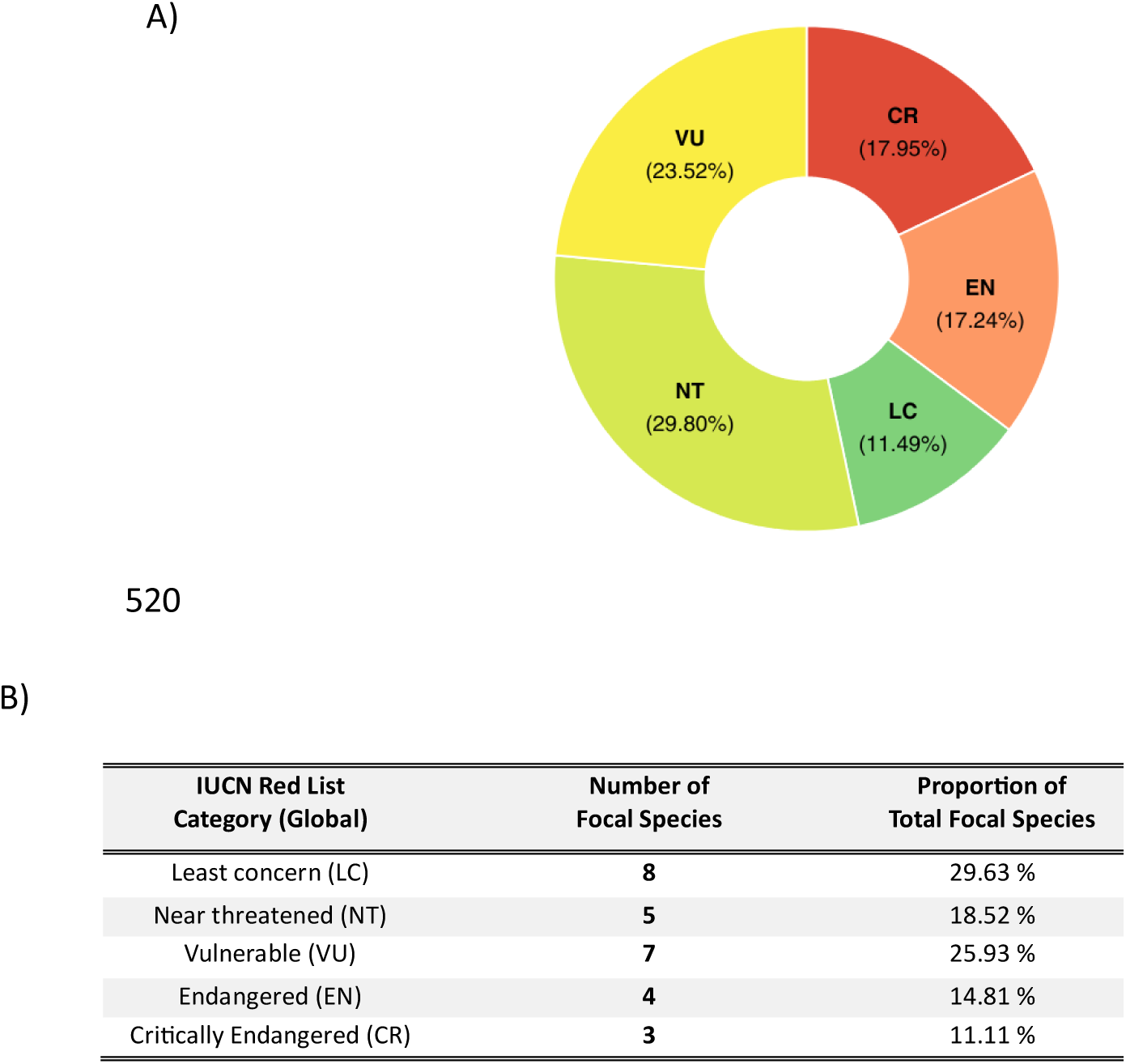
A) Quantity of papers examining focal species in each IUCN Red List category [global-level assessment (IUCN, 2025; See Table S1 for species-specific assessments used)], B) the number of focal species selected for this study in each IUCN red list category.

#### Geographic distribution

Within FAO division 27 (Northeast Atlantic), the included articles covered 36 ICES subareas, whereas in FAO division 34 (Eastern Central Atlantic), they spanned just three CECAF subareas (1.1, 1.2, and 2). Selected research was unevenly distributed, with hotspots in Portuguese inshore waters and around the UK and East of Ireland (Figure 7). ICES subarea 6.a (West of Scotland) was the most represented, appearing in 26 studies, perhaps partly reflecting a focus on UK priority species. Notably, 38.5 % (10 studies) of the research effort in 6.a was focused solely on *C. maximus.* Overall, offshore areas received less attention than coastal waters; for example, parts of the Oceanic Northeast Atlantic (ICES subareas 12.a, 12.c, and 10.b) were absent from all publications, likely due to their inaccessibility. Some regions only partially overlapped with the European Atlantic boundary (e.g. 5.b; 4.a), therefore, actual research effort is likely underestimated in these areas.

**Figure 7.**
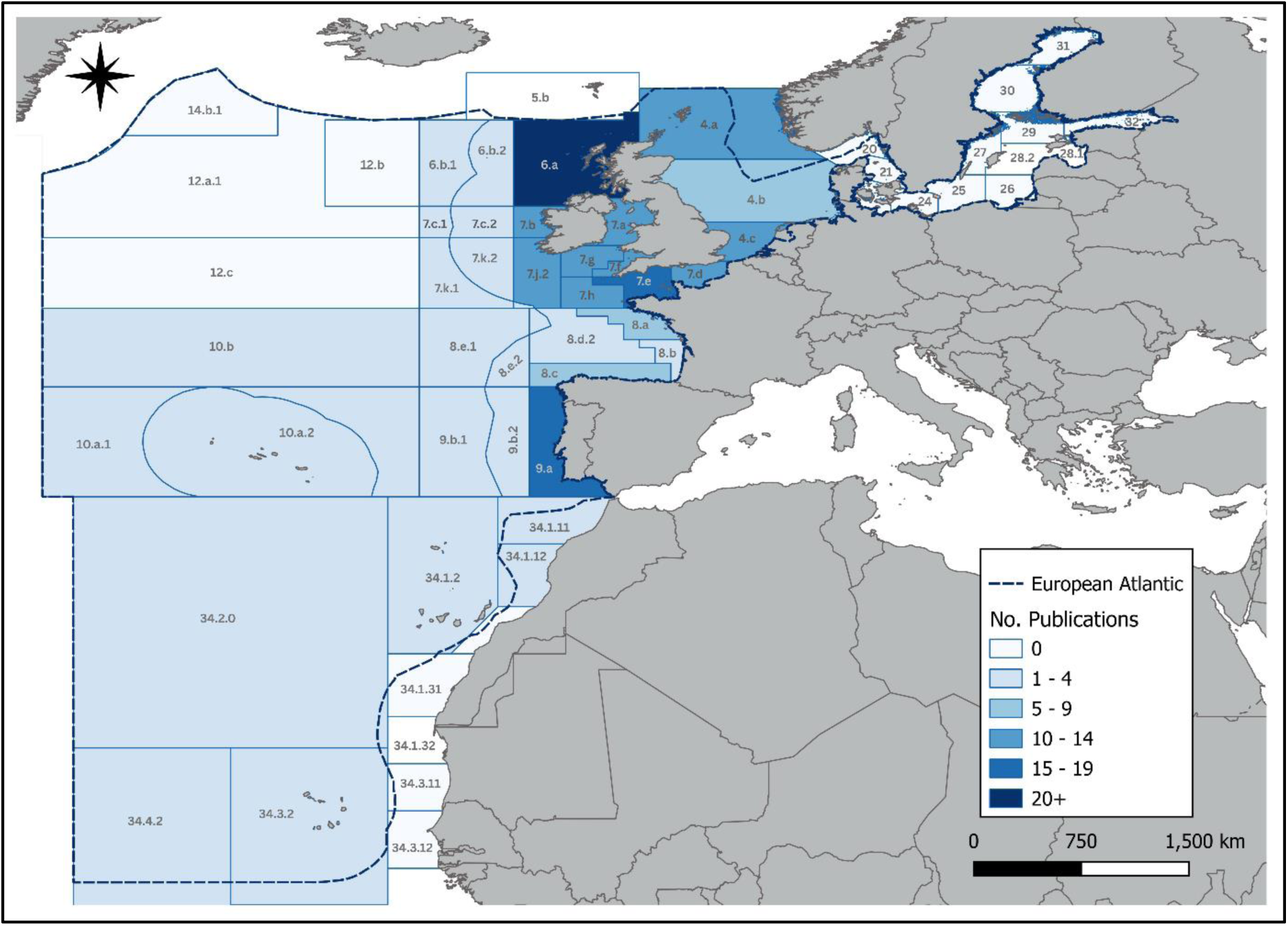
Spatial distribution of peer-reviewed research effort concerning critical chondrichthyan habitats in the European Atlantic, for UK priority species. Shading indicates the number of publications that sampled each ICES subarea division (FAO Area 27) or CECAF statistical division (FAO Area 34). Shapefiles for ICES Statistical Areas and CECAF Areas were sourced from the ICES Metadata Catalogue (ICES, 2005) and the FAO GeoNetwork (FAO, 2024) respectively. The European Atlantic boundary was sourced from the IUCN SSC Shark Specialist Group (IUCN SSC Shark Specialist Group, 2022).

### 4.2 Critical habitat evidence in the European Atlantic

The assessment of all extracted critical area evidence, according to the requirements met and associated quality assessment (see Table S8 for a detailed table of the requirements met and risk of bias per area), demonstrated taxonomic and critical area category biases (Figure 8). *Raja clavata* and *S. squatina* were most documented, whilst *D. nidarosiensis* was scarcely represented. Overall, most evidence was rated *Weak/ Cannot Assess* (< 1.5 requirement score), with few *Strong* (> 1.5 requirement score) records, even for well-studied species, indicating a need for targeted research into these areas. Further, where several *Strong* records were present for a species, these were not equally distributed among critical area types; for instance, *S. squatina* had the most *Strong* records, but these were all immature areas. In addition, more research effort (Figure 5) does not necessarily translate into the identification of more *Strong* and *Moderate* areas (e.g. *R. montagui* vs *C. maximus* (Figure 8). In terms of critical areas, immature areas attracted the most research, which translated into the most *Strong* evidence areas. Whereas, no *Strong* evidence resting areas were identified. This uneven research distribution signifies a need for a more systematic approach to critical area research.

**Figure 8.**
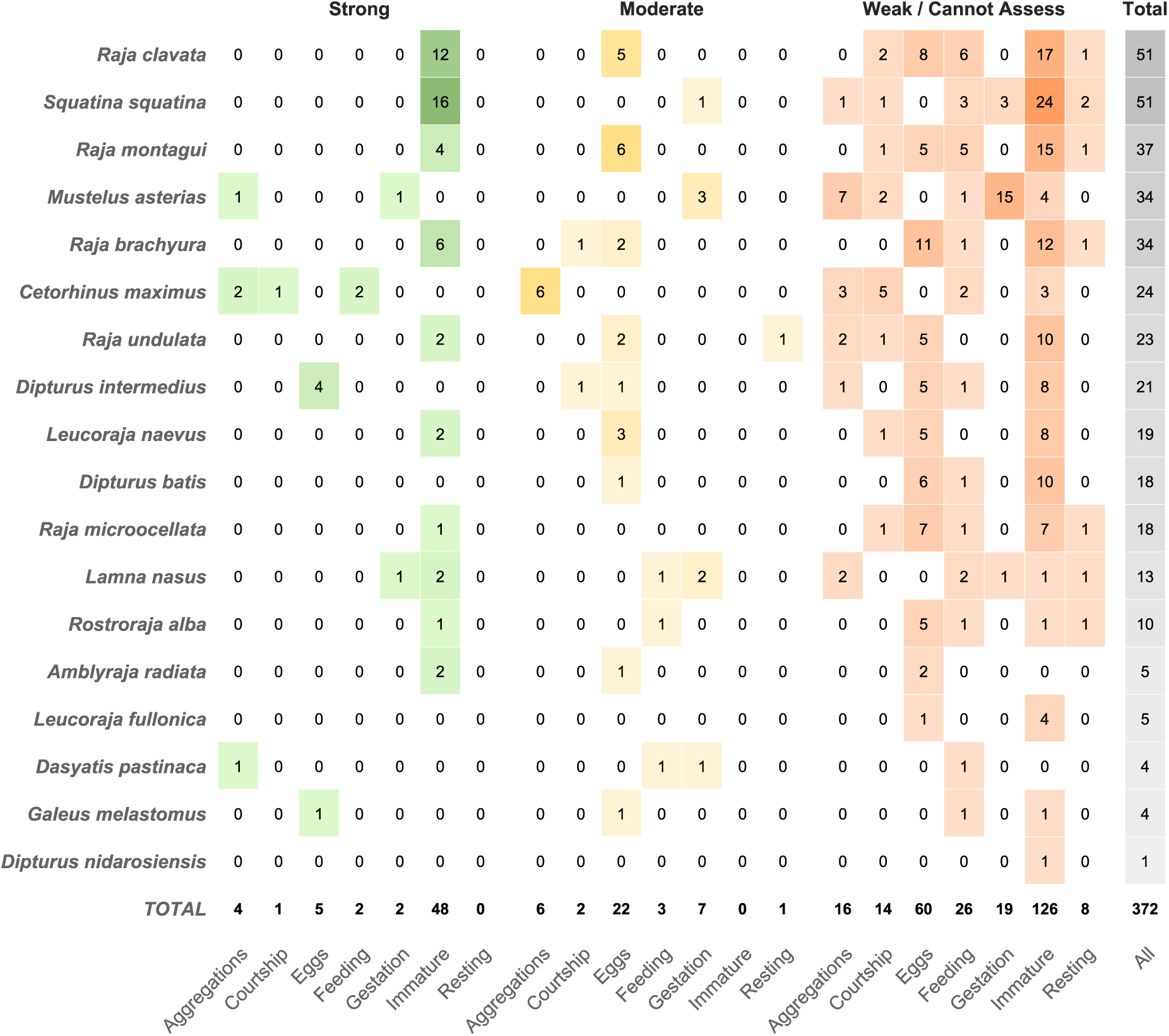
Matrix demonstrating the distribution and strength of evidence for critical area categories per species. The shade and colour of tiles indicates the number of areas identified. The values are based on the number of areas extracted (Table S8) prior-to processing for mapping.

Immature areas comprised over three-quarters (77 %) of all *Strong* critical areas identified and combined with egg-laying areas constituted 78 % of all *Strong* and *Moderate* reproductive habitats studied. This likely reflects the conceptual clarity of nursery and egg-nursery habitats, facilitating their study (Heupel *et al*., 2007, 2019; Martins *et al*., 2018). A decade after the shark nursery paradigm was proposed, Heupel *et al*. (2019) found the concept had been widely adopted (398 citations as of June 2018), representing growing interest in the study of early life stage chondrichthyan habitats. This review also shows a post-2019 rise in immature area studies (Figure S3), likely influenced by further conceptual developments (Martins *et al*., 2018; Heupel *et al*., 2019) or increased interest galvanised by the ISRA process (Hyde *et al*., 2022). Although Heupel *et al*. (2019) noted limited research effort on temperate species, this review found a substantial increase in studies after 2019 (Figure 4), in the European Atlantic, a predominantly temperate and warm-temperate region.

In contrast, habitats used exclusively by mature chondrichthyans for reproductive purposes (mating and gestation/parturition areas) were understudied, representing just 12 % of *Strong* and *Moderate* evidence critical areas. This gap is significant because protecting mature adults, particularly pregnant females, is widely supported for conserving slow-growing fishes, by helping maintain high population recruitment (Simpfendorfer, 1999; Prince, 2005; Wiegand, Hunter & Dulvy, 2011). Expanding research on reproductive habitats used by mature individuals is therefore a conservation priority.

#### Aggregations

Evidence for aggregations was found in 13 studies, identifying 26 species-specific areas (Figures 8–10). Four areas demonstrated *Strong* aggregation area evidence (≥ 2 requirements met), whilst six exhibited *Moderate* evidence [1.5 requirements met (see Table S8 for a species-specific area breakdown)]. 80 % of these were for *C. maximus* around Scotland and Ireland, highlighting the Northwest British Isles as a key region for this species.

**Figure 9.**
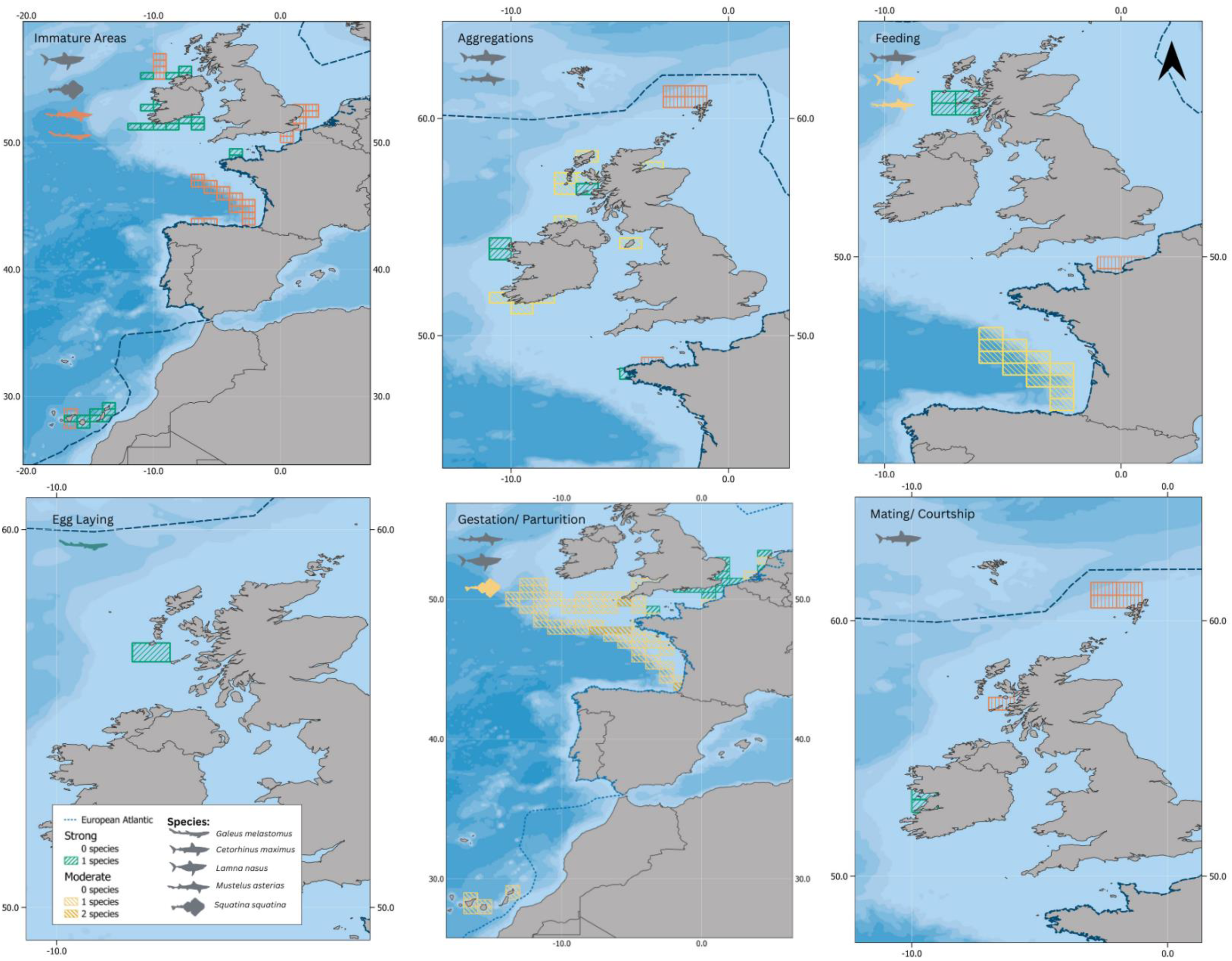
The distribution of potential critical areas in the European Atlantic for shark species by ICES rectangle meeting one or more requirements. Rectangle colours represent the strength of evidence, based on the number of requirements met (1 = Weak, 1.5 = Moderate, > 1.5 = Strong). Shark icon colours indicate the evidence strength associated with each species, with grey icons representing more than one strength categories. Silhouettes are from PhyloPic (Keesey, 2023) and were selected based on general body form rather than species identity. Contributors: I. Contreras (2022; <u>CC BY 3.0</u>); G. Dera (2023; <u>CC0 1.0)</u>; S. O’Connor (2023;<u>CC BY 4.0</u>); B. Lang. (2016; <u>CC0 1.0)</u>.

**Figure 10.**
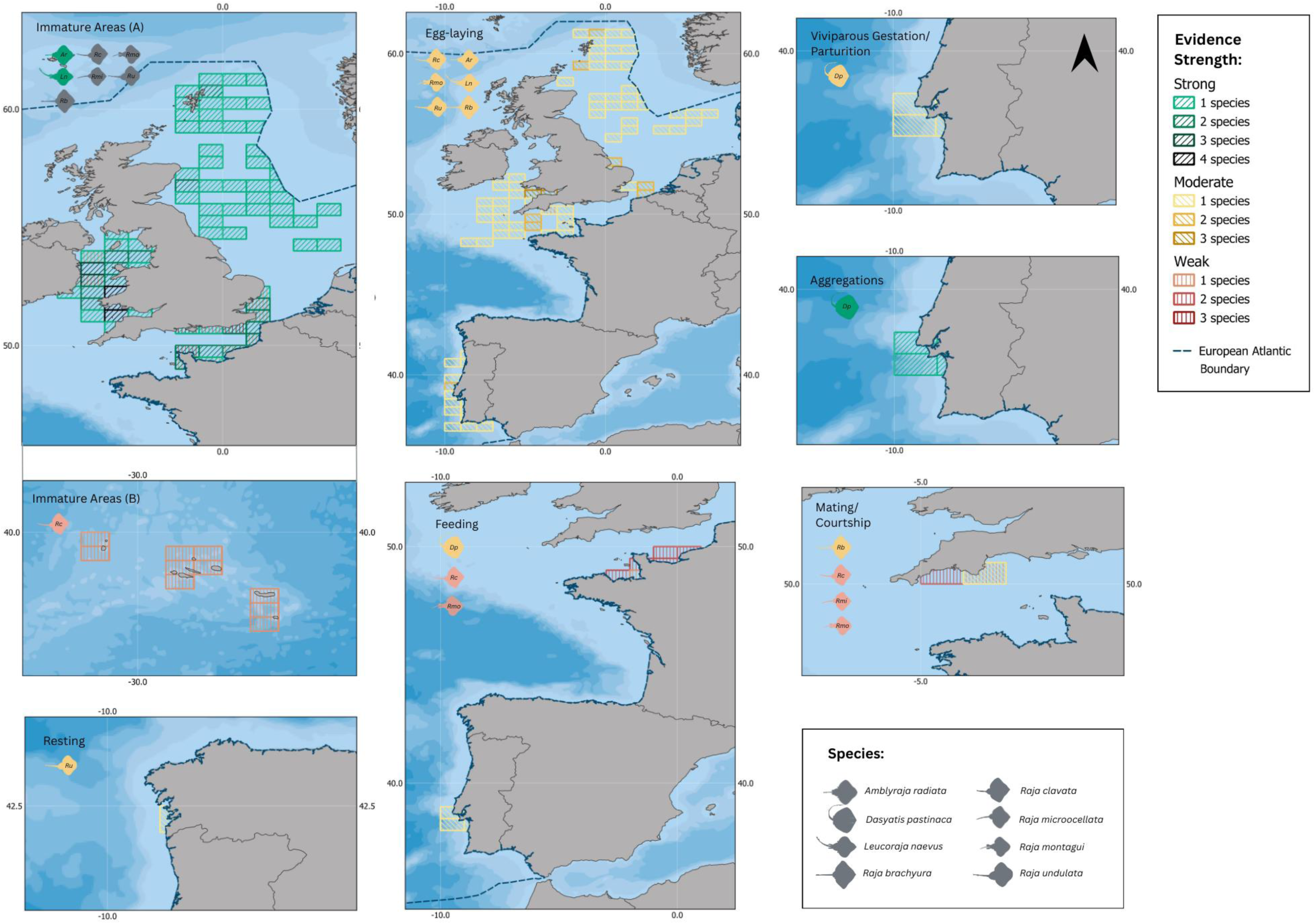
Distribution of all potential critical areas in the European Atlantic for small [< 2 m total length (TL)] batoid species by ICES rectangle meeting one or more requirements. Rectangle colours represent the strength of evidence, based on the number of requirements met (1 = Weak, 1.5 = Moderate, > 1.5 = Strong). Silhouettes are from PhyloPic (Keesey, 2023) and were selected based on general body form rather than species identity. Contributors: M. Kolhmann (2019; <u>CC0 1.0</u>); I. Contreras (2022; <u>CC BY 3.0</u>); N. Hermann (2021 & 2023; <u>CC0 1.0</u>); I. Braasch (2022; <u>CC0 1.0</u>); M. Pélissié (2022; <u>CC0 1.0</u>).

Definitions of aggregation varied: six studies omitted a clear definition, reinforcing the need for a unified paradigm (McInturf *et al*., 2023). Six studies applied a lower bound of two individuals (Gore *et al*., 2019; Kraft *et al*., 2024b; Thorburn *et al*., 2024) or implied this threshold using “multiple individuals together” (Lieber *et al*., 2020; Biton-Porsmoguer & Lloret, 2023), while one required groups of “more than two” (Meyers *et al*., 2017), aligning with stricter evidence standards for spatial management, such as those applied for the ISRA process (Hyde *et al*., 2022). One study described multispecies assemblages as “preferential aggregation sites,” applying aggregation to signal elevated diversity rather than density (Serra-Pereira *et al*., 2014a).

The small sample of literature presented here supports using ≥ two individuals as a provisional definition of an aggregation (McInturf et al., 2023). However, since pairs may co-occur by chance, additional criteria are required to strengthen inference: evaluating shared social, environmental, or biological drivers; assessing temporal persistence; and clearly specifying spatiotemporal windows of co-occurrence (McInturf *et al*., 2023). These safeguards distinguish true aggregations from general high-use or high-density areas, in which individuals are not required to exhibit close and deliberate spatiotemporal proximity. By contrast, once a driver is identified, the ISRA framework reclassifies aggregation areas according to said driver [e.g., a feeding aggregation becomes a feeding area (Hyde *et al*., 2022)].

Temporal windows were rarely specified; when reported, they were typically ≤ 24 hours (Biton-Porsmoguer & Lloret, 2023; Thorburn *et al*., 2024) but occasionally shorter [15 minutes (Kraft *et al*., 2024b)]. Spatial windows varied by method: acoustic telemetry studies used co-detections at the same receiver, corresponding to a ≈ 200–600 m co-detection sphere depending on tag type and environmental conditions (Carlson et al., 2023; Kraft et al., 2024b; Thorburn et al., 2024), visual surveys typically used ≈ 100 m proximity (Gore et al., 2019), and the recreational angling study defined aggregations as sharks caught on the same beach on the same day (Biton-Porsmoguer & Lloret, 2023).

Greater consistency in definitions, transparency in spatiotemporal thresholds used and focus on underlying drivers would improve comparability and strengthen identification of genuine aggregations. We recommend defining provisional aggregations as ≥ two individuals with clearly specified spatiotemporal criteria (McInturf *et al*., 2023). When management or resource-use decisions require stronger evidence, a stricter threshold of ≥ three individuals may be preferable, particularly if the underlying driver is unknown (Hyde *et al*., 2022).

#### Resting

Resting was the least-studied area type, with evidence from six studies covering nine areas and eight species (Figures 8–10). Evidence was not restricted to benthic species; for example, the Bay of Biscay was proposed as a potential resting area for *L. nasus*, though empirical support was limited (Carrasco-Puig *et al*., 2024). Most evidence was *Weak/Cannot Assess*, failing to meet criteria for persistent habitat use (R2–R3; Table 1). Moderate evidence was reported for *R. undulata* in Ría de Vigo, Spain, where acoustic telemetry indicated lower daytime activity and increased site occupancy, confirming preferred resting use (Leeb *et al*., 2021). When activity levels were examined, elevated nocturnal or crepuscular activity was observed in several species [*S. squatina*, *R. undulata*, *R. alba* (Sousa *et al*., 2019; Tuya, Asensio & Navarro, 2020; Leeb *et al*., 2021; Mead *et al*., 2023)].

The limited resting evidence likely reflects both methodological challenges and physiological constraints. Obligate ram-ventilators (e.g., *C. maximus*, *L. nasus*, *A. vulpinus*) must swim continuously, so rest in these species may be cryptic, expressed as slower swimming or reduced distances travelled rather than complete inactivity (Kelly *et al*., 2019). By contrast, species capable of buccal pumping or drawing water over their gills via spiracles can rest while stationary (Kelly *et al*., 2020, 2021).

Consequently, both the detectability of critical resting areas and their suitability for management likely vary with ventilation strategy. Circadian variability, both between and within species, further complicates identification. For example, in Portugal, female *R. alba* exhibited clear crepuscular activity and diurnal rest phases, whereas males alternated between states more frequently, indicative of cathemeral (irregular diurnal and nocturnal) activity (Sousa *et al*., 2019). Only a small proportion (11 %) of focal species are confirmed obligate ram-ventilators, suggesting most are capable of rest. Thus, the scarcity of identified potential resting areas more likely reflects a research gap. Resting elasmobranchs are considered particularly vulnerable to towed fishing gears (Thorburn *et al*., 2022), yet their limited mobility makes them well-suited to spatial management. Simple gear modifications (e.g. removal of tickler chains from trawls) are also available to reduce mortality (McIntyre *et al*., 2015; Kynoch, Fryer & Neat, 2015), therefore, these areas represent a key knowledge and management opportunity gap.

#### Feeding Areas

Evidence for feeding areas was found in 21 studies, relating to 31 species-specific areas and 13 species (Figures 8–11). This translated into two *Strong* and three *Moderate* strength evidence areas (see S7 for further detail). All remaining evidence scored *Weak/ Cannot Assess*, with dietary studies examining stomach content featuring heavily (13 areas). For a feeding area to be considered critical, relative area use for feeding (R2-R3) and/ or relative feeding rates (R1) must be higher than other areas. Although not primarily focused on critical areas, some dietary studies provided relevant data for this purpose [e.g. repeated use of the Celtic Sea over multiple years by *D. batis* (Brown-Vuillemin *et al*., 2020)] by reporting sample timings and locations. Some also included the vacuity index [proportion of empty stomachs (Biton-Porsmoguer, 2022)], which, when presented by area, represents a straightforward metric to compare relative feeding rates (standardised by effort, R1) and thus increase the relevance of dietary studies to critical feeding area identification.

**Figure 11.**
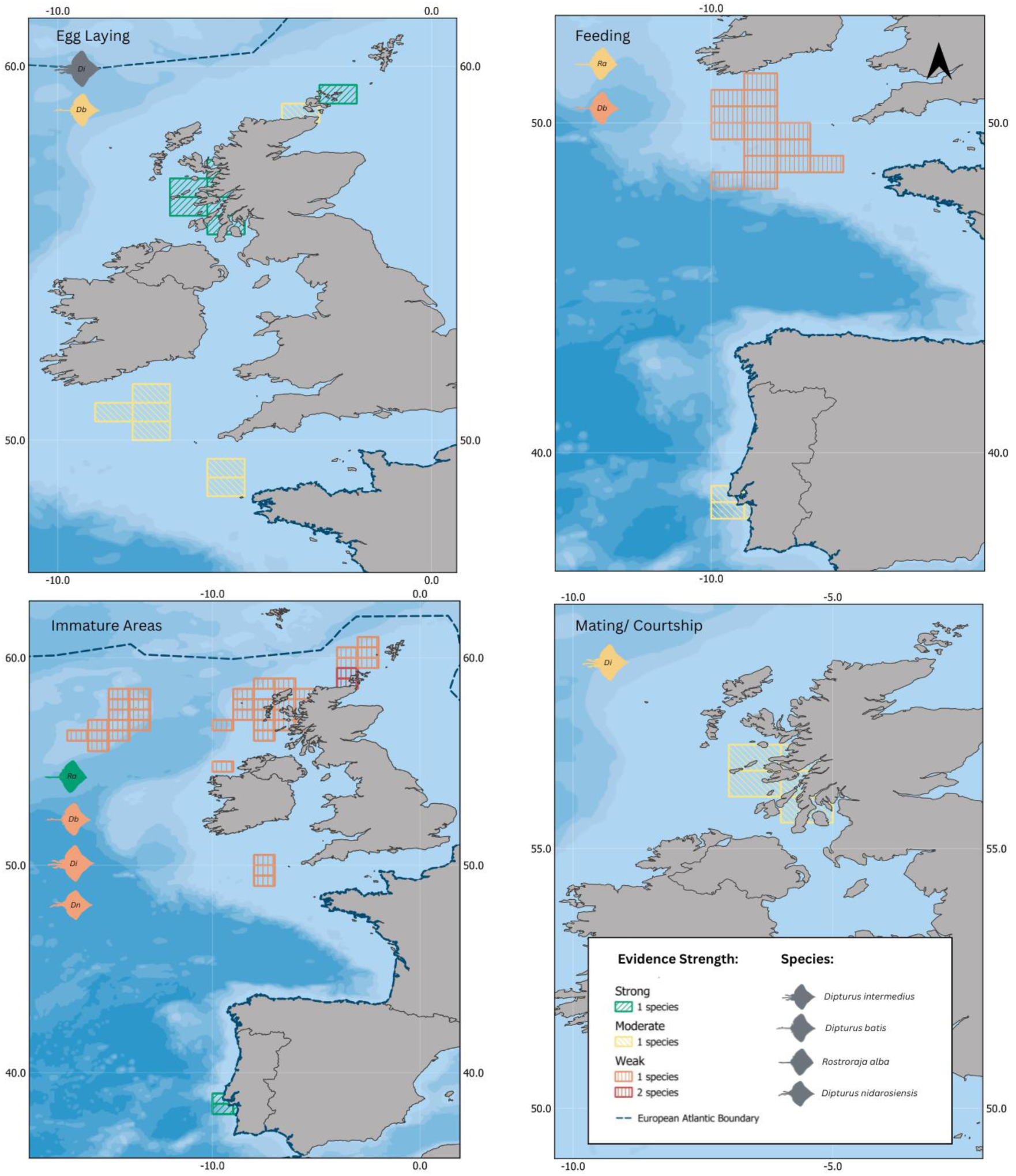
Distribution of potential critical areas in the European Atlantic for large (> 2 m total length) batoid species* by ICES rectangle meeting 1 or more requirements. Rectangle colours represent the strength of evidence, based on the number of requirements met (1 = Weak, 1.5 = Moderate, > 1.5 = Strong). Silhouettes are from PhyloPic (Keesey, 2023) and were selected based on general body form rather than species identity. Contributors: I. Contreras (2022; <u>CC BY 3.0</u>); G. Dera (2023; <u>CC0 1.0</u>); T. Rehill (2023; <u>CC0 1.0</u>).

#### Mating/ Courtship

Mating and courtship evidence was identified in 13 studies, spanning 17 species-specific areas and 10 species (Figures 8–11). Evidence was *Strong* for one area and *Moderate* (downgraded due to lack of direct mating observations) for two (see table S8 for species-specific area breakdown). *Strong* evidence was found for *C. maximus* courtship off County Clare and County Galway, Ireland, where large circling groups (toruses) were observed over multiple days (R2; max. 5 days in 2021) and across three years (Sims *et al*., 2022). Multiple indicators confirmed the courtship function: mixed sexes, mating scars (abrasions on claspers and pectoral fins), exclusivity to mature individuals and absence of feeding (Sims *et al*., 2022).

Spatial overlap of sexes can be used to examine potential mating areas (Simpson *et al*., 2021) and is more compellingly linked to mating when supported by maturity or active reproductive status data. For example, in the Loch Sunart to Sound of Jura MPA (Scotland), seasonal (March–April) site fidelity of *D. intermedius* was attributed to potential mating, supported by mature male-female overlap and later confirmation of female reproductive activity through hormonal change-point analysis (Lavender *et al*., 2022; Thorburn *et al*., 2023). Breaching was also identified as a potential mating behaviour for *C. maximus* (Hayes *et al*., 2018), however, there remains some ambiguity around the purpose of this behaviour (Gore *et al*., 2019; Sims *et al*., 2022; Berrow *et al*., 2025).

Overall evidence strength for mating areas was often limited by a lack of direct observational data to confirm mating (Hayes *et al*., 2018; Simpson, Humphries & Sims, 2021). Traditional approaches, such as fisheries specimen examinations, can provide useful insights into reproductive activity (Maia *et al*., 2012; Serra-Pereira, Erzini & Figueiredo, 2015; McCully Phillips & Ellis, 2015; Santos *et al*., 2021). In-water observation and video can also yield fortuitous mating observations (Vossgaetter *et al*., 2025), meanwhile angler-led citizen science databases and fishers’ experiential knowledge can offer further insight, as elasmobranchs often exhibit recent signs of mating [e.g., swollen claspers, mating scars (Mawer *et al*., 2025)] and are occasionally directly observed in copulatory embrace during capture (M. Larkin, recreational angler, pers. comm.). More detailed insights could be gained through wider application of non-invasive methods, such as using ultrasonography to examine reproductive systems (Whittamore *et al*., 2010; Thorburn *et al*., 2023).

#### Viviparous Gestation and Parturition

Evidence for gestation and/or parturition areas for viviparous species was found in only six studies, covering 28 species-specific areas, for four of the six focal viviparous species: *L. nasus, D. pastinaca, M. asterias* and *S. squatina* (Figures 8–9). Of these, two areas were classified as *Strong* and four as *Moderate* (Table S8). The strong evidence gestation areas were for *L. nasus* in the Tregor Area, France (Jung *et al*., 2024a) and the Eastern English Channel/ Southern North Sea for *M. asterias* (McCully Phillips & Ellis, 2015; Silva & Ellis, 2019; Griffiths *et al*., 2020; Brevé *et al*., 2020). Eleven other areas met a single requirement, signifying potential importance (Figure 9).

In general, gestation evidence was difficult to entangle from parturition evidence, and indeed, there was evidence for colocation of these two reproductive behaviours (e.g. Tregor Area, Bay of Biscay, Celtic Sea: *L. nasus*; Western English Channel: *M. asterias* (McCully Phillips & Ellis, 2015; Biais *et al*., 2017; Jung *et al*., 2024a)). Use of novel technologies such as birth-alert tags (Sulikowski & Hammerschlag, 2023) may reveal fine-scale habitat-partitioning between these two areas, resulting from differences in resource and refuge requirements between neonates and pregnant females. Most evidence for gestation/parturition areas was found in the warmer parts of species’ ranges, with all areas being located below 53°N. This aligns with the common use of warmer waters by pregnant elasmobranchs, to speed up metabolic rate and resultant embryonic development, as seen in leopard sharks [*Triakis semifasciata* (Hight & Lowe, 2007)]. Evidence for gestation areas was also concentrated in productive, nutrient-rich waters, such as coastal shelf and island upwelling regions around the UK and Canary Islands, perhaps reflecting the intensive energy requirements for both embryonic development and growth-to-maturity. However, this trend may also be an artefact of uneven research effort, which was concentrated in coastal waters, and therefore requires further investigation.

#### Immature Areas (Free-swimming Stages)

Evidence for immature areas, was found in 26 studies, resulting in 174 species-specific areas (Figures 8–11). Forty-eight areas were classified as *Strong* for 10 species, and all other areas were *Weak/ Cannot Assess* (Table S8). Most *Strong* evidence originated from a small number of studies (e.g., Jiménez-Alvarado *et al*., 2020; Ellis *et al*., 2024) signalling a need for broader research efforts across species and regions. Not all chondrichthyan species are expected to exhibit spatially explicit immature areas (Beck *et al*., 2001; Heupel *et al*., 2007), which may explain why evidence was lacking or limited for several species (Figure 8). Potential areas of multispecies importance were evident both from primary data in Portugal and the Canary Islands (Serra-Pereira *et al*., 2014; Tuya *et al*., 2020) and by synthesising species-specific areas [e.g. Bristol Channel, the Irish Sea and eastern English Channel for several skate species (Figure 10)].

Shark nursery literature increasingly recognises the limitations of focusing solely on neonate and YOY habitats, as later juvenile and subadult stages may contribute equally, or more, to population recruitment, particularly in slow-growing, late-maturing species with ontogenetic habitat partitioning (Heppell, Crowder & Menzel, 1999; Simpfendorfer, 1999; Gallucci, Taylor & Erzini, 2006; Kinney & Simpfendorfer, 2009). Earlier frameworks encompassed these intermediate stages by dividing areas into *primary* and *secondary* nursery areas (Simpfendorfer & Milward, 1993), but the stricter early-juvenile-focused definition in Heupel *et al*. (2007) has since become dominant, streamlining the criteria and reducing potential confusion. This shift appears to have narrowed conservation planning frameworks towards the earliest juveniles whilst overlooking older juvenile stages (Hyde *et al*., 2022).

To address this gap, this review examined all potentially critical immature areas, including both potential: critical juvenile areas and critical early juvenile areas (Figures 2–3; Table S6). This approach identified 23 *Strong*-evidence early juvenile areas for six species, where sites with neonate, YOY, or first-cohort observations met > 1.5 requirements [R1 – R3 (Cameron et al., 2019; Ellis et al., 2024a)]. Areas in this category represent potential Nursery Areas (*sensu* Figure 3) since the requirements (R1 – R3) were based on the criteria in Heupel *et al*. (2007); however, since no areas met all three criteria, further study is required to confirm their status. One study concluded that Playa del Castillo for *S. squatina* qualified as a Nursery Area (*sensu* Figure 3), however, the size threshold used included juveniles larger than YOY and the lack of effort-standardised abundance data raised uncertainty over whether the first criterion from Heupel *et al*. (2007) was satisfied, thus, further investigation is warranted. Meanwhile, 24 areas exhibited *Strong* evidence for potential Critical Juvenile Areas [*sensu* Figure 3 (Jiménez-Alvarado *et al*., 2020; Ellis *et al*., 2024; Kraft, Winkler & Abecasis, 2024a; Jung *et al*., 2024b)] where size thresholds included Juveniles [> size at one year (Figure 2)]. Integrating species-specific population dynamics to identify the size classes most important for population recruitment is essential for guiding management decisions, ensuring that critical immature areas are prioritized based on their contribution to population sustainability (Kinney & Simpfendorfer, 2009).

The broader nursery-role hypothesis literature emphasises quantifying the functional contribution of recruits to adult populations through metrics such as growth, survival, and juvenile-to-adult recruitment (Beck *et al*., 2001). This review found limited engagement with such metrics: growth-to-maturity evidence was only reported for *D. intermedius* and *L. nasus* from mark-recapture studies (Cameron *et al*., 2019; Thorburn *et al*., 2023), and was not directly linked findings to immature habitat function. This gap likely reflects both the comparatively limited research effort on chondrichthyan early life habitats (9 %) relative to other marine taxa [e.g. 76 % for actinopterygians (Ciotti *et al*., 2025)] and the logistical challenges of quantifying such metrics for rare or slow-growing species, which led to their exclusion from the formal shark nursery concept (Heupel *et al*., 2007). Heupel *et al*. (2007) proposed that areas with high juvenile density relative to surrounding habitats, and consistent use, could reasonably be assumed to contribute disproportionately to adult populations. However, few areas reviewed here satisfied both criteria, indicating that evidence for high relative recruitment, even when inferred indirectly, remains limited. Direct measurement through archival tissue sampling is challenging for chondrichthyans, since poor calcification of hard tissues (i.e. vertebrae) often limits reliability for age determination (Matta *et al*., 2017) meanwhile lethal sampling to access such tissues is less justifiable for threatened, slow-growing species (Hammerschlag & Sulikowski, 2011). Further development and validation of non-lethal archival tissue sampling (e.g. caudal thorns and spines) for growth indices could fill this methodological gap (McMillan *et al*., 2017). Similarly, the larger size and lower mortality in early life stages makes artificial tagging another viable option for directly measuring growth and recruitment in chondrichthyans.

Overall, since the strongest evidence of functional habitat value will ultimately come from direct measures of growth, survival, and juvenile-to-adult recruitment, such metrics should be pursued where feasible. However, the critical area definition proposed here (Table 1) demonstrates a practical, framework, suited to current evidence limitations, whilst enabling a more inclusive lens to examine all potentially critical immature areas, not just those for the earliest life stages.

#### Egg-laying Areas

Egg-laying areas were comparatively well studied, with 16 publications covering 12 species (Figures 8; 10–11). Evidence was *Strong* for five areas and *Moderate* for 22 (Table S8). Most *Strong* evidence came from Scottish waters for *D. intermedius* and *G. melastomus* (Phillips *et al*., 2021; Thorburn *et al*., 2021a, 2023; Dodd *et al*., 2022; Schwanck *et al*., 2024) indicating regional significance. The only *Strong*-evidence shark example was the Mingulay Reef, where *G. melastomus* deposited egg cases within *Lophelia pertusa* reefs under highly specific environmental conditions (Henry *et al*., 2013). Interestingly, this area was not captured by the European Atlantic ISRA process. As with nursery areas, substantial evidence was concentrated in a small number of studies (e.g., Ellis *et al*., 2024), however, sites were typically limited to *Moderate* evidence due to lack of interannual occurrence data.

This review aimed to identify all potential Critical Egg-laying Areas (*sensu* Figure 3) of possible value to spatial management decisions. The approach adopted, using the critical area definition (R1–R3), differed from Martins *et al*. (2018) in that broader evidence types could be considered to signal potential importance. In addition to in-situ eggs (attached to substrate), stranded egg cases and egg-bearing and mature female habitat use were also considered. Meanwhile, the greater density criterion (R1) could be more generally satisfied by a greater density/ frequency of occurrence of any of these evidence types, rather than exclusively by in-situ eggs. However, *Strong* evidence was limited to egg-bearing females and in-situ eggs. Therefore, areas potentially satisfying the criteria for Egg Nurseries [*sensu* Figure 3 (Hoff, 2016; Martins *et al*., 2018)] can be found in the *Strong* category.

The Red Rocks and Longay MPA scored highest and satisfied more formal egg nursery criteria than any other site (Dodd *et al*., 2022). Because the Hoff (2016) and Martins *et al*. (2018) egg nursery frameworks are largely overlapping, with Hoff’s requirement that eggs contact the benthos effectively embedded within Martins’ first criterion, both yielded similar assessments. The only unmet criterion across both frameworks was post-hatching habitat partitioning away from the oviposition site (Dodd *et al*., 2022), evidence for which was rarely presented in the included literature [n = 14 areas (Serra-Pereira *et al*., 2014; Ellis *et al*., 2024)]. This likely reflects the practical difficulty of proving absence, which necessitates extensive sampling. Where such segregation has been demonstrated [e.g., Loch Sunart to Sound of Jura MPA (Dodd *et al*., 2022)], evidence strongly supports Egg Nursery Area function (Martins *et al*., 2018). However, Critical Egg-laying Areas (*sensu* Figure 3) without confirmed segregation should not be discounted, as overlapping egg-laying and early juvenile areas may represent priority targets for conservation, since management could concurrently protect both vulnerable life stages, along with egg-bearing females during oviposition.

The requirement to support high egg densities is common to both key egg-nursery concepts (Hoff, 2016; Martins *et al*., 2018). While the SACFOR scale has occasionally been applied as a universal “high density” threshold (Phillips *et al*., 2021), this risks overlooking species-specific variation in egg-laying behaviour. In line with the conclusions of this review and Dodd *et al*. (2022), we instead recommend evaluating relative egg-laying density against species-specific baselines of typical egg densities. For species with widespread egg-laying, thresholds can be adjusted progressively as additional data become available, for example, prioritising areas that exceed the mean density observed across known egg-laying sites, to identify the most ecologically important habitats for conservation.

Finally, potentially critical communal egg-laying areas were identified in waters around the British Isles, where *Moderate* evidence for up to three species overlapped. These included The Wash, the Western English Channel, Bristol Channel, waters adjacent to the Shetland Islands and the Outer Thames and Belgian Coastal waters. All of these were found in coastal waters < 100 m (Figure 10), though this should be interpreted with consideration of uneven sampling effort (Figure 7). These sites are strong candidates for towed-gear restrictions, given the vulnerability of egg cases to physical disturbance (North Pacific Fishery Management Council, 2014).

### 4.3 Considerations and Future Directions

#### Addressing Conceptual and Methodological Biases in Critical Area Research

Recognising conceptual and methodological biases is essential for robustly identifying the most critical areas. Framing areas along two axes, completeness of evidence and strength of evidence (Figure 12), helps examine how such biases shape inferred levels of importance. Well-studied areas meeting evidence thresholds (e.g. R1 – R3) are classified as “known critical areas” (Q2) while “unknown areas” have no available data (Q1 & Q3; Figure 10), leaving their ecological importance entirely uncertain. Between these lie data-incomplete areas, where partial or inconclusive evidence prevents confident assessment.

**Figure 12.**
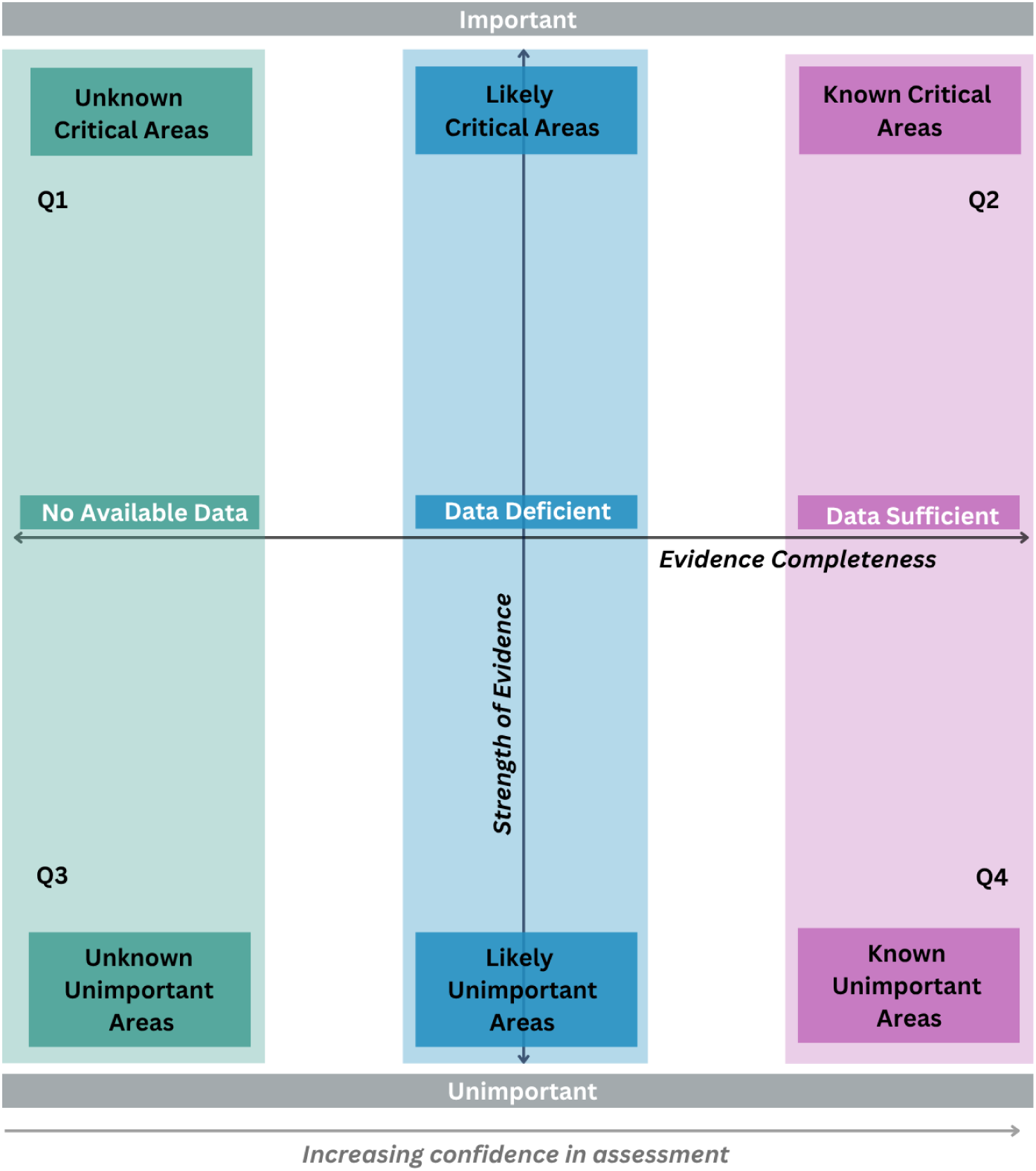
Conceptual framework organising critical area knowledge and blind spots along the two axes: evidence completeness and strength of evidence.

The uneven distribution of research effort introduces a positive-detection bias, whereby heavily studied regions are more likely to appear critical than under-sampled ones. This bias should diminish as taxonomic and spatial coverage expands, and this review highlights priority regions and taxa to target this effort (Figures 5;7). Yet, since the evidence base is currently limited (Figure 8), this bias should be mitigated where feasible. Requirement R1 has potential to reduce this bias by evaluating candidate critical area importance relative to other surveyed areas. However, R1 was rarely satisfied in the examined areas (Figure 13), limiting the confidence of many area assessments in this review. Broader, effort-standardised habitat-use studies (e.g., scientific trawl surveys) and establishing species-specific baselines of “normal” area use would provide the comparative context needed to distinguish genuinely high-use areas from those that merely appear important due to uneven sampling.

**Figure 13.**
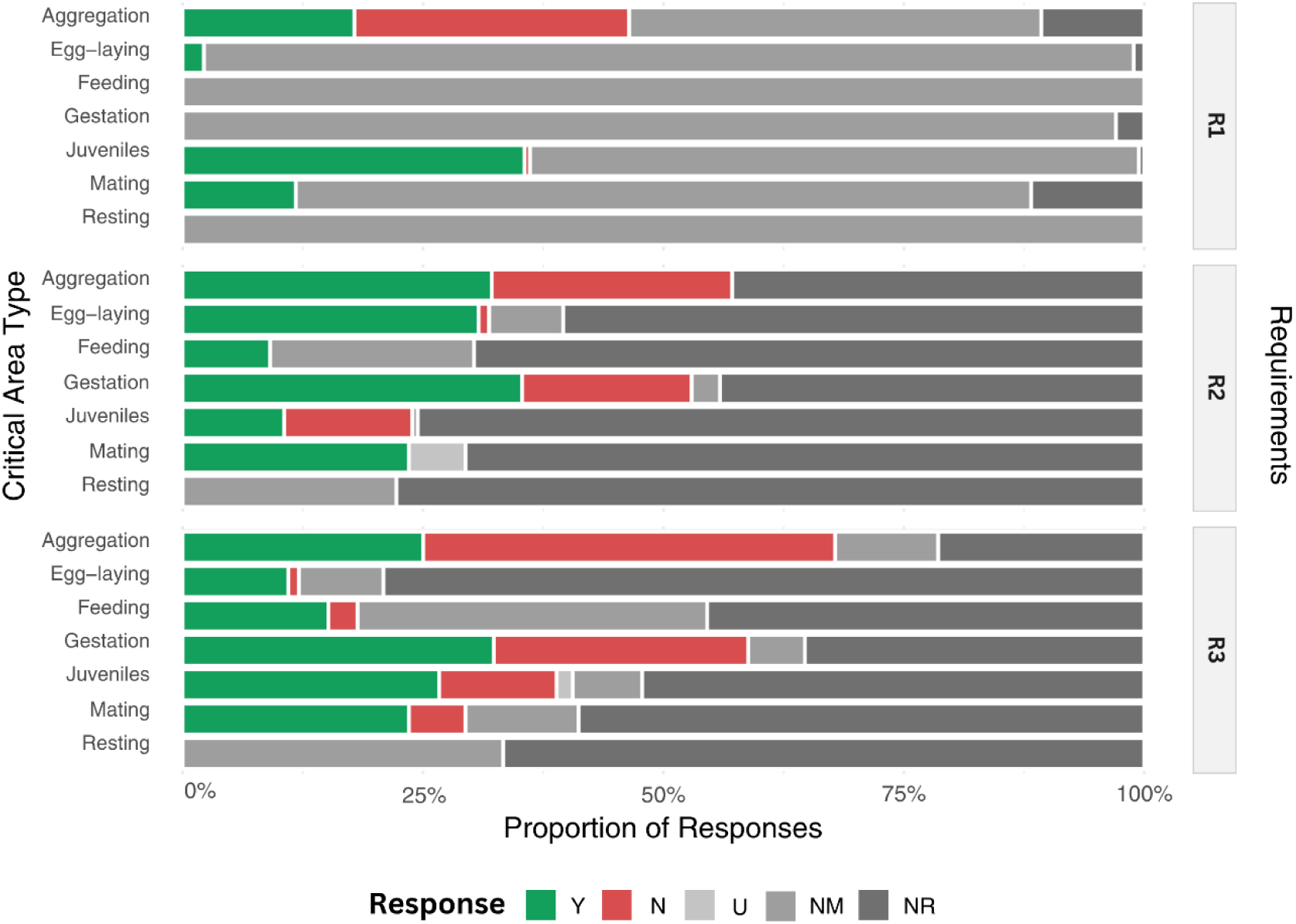
Proportion of responses (Y = Yes, N = No, U = Unsure, NM = Not Measured, NR = Not Reported) for each of the three critical area definition requirements proposed by this review.

Distinguishing genuinely low-importance areas (Q4) from data-poor areas (Q1 & Q3) was also challenging, as many requirements were assigned “not measured or reported” due to insufficient reporting resolution or data availability (Figure 13). Clearer evidence requirements and standardised reporting are therefore needed to better separate true low-importance sites from those limited by data scarcity, grouped here as *Weak/ Cannot Assess*. Confirming unimportance is inherently difficult, as it demands sufficiently thorough sampling to substantiate negative results. Amongst the methods employed by the included literature, acoustic telemetry may be the only method which enables temporally sufficient site-based examination of all three critical area requirements (R1 can be assessed by examining the proportion of tagged individuals using the area relative to other areas), assuming a representative sample size and multi-year dataset is available. Eight papers in this study employed this method with a sample size of ≥ 30 individuals (Table S9). Lower use regions of the receiver arrays in these studies [i.e. where critical area requirements (R1-R3) are not met], could therefore be tentatively categorised as “known unimportant” (Q4) and used to inform habitat models and spatial management. However, attention should be paid to potential sources of bias, such as receiver site selection.

Lack of reporting or measurement was common when critical area identification was not a primary study aim. Feeding areas were especially affected: ≈ 90 % of requirement data were NR or NM, leaving only 16 % of sites with *Moderate* or *Strong* evidence. This reflected the dominance of dietary studies (Biton-Porsmoguer, 2020; Brown-Vuillemin *et al*., 2020; Biton-Porsmoguer, 2022), primarily examining trophic relationships rather than area importance. However, the value of studies not directly focused on habitat identification was enhanced when detailed sampling metadata were provided, such as sampling location and total length of individuals (Lieber *et al.,* 2020; Biton-Porsmoguer, 2020).

#### Improving Data Quality and Expanding Evidence Bases

##### Data clarity and standardisation

The reviewed literature revealed terminology inconsistencies, such as interchangeable use of *critical* and *essential*, and vague reproductive terms such as *spawn*, *breed*, or *reproduce*. These inconsistencies hinder cross-study synthesis and obscure true area function. To support more consistent evaluation of critical area requirements and maximise available data in line with the highlighted evidence needs, this review proposes standardised research and reporting guidelines (Table 7), which can be included as supplementary material, when not directly relevant to study aims. For acoustic telemetry data, abacus plots partitioned by demographic group (sex, adults, juveniles, neonates, YOYs) and focal areas provide an efficient format to satisfy multiple reporting guidelines simultaneously.

**Table 7.**
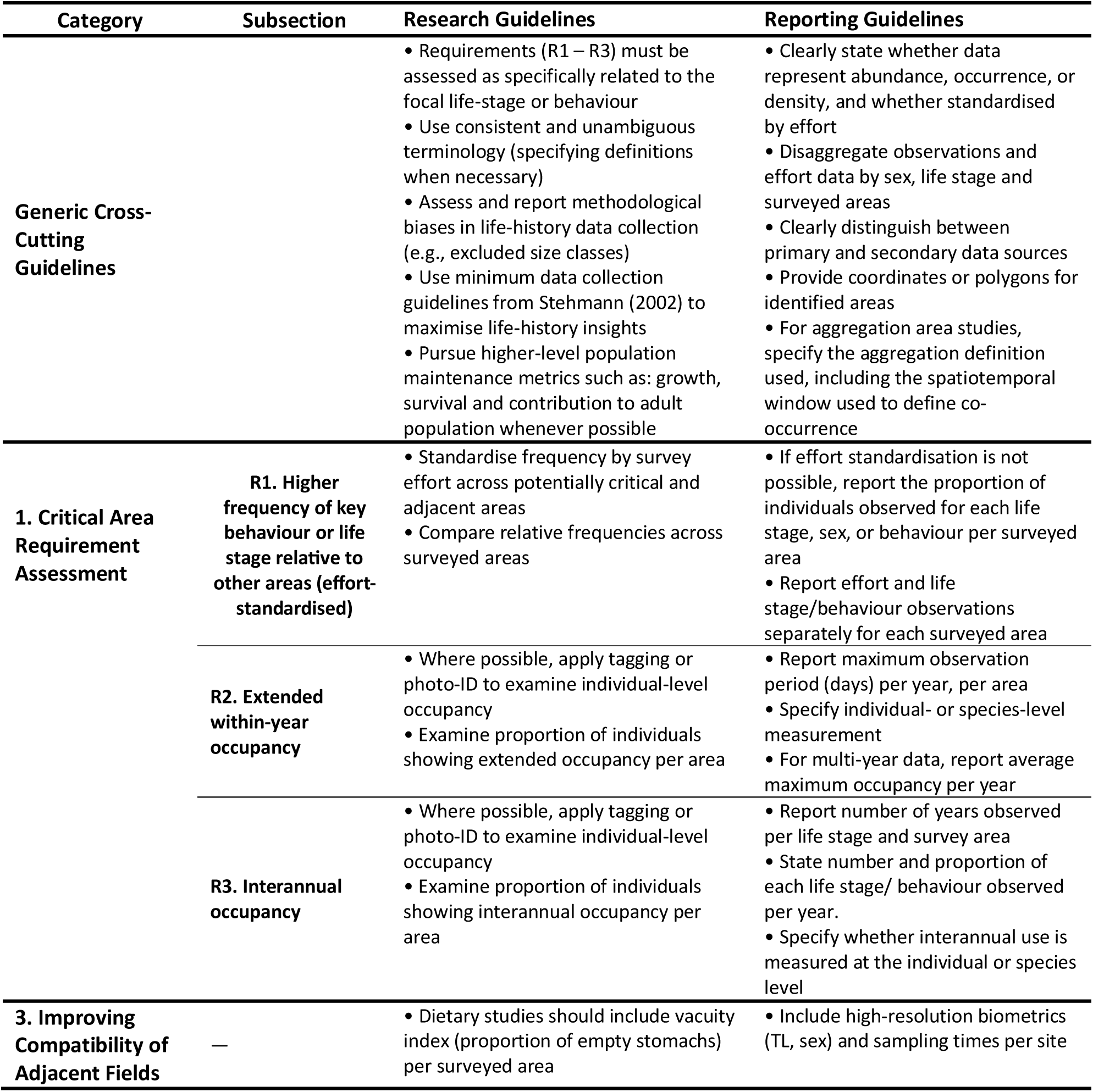
Standardised research and reporting guidelines for critical area research.

##### Broadening Evidence Inclusion

This review prioritised observational over modelled data to ensure that identified critical areas reflect realised habitat use, consistent with ISRA guidance (IUCN SSC Shark Specialist Group, 2024). While such studies were excluded from this review’s primary analysis, modelling remains valuable for contextualising habitat use over broad spatial scales, identifying potential critical areas in under-sampled regions (Figure 7), and anticipating climate-driven shifts (Faure-Beaulieu *et al*., 2023; Sun *et al*., 2024; Womersley *et al*., 2024; Ruiz-García *et al*., 2025). Therefore, while observational data remain essential for validating current realised habitat use, modelling represents a valuable complementary tool for prioritising field research and future-proofing management decisions.

Historical data (> 15 years old), although excluded from the main analysis, can substantially strengthen critical area evidence. Older sources may reveal potential critical areas for taxa absent from contemporary studies [such as *S. stellaris* in the case of the included literature (Orton, 1926)] and can reinforce contemporary evidence. For example, historical records of gravid *L. nasus* along the Celtic Shelf Break (Biais *et al., 2017*), when combined with *Moderate* contemporary evidence, strengthen support for its role as a gestation site. Incorporating historical research is particularly crucial for addressing shifting baseline syndrome (Pauly, 1995), ensuring that restoration ambitions are not constrained by currently depleted habitats (Moore & Hiddink, 2022).

Where modern observations are completely lacking for a species, applying the critical area definition (Table 1) to historical data could be used to identify “historically critical areas”, to guide contemporary observational research. For threatened species, if multiple requirements (R1-R3) are met by historical data and contemporary occurrence is also confirmed, these areas could serve as candidate sites for restoration efforts. Integrating historical and contemporary data in this way strengthens the overall evidence base, guides future research, and acknowledges formerly significant sites, while focusing management on areas that remain ecologically relevant today.

Another limitation arises when identification requires a clear area function, as this omits high-use sites (meeting habitat-use requirements R1–R3) of currently unknown function (see Table S10) where individuals are not interacting in close enough proximity to represent aggregation areas. Such sites may be particularly important for data-limited or threatened species [e.g. *Alopias vulpinus* in Western English Channel (Silva & Ellis, 2019)], whereby obtaining detailed life-stage or behavioural data is challenging. Applying R1–R3 independently of, or prior to, confirming function could help capture these areas of “high relative use” in future work, whilst maintaining strong evidence standards. Finally, cross-disciplinary synthesis, including studies not explicitly focused on critical area identification, could further strengthen the evidence base. Adoption of the proposed standardised reporting guidelines, especially for sampling metadata, will facilitate such integration.

##### Overall Evaluation of Testable Critical Area Definition

Applying the critical area definition (Table 1) facilitated a rapid and objective assessment of published peer-reviewed data, resulting in the identification of 103 areas with *Strong* or *Moderate* evidence. These areas were submitted for consideration in the ISRA process, and 94 % of the *Strong* areas were captured in the final delineations (84 % formally designated as ISRAs; Jabado *et al*., 2025). Discrepancies appeared largely to reflect the ISRA’s stricter requirements for early juvenile areas (considering only neonates/YOY), whereas this review included a broader range of immature life stages. This outcome underscores the utility of the critical area definition in helping researchers organise and present evidence in ways that support and complement spatial conservation planning processes.

While the generic thresholds applied inevitably oversimplified inter– and intraspecific variability in habitat use patterns, thresholds generally appeared to adequately filter out non-significant habitat use. Occasionally, they appeared too lenient, for instance *M. asterias* formed aggregations in most areas it occurred, suggesting that further comparisons of aggregation density, stability, or function may be required to highlight genuinely critical aggregation areas (Biton-Porsmoguer & Lloret, 2023). Conversely, the thresholds may be lowered for areas associated with short duration life-history events such as mating, courtship or use of a migratory corridor (McCauley, Papastamatiou & Young, 2010; Sequeira *et al*., 2025). However, even for mating, coincidental habitat use by both sexes is often necessary over extended timescales in line with the thresholds applied (Simpson *et al*., 2021; Sims *et al*., 2022; Lavender *et al*., 2022; Pratt *et al*., 2022). Meanwhile, migratory corridors may be better assessed via interannual occupancy (R3) and comparative density (R1) alone, since extended within-year occupancy (R2) is unlikely over small, management-relevant spatial scales.

Telemetry datasets are particularly powerful for refining region– and species-specific thresholds, as continuous tracking captures within– and inter-year occupancy more reliably than periodic surveys. As telemetry coverage expands taxonomically and spatially, thresholds can be tailored to species-and habitat type-specific patterns by establishing baseline occupancy, interannual use (Ellis et al., 2024), and density or frequency of use. However, if average habitat use is calculated, consideration should be given to potential sources of bias (e.g. receiver site selection, individual variability, spatial scale). This approach would ground area importance assessments in observed habitat use, rather than arbitrary cut-offs. Overall, in the absence of widely accepted species baselines, the thresholds applied here provide a practical reference for achieving comparability across data-limited species and regions.

##### Implications for Management and Conservation

This review highlights areas most suitable for spatial management based on extracted information from peer-reviewed literature. Priority candidate sites include those with *Strong* evidence associated with multiple-species (e.g., Bristol Channel and Irish Sea for immature skate areas) and or multiple overlapping functions (e.g., western Scotland). However, since this review excluded grey literature and focused on UK-priority chondrichthyan species (McCully Phillips, 2020) these results should be considered in combination with the final selected ISRA areas for the European Atlantic for greater taxonomic coverage and integration of unpublished data (Jabado *et al*., 2025).

Further, to predict the location of critical areas in under-surveyed regions and inform conservation policies that often target habitat features rather than chondrichthyan species (Council Directive 92/43/EEC, 2000; The Conservation of Habitats and Species Regulations, 2017), critical habitat variables could be extracted from *Strong* and *Moderate* evidence areas. This could be particularly valuable for species exhibiting highly specific critical habitat preferences in vulnerable habitats [e.g *G. melastomus* egg-laying in *L. pertusa* coral (Henry *et al*., 2013)].

Even *Strong* evidence single-species critical areas outside existing MPAs, or within MPAs with inadequate fisheries management, should be considered for management re-evaluation. Primary literature should be consulted for accurate area boundaries and seasonality of use, meanwhile social factors, such as appropriate fishery interventions and community support, should be integrated to enhance conservation effectiveness (MacKeracher *et al*., 2019; Renn *et al*., 2024). Strategies beyond MPAs, including seasonal closures, gear restrictions (e.g., tickler chains, escape hatches), or slot sizes to protect breeding stock, should also be considered (Kynoch *et al*., 2015; Campbell *et al*., 2020), particularly when these can achieve conservation objectives while imposing lower social costs on local communities. Vertical and migratory habitat use, though only briefly addressed here, further affect exposure to fishing threats and area connectivity, and should also inform spatial management decisions (Doherty *et al.,* 2017a; Griffiths *et al.,* 2020).

Regarding bio-physical drivers of MPA success (MacKeracher *et al*., 2019), conservation benefits ultimately depend on the feasibility of protecting the target critical area based on: a) the mobility of associated individuals and b) the contribution of the targeted individuals to overall population maintenance. While nursery areas are commonly prioritised for protection due to the lower dispersal of juveniles relative to adults (Ovenden, 2013; McMillan *et al*., 2021), evidence suggests that survival of age 0–1 individuals often has limited influence on population growth for many species (Heppell *et al*., 1999; Simpfendorfer, 1999; Frisk, 2002; Gallucci *et al*., 2006; Kinney & Simpfendorfer, 2009). Instead, protection of subadult and mature size classes may be more effective, especially for slow-growing species, which experience significant recovery lags following depletion of larger, reproductive size classes (Simpfendorfer & Kyne, 2009; Walker *et al*., 2021). Protection of said breeding stock underpins the few shark fisheries considered sustainable to date, such as “gauntlet fisheries” in South Australia, where a limited range of smaller size classes are harvested (Simpfendorfer, 1999; Prince, 2005). Rest areas for such size classes may constitute effective spatial management targets where dispersal is limited. Meanwhile, the protection of mating areas (Garla, Veras & Garrone-Neto, 2022; Sims *et al*., 2022) may also represent a key management priority since disruption of these sites could compromise reproductive success regardless of stock structure.

Generally, the best candidates for spatial management are smaller areas where individuals show strong site associations during life stages and/or functions that strongly influence population maintenance. Therefore, species-specific, size-structured population models should inform these decisions. In critical areas associated with high mobility life stages, site-independent fishery interventions may be more suitable (e.g. gear restrictions, modifications, reduced quota of species or their bycatch fisheries). Finally, to maintain protection throughout ontogeny, spatial management should aim for well-integrated MPA networks encompassing diverse critical area functions (Nagelkerken *et al*., 2015; Faure-Beaulieu *et al*., 2023).

## 5. Conclusions

1. The critical area definition proposed here provides a robust, generalisable framework to evaluate relative area importance, enabling researchers to objectively compare candidate critical areas. Its utility could be strengthened by developing species– and region-specific thresholds for importance based on normal habitat use levels. Where feasible, quantifying area-based population maintenance benefits (i.e. through growth, survival and adult contribution) would further supplement these assessments. Adoption of research and reporting guidelines will enhance consistency, comparability, and spatial coverage of future critical area research, increasing value to both research synthesis and spatial management decisions.
2. Application of the critical area definition identified potential critical areas with the strongest available evidence from the included peer-reviewed literature, providing a foundation to guide future research. Key biases and blind spots remain: offshore regions, species occupying deep waters (e.g. chimaeras), and habitats beyond early life stages are underrepresented, while positive-detection bias favours well-studied areas. Addressing these gaps requires systematic research targeting under-sampled regions, broader taxonomic and habitat coverage, and greater attention to high-use but functionally undefined areas. Meanwhile, greater attention to obtaining effort-standardised abundance data would increase reliability of future assessments.
3. Ultimately, critical area assessments are only as robust as available evidence. As datasets expand spatially, temporally (e.g. increased application of telemetry), and taxonomically, stricter evaluation standards, such as comparing habitat use to defined species-specific baseline levels, can be progressively applied, thereby increasing confidence in critical area delineation as evidence improves. By clarifying terminology and evidence requirements, and by highlighting priority research gaps, this review provides a roadmap for designing future studies that maximise their value to spatial management decisions and advance our understanding of chondrichthyan ecology.

## Funding

This work has received funding from the Natural Environment Research Council (NERC) through United Kingdom Research and Innovation (UKRI) for the Centre of Doctoral Training in Sustainable Management of UK Marine Resources (CDT SuMMeR) under grant agreement NE/W007215/1.

## Acknowledgements

The authors thank Kim Davis at the University of Plymouth for her expert guidance in systematic review methodology and support with literature searching, which greatly enhanced the quality and rigor of this study. We also extend our sincere thanks to Ryan Charles (ISRA IUCN SSG) for his significant support and expertise in improving the value of this manuscript to future area-based conservation decisions and facilitating close alignment with the ISRA process.

## Author contributions

**Conceptualisation:** C.R., A.E., B.C.

**Funding acquisition:** E.S.

**Investigation:** C.R.

**Methodology:** C.R., B.C.

**Software:** C.R.

**Supervision:** E.S., B.C., R.T., D.S., P.H.

**Validation:** C.R.

**Visualisation:** C.R.

**Writing – original draft preparation**: C.R.

**Writing – review & editing:** C.R., B.C., D.S., A.E., R.T., E.S.

## Notes

### Competing Interest Statement

The authors have declared no competing interest.

