## Supplementary Materials for "Tracks, Maps and Gaps: A Testable Research Definition for Critical Chondrichthyan Areas, European Atlantic Insights"

### Supplementary Tables

#### Table S1. IUCN Red list assessments

| Species | Europe IUCN Red listing | Global IUCN Red Listing | References |
|---|---|---|---|
| **Chimaeriformes** | | | |
| *Chimaera monstrosa* | NT | VU | Finucci, B. 2020. *Chimaera monstrosa*. *The IUCN Red List of Threatened Species* 2020: e.T63114A124459382.<br>Dagit, D.D. & Hareide, N.-R. 2015. *Chimaera monstrosa (Europe assessment)*. *The IUCN Red List of Threatened Species* 2015: e.T63114A48912471. |
| *Harriotta raleighana* | LC | LC | Finucci, B., Kulka, D.W. & Armstrong, A.O. 2025. *Harriotta raleighana*. *The IUCN Red List of Threatened Species* 2025: e.T278016856A268401245.<br>Buscher, E. & Walls, R. 2015. *Harriotta raleighana (Europe assessment)*. *The IUCN Red List of Threatened Species* 2015: e.T278016856A278017173. |
| *Hydrolagus mirabilis* | LC | LC | Finucci, B. 2020. *Hydrolagus mirabilis*. *The IUCN Red List of Threatened Species* 2020: e.T63104A124458962.<br>Walls, R. 2015. *Hydrolagus mirabilis (Europe assessment)*. *The IUCN Red List of Threatened Species* 2015: e.T63104A48909891. |
| *Hydrolagus pallidus* | LC | LC | Finucci, B. 2020. *Hydrolagus pallidus*. *The IUCN Red List of Threatened Species* 2020: e.T63103A124458813.<br>Finucci, B. 2020. *Hydrolagus pallidus*. *The IUCN Red List of Threatened Species* 2020: e.T63103A124458813. |
| *Hydrolagus affinis* | LC | LC | Finucci, B. 2020. *Hydrolagus affinis*. *The IUCN Red List of Threatened Species* 2020: e.T63123A124460832.<br>Walls, R. 2015. *Hydrolagus affinis (Europe assessment)*. *The IUCN Red List of Threatened Species* 2015: e.T63123A48917442. |
| *Rhinochimaera atlantica* | LC | LC | Finucci, B. 2020. *Rhinochimaera atlantica*. *The IUCN Red List of Threatened Species* 2020: e.T60145A124444407.<br>Walls, R. 2015. *Rhinochimaera atlantica (Europe assessment)*. *The IUCN Red List of Threatened Species* 2015: e.T60145A48939960. |
| Carcharhiniformes | | | |
| *Mustelus asterias* | NT | NT | Jabado, R.W., Ellis, J.R., McCully-Phillips, S.R., Dulvy, N.K., Farrell, E.D., Mancusi, C. & Derrick, D. 2021. *Mustelus asterias*. *The IUCN Red List of Threatened Species* 2021: e.T39357A124405496.<br>Farrell, E., McCully, S., Dulvy, N., Mancusi, C. & Ellis, J. 2015. *Mustelus asterias (Europe assessment)*. *The IUCN Red List of Threatened Species* 2015: e.T39357A48940630. Accessed on 01 December 2025. |
| *Galeus melastomus* | LC | LC | Farrell, E., McCully, S., Dulvy, N., Mancusi, C. & Ellis, J. 2015. *Mustelus asterias (Europe assessment)*. *The IUCN Red List of Threatened Species* 2015: e.T39357A48940630. Accessed on 01 December 2025.<br>Abella, A., Serena, F., Mancusi, C., Ungaro, N., Hareide, N.-R., Guallart, J., Coelho, R.P. & Crozier, P. 2015. *Galeus melastomus (Europe assessment)*. *The IUCN Red List of Threatened Species* 2015: e.T161398A48915013. Accessed on 01 December 2025. |
| *Scyliorhinus stellaris* | NT | VU | Finucci, B., Derrick, D. & Pacoureau, N. 2021. *Scyliorhinus stellaris*. *The IUCN Red List of Threatened Species* 2021: e.T161484A124493465.<br>Ellis, J., Serena, F., Mancusi, C., Haka, F., Morey, G., Guallart, J. & Schembri, T. 2015. *Scyliorhinus stellaris (Europe assessment)*. *The IUCN Red List of Threatened Species* 2015: e.T161484A48923567. Accessed on 01 December 2025. |
| Lamniformes | | | |
| *Lamna nasus* | CR | VU | Rigby, C.L., Barreto, R., Carlson, J., Fernando, D., Fordham, S., Francis, M.P., Herman, K., Jabado, R.W., Liu, K.M., Marshall, A., Pacoureau, N., Romanov, E., Sherley, R.B. & Winker, H. 2019. *Lamna nasus*. *The IUCN Red List of Threatened Species* 2019: e.T11200A500969.<br>Ellis, J., Farrell, E., Jung, A., McCully, S., Sims, D. & Soldo, A. 2015. *Lamna nasus (Europe assessment)*. *The IUCN Red List of Threatened Species* 2015: e.T11200A48916453. |
| *Alopias vulpinus* | EN | VU | Rigby, C.L., Barreto, R., Fernando, D., Carlson, J., Charles, R., Fordham, S., Francis, M.P., Herman, K., Jabado, R.W., Liu, K.M., Marshall, A., Pacoureau, N., Romanov, E., Sherley, R.B. & Winker, H. 2022. *Alopias vulpinus* (amended version of 2019 assessment). *The IUCN Red List of Threatened Species* 2022: e.T39339A212641186.<br>Ellis, J., Farrell, E., Jung, A., McCully, S., Sims, D. & Soldo, A. 2015. *Lamna nasus (Europe assessment)*. *The IUCN Red List of Threatened Species* 2015: e.T11200A48916453. |
| *Cetorhinus maximus* | EN | EN | Rigby, C.L., Barreto, R., Carlson, J., Fernando, D., Fordham, S., Francis, M.P., Herman, K., Jabado, R.W., Liu, K.M., Marshall, A., Romanov, E. & Kyne, P.M. 2021. *Cetorhinus maximus* (amended version of 2019 assessment). *The IUCN Red List of Threatened Species* 2021: e.T4292A194720078. Sims, D., |

| | | | |
|---|---|---|---|
| | | | Fowler, S.L., Clò, S., Jung, A., Soldo, A. & Bariche, M. 2015. *Cetorhinus maximus (Europe assessment)*. *The IUCN Red List of Threatened Species* 2015: e.T4292A48953216. |
| Myliobatiformes | | | |
| *Dasyatis pastinaca* | VU | VU | Jabado, R.W., Chartrain, E., De Bruyne, G., Derrick, D., Dia, M., Diop, M., Doherty, P., Leurs, G.H.L., Metcalfe, K., Pacoureau, N., Pires, J.D., Ratão, S., Seidu, I., Serena, F., Soares, A.-L., Tamo, A., VanderWright, W.J. & Williams, A.B. 2021. *Dasyatis pastinaca*. *The IUCN Red List of Threatened Species* 2021: e.T161453A124488102.<br>Serena, F., Mancusi, C., Morey, G. & Ellis, J.R. 2015. *Dasyatis pastinaca (Europe assessment)*. *The IUCN Red List of Threatened Species* 2015: e.T161453A48933979. |
| Rajiformes | | | |
| *Dipturus intermedius* | CR | CR | Ellis, J.R., McCully-Phillips, S.R., Sims, D., Walls, R.H.L., Cheok, J., Derrick, D. & Dulvy, N.K. 2024. *Dipturus intermedius* (amended version of 2021 assessment). *The IUCN Red List of Threatened Species* 2024: e.T18903491A256581177. |
| *Dipturus batis* | CR | CR | Ellis, J.R., McCully-Phillips, S.R., Sims, D., Derrick, D., Cheok, J. & Dulvy, N.K. 2024. *Dipturus batis* (amended version of 2021 assessment). *The IUCN Red List of Threatened Species* 2024: e.T203364219A256580832. |
| *Dipturus nidarosiensis* | NT | EN | Finucci, B., Lynghammar, A. & Charles, R. 2024. *Dipturus nidarosiensis*. *The IUCN Red List of Threatened Species* 2024: e.T161729A124534517.<br>Stehmann, M.F.W., Ellis, J., Walls, R. & Lynghammar, A. 2015. *Dipturus nidarosiensis (Europe assessment)* (errata version published in 2024). *The IUCN Red List of Threatened Species* 2015: e.T161729A258904627. |
| *Leucoraja circularis* | EN | EN | Finucci, B., Ellis, J.R., McCully-Phillips, S.R., Serena, F. & Charles, R. 2025. *Leucoraja circularis*. *The IUCN Red List of Threatened Species* 2025: e.T161464A124490090. |
| *Leucoraja fullonica* | VU | VU | Finucci, B., Rigby, C.L., Ellis, J.R., McCully-Phillips, S.R., Serena, F. & Charles, R. 2025. *Leucoraja fullonica*. *The IUCN Red List of Threatened Species* 2025: e.T161461A124489576. |
| *Leucoraja naevus* | LC | LC | Finucci, B., Ellis, J.R., McCully-Phillips, S.R., Pacoureau, N., Rohner, C.A. & Serena, F. 2025. *Leucoraja naevus*. *The IUCN Red List of Threatened Species* 2025: e.T161626A124517373. |
| *Raja brachyura* | NT | NT | Finucci, B., McCully-Phillips, S.R., Ellis, J.R., Serena, F., Soldo, A., Pacoureau, N. & Charles, R. 2024. *Raja brachyura*. *The IUCN Red List of Threatened Species* 2024: e.T161691A183779320.<br>McCully, S., Serena, F., Walls, R.H.L., Morey, G & Ellis, J.R. 2015. *Raja brachyura (Europe assessment)*. *The IUCN Red List of Threatened Species* 2015: e.T161691A48907330. Accessed on 01 December 2025. |
| *Raja microocellata* | NT | NT | Rigby, C.L., Ellis, J.R., McCully-Phillips, S.R & Rohner, C.A. 2025. *Raja microocellata*. *The IUCN Red List of Threatened Species* 2025: e.T39400A183780223.<br>McCully, S., Serena, F., Walls, R.H.L., Morey, G & Ellis, J.R. 2015. *Raja brachyura (Europe assessment)*. *The IUCN Red List of Threatened Species* 2015: e.T161691A48907330. Accessed on 01 December 2025. |
| *Raja montagui* | LC | LC | Rigby, C.L., McCully-Phillips, S.R., Ellis, J.R., Serena, F., Soldo, A., Pacoureau, N. & Charles, R. 2024. *Raja montagui*. *The IUCN Red List of Threatened Species* 2024: e.T63146A183780480.<br>Ellis, J.R., Walls, R.H.L., Serena, F. & Dulvy, N.K. 2015. *Raja montagui (Europe assessment)*. *The IUCN Red List of Threatened Species* 2015: e.T63146A48919726. Accessed on 01 December 2025. |
| *Raja undulata* | NT | NT | Finucci, B., McCully-Phillips, S.R., Ellis, J.R., Giovos, I., Serena, F., Soldo, A., Pacoureau, N. & García, E. 2024. *Raja undulata*. *The IUCN Red List of Threatened Species* 2024: e.T161425A183780918.<br>Ellis, J.R., McCully, S. & Walls, R.H.L. 2015. *Raja undulata (Europe assessment)*. *The IUCN Red List of Threatened Species* 2015: e.T161425A48909382. Accessed on 01 December 2025. |
| *Raja clavata* | NT | NT | Finucci, B., McCully-Phillips, S.R., Ellis, J.R., Giovos, I., Serena, F., Soldo, A., Pacoureau, N. & Charles, R. 2024. *Raja clavata*. *The IUCN Red List of Threatened Species* 2024: e.T39399A183779744.<br>Ellis, J., Dulvy, N., Walls, R. & Serena, F. 2016. *Raja clavata (Europe assessment)*. *The IUCN Red List of Threatened Species* 2016: e.T39399A103111648. Accessed on 01 December 2025. |
| *Rostroraja alba* | CR | EN | Serena, F., Charles, R., Ellis, J.R., Johnston, G. & Pacoureau, N. 2024. *Rostroraja alba*. *The IUCN Red List of Threatened Species* 2024: e.T61408A183781512.<br>Ellis, J., Morey, G & Walls, R. 2015. *Rostroraja alba (Europe assessment)*. *The IUCN Red List of Threatened Species* 2015: e.T61408A48954174. Accessed on 01 December 2025. |
| *Amblyraja radiata* | LC | VU | Kulka, D.W., Ellis, J.R., Anderson, B., Cotton, C.F., Derrick, D., Pacoureau, N. & Dulvy, N.K. 2020. *Amblyraja radiata*. *The IUCN Red List of Threatened Species* 2020: e.T161542A124503504.<br>Kulka, D.W., Ellis, J.R., Anderson, B., Cotton, C.F., Derrick, D., Pacoureau, N. & Dulvy, N.K. 2020. *Amblyraja radiata*. *The IUCN Red List of Threatened Species* 2020: e.T161542A124503504. |
| Squatiniformes | | | |
| *Squatina squatina* | CR | CR | Morey, G, Barker, J., Hood, A., Gordon, C., Bartolí, A., Meyers, E.K.M., Ellis, J.R., Sharp, R., Jiménez-Alvarado, D. & Pollom, R. 2019. *Squatina squatina*. *The IUCN Red List of Threatened Species* 2019: e.T39332A117498371. |

**Table S2. Studies identifying potentially critical movement areas.**

| Reference | Location | Species |
|---|---|---|
| **Sousa et al., 2019** | Between Professor Luiz Saldanha Marine Park and St. Andre Lagoon, Portugal | *Rostroraja alba* |
| **Tuya et al., 2020** | Las Canteras Beach, Gran Canaria | Several species |
| **Thorburn et al., 2021** | Loch Sunart to the Sound of Jura MPA, Scotland and surrounding waters. | *Dipturus intermedius* |
| **Thorburn et al., 2024** | Scotland, Republic of Ireland and Northern Ireland | *Cetorhinus maximus* |
| **Simpson et al., 2021** | Plymouth Coastal waters | *Raja brachyura,*<br>*Raja clavata,*<br>*Raja montagui,*<br>*Raja microocellata* |
| **Neat et al., 2015** | Sound of Jura and Surrounding waters | *Dipturus intermedius* |
| **McCully-Phillips & Ellis, 2015** | Southern North Sea & Eastern English Channel | *Mustelus asterias* |
| **Mead et al., 2023** | La Graciosa Marine Reserve, Canary Islands | *Squatina squatina* |
| **Meyers et al., 2017** | Canary Islands | *Squatina squatina* |
| **Leiber et al., 2020** | West Coast of Scotland | *Cetorhinus maximus* |
| **Leeb et al., 2021** | Ria de Vigo, Spain | *Raja undulata* |
| **Lavender et al., 2022** | Loch Sunart to the Sound of Jura MPA, Scotland | *Dipturus intermedius* |
| **Kraft et al., 2024** | Professor Luís Saldanha Marine Park (LSMP), Portugal | *Rostroraja alba* |
| **Kraft et al., 2023** | LSMP to the Reserva Natural do Estú ario do Sado, Portugal | *Dasyatis pastinaca* |
| **Griffiths et al., 2020** | English Channel | *Mustelus asterias* |
| **Dolton et al., 2020** | Between IoM and Scotland | *Cetorhinus maximus* |
| **Dolton et al., 2020** | Between British Isles and Norway | *Cetorhinus maximus* |
| **Dolton et al., 2020** | Between British Isles and Morocco | *Cetorhinus maximus* |
| **Doherty et al., 2017** | Continental Shelf off Western Ireland | *Cetorhinus maximus* |
| **Doherty et al., 2017** | Celtic Sea | *Cetorhinus maximus* |
| **Doherty et al., 2017** | Between UK and North Africa | *Cetorhinus maximus* |
| **Doherty et al., 2017** | Between UK and Iberian Peninsula | *Cetorhinus maximus* |
| **Doherty et al., 2017** | Between UK and Bay of Biscay | *Cetorhinus maximus* |
| **Doherty et al., 2017(a)** | Sea of Hebrides MPA, Scotland | *Cetorhinus maximus* |
| **Daban et al., 2024** | Cies Islands (Surrounding waters), Galicia | *Raja undulata* |
| **Carrasco-Puig et al., 2024** | Asturias, Bay of Biscay | *Lamna nasus* |
| **Cameron et al., 2019** | Northward Movement from Irish Waters | *Lamna nasus* |
| **Brevé et al., 2020** | Movement from English Channel (summer) to Bay of Biscay (In winter months) | *Mustelus asterias* |
| **Brevé et al., 2016** | Movement from English Channel (summer) to Bay of Biscay (In winter months) | *Mustelus asterias* |
| **Bortoluzzi et al., 2024** | Rockall Trough | *Lamna nasus* |
| **Bird et al., 2020** | southward movements from the south-eastern part of Division 6.a into 7.a, | Multiple skate species |
| **Bird et al., 2020** | UK waters | *Raja clavata* |
| **Bird et al., 2020** | UK waters | *Raja brachyura* |
| **Bird et al., 2020** | UK waters | *Raja montagui* |
| **Bird et al., 2020** | 6.a - 7.a (UK waters) | *Leucoraja naevus* |
| **Bird et al., 2020** | 7.e - 7.f (UK waters) | *Raja undulata* |
| **Biais et al., 2017** | Bay Biscay - Bay Biscay | *Lamna nasus* |
| **Biais et al., 2017** | NW Migration to Western Ireland | *Lamna nasus* |
| **Barker et al., 2022** | Welsh waters (Inshore/ Offshore movement) | *Squatina squatina* |

| | | |
|---|---|---|
| **Papadopoulo et al., 2023** | Between Ria de Vigo and South Cies Island MPA, Spain | *Raja clavata* |
| **Kraft et al., 2024** | Professor Luiz Saldanha Marine Park (LSMP) and the Setúbal peninsula and surrounding waters | *Raja clavata* |
| **Johnston et al., 2019** | Between Ireland and Cape Co, USA | *Cetorhinus maximus* |
| **Johnston et al., 2022** | Malin Head Ireland and UK/Irish coastal waters | *Cetorhinus maximus* |
| **Johnston et al., 2022** | Malin Head Ireland and tropical/ subtropical offshore waters | *Cetorhinus maximus* |

#### Table S3. Final search strings

| Database | Final Search string |
|---|---|
| **Web of Science core collection** | "Squatina squatina" OR "Dipturus intermedius" OR "Dipturus batis" OR "Leucoraja circularis" OR "Raja brachyura" OR "Raja microocellata" OR "Mustelus asterias" OR "Leucoraja fullonica" OR "Dipturus nidarosiensis" OR "Lamna nasus" OR "Leucoraja naevus" OR "Raja montagui" OR "Rostroraja alba" OR "Galeus melastomus" OR "Scyliorhinus stellaris" OR "Raja undulata" OR "Raja clavata" OR "Alopias vulpinus" OR "Cetorhinus maximus" OR "Amblyraja radiata" OR "Dasyatis pastinaca" OR “Chimaera monstrosa” OR “Hydrolagus mirabilis” OR “Hydrolagus pallidus” OR “Hydrolagus affinis” OR “Harriotta raleighana” OR “Rhinochimaera atlantica”<br>AND<br>Habitat* OR reproduc* OR breed* OR mate OR mating OR nursery OR nurseries OR egg* OR gestat* OR parturition OR juvenile* OR birth* OR pup* OR spawn* OR feed* OR diet* OR forag* OR hunt* OR develop* OR grow* OR rest* OR refug* OR movement OR migrat* OR aggregat* OR assemblage* OR diversity OR distinct* |
| **Scopus** | {Squatina squatina} OR {Dipturus intermedius} OR {Dipturus batis} OR {Leucoraja circularis} OR {Raja brachyura} OR {Raja microocellata} OR {Mustelus asterias} OR {Leucoraja fullonica} OR {Dipturus nidarosiensis} OR {Lamna nasus} OR {Leucoraja naevus} OR {Raja montagui} OR {Rostroraja alba} OR {Galeus melastomus} OR {Scyliorhinus stellaris} OR {Raja undulata} OR {Raja clavata} OR {Alopias vulpinus} OR {Cetorhinus maximus} OR {Amblyraja radiata} OR {Dasyatis pastinaca} OR {Chimaera monstrosa} OR {Hydrolagus mirabilis} OR {Hydrolagus pallidus} OR {Hydrolagus affinis} OR {Harriotta raleighana} OR {Rhinochimaera atlantica}<br>AND<br>Habitat* OR reproduc* OR breed* OR mate OR mating OR nursery OR nurseries OR egg* OR gestat* OR parturition OR juvenile* OR birth* OR pup* OR spawn* OR feed* OR diet* OR forag* OR hunt* OR develop* OR grow* OR rest* OR refug* OR movement OR migrat* OR aggregat* OR assemblage* OR diversity OR distinct* |

Searches used only accepted Latin names. A comparison using *Alopias vulpinus* (17 English common names listed on FishBase) demonstrated that inclusion of common names produced an order of magnitude more results (WoS: 504; Scopus: 733) than Latin names alone (WoS: 78; Scopus: 72), but these records contained a high proportion of irrelevant studies due to overlapping use of common names across species. To address the common misidentification of *Mustelus asterias* as *Mustelus mustelus*, only *M. asterias* was included in the search string, and all *M. mustelus* records in UK waters in the included studies were assumed to be *M. asterias* (Farrell, Clarke & Mariani, 2009). Furthermore, recent changes to the common skate complex fell within the search window; therefore, early results were interpreted with caution and generic habitat use data for the species-complex were excluded (Iglesias, Toulhoat & Sellos, 2010; Garbett *et al.*, 2023). Whilst the inclusion of the most recent *Dipturus batis* synonym, *Dipturus flossada*, yielded no additional results, *D. flossada* was used in some included studies for which the results were interpreted as *D. batis* records.

**Table S4. Inclusion/ exclusion criteria based on the Population, Concept, Context (PCC) framework (Peters et al., 2020).**

| | Inclusion criteria | Exclusion criteria |
|---|---|---|
| **Population** | The study includes habitat use information for one or more of the focal species (Table 2). | Study species are not listed in Table 2. |
| **Concept** | The study explicitly specified the identification of an important, or potentially important chondrichthyan habitat (based on the author's own interpretation of primary study data).<br><br>**OR**<br><br>The study identified a geographic area where a key life-stage, life history behaviour or event related to one or more ISRA categories potentially takes place. | The occurrence of chondrichthyans in the habitat is dependent on impermanent human influence in the area (e.g. food provisioning sites, oil & gas rigs, aquaculture sites).<br><br>No information is presented on the geographic location of the important habitat.<br><br>The important habitat was identified based on modelled data (e.g. species distribution modelling).<br><br>The important area is based on the identification of a subpopulation of chondrichthyans (i.e. genetically isolated group).<br><br>Feeding habitats are identified from stable isotopes alone, as which does not confirm recent feeding. |
| **Context** | One or more of the critical area observations were made in the European Atlantic (as delineated by the ISRA e-Atlas https://sharkrayareas.org/e-atlas/)<br><br>At least some of the study data must have been collected within the past 15 years (during or after January 2010) to represent contemporary critical areas. | Species observations were made within the European Atlantic, but any critical areas identified fell outside the European Atlantic. |
| **Other** | The article must be written in English.<br><br>The identification of the potential critical area(s) must be based on the interpretation of original data. | Non-English articles.<br><br>No primary data is presented (e.g. review papers). |

**Table S5. All included papers**

| Year | Author | Title |
|---|---|---|
| **2012** | Maia, C., Erzini, K., Serra-Pereira, B., Figueiredo, I. | Reproductive biology of cuckoo ray *Leucoraja naevus* |
| **2013** | Henry, L.-A., Navas, J., Hennige, S., et al. | Cold-water coral reef habitats benefit recreationally valuable sharks |
| **2014** | Serra-Pereira, B., Erzini, K., Maia, C., Figueiredo, I. | Identification of potential essential fish habitats for skates based on fishers' knowledge |
| **2015** | Cameron, L., Grabowski, J., Houghton, J., et al. | Population structure and spatial distribution of porbeagles (*Lamna nasus*) in Irish waters |

| | | |
|---|---|---|
| **2015** | Maia, C., Serra-Pereira, B., Erzini, K., et al. | How is the morphology of the oviducal gland and of the resulting egg capsule associated with the egg laying habitats of Rajidae species? |
| **2015** | McCully Phillips, S., Ellis, J. | Reproductive characteristics and life-history relationships of starry smooth-hound *Mustelus asterias* in British waters |
| **2015** | Neat, F., Burns, F., Jones, E., et al. | The diversity, distribution and status of deep-water elasmobranchs in the Rockall Trough, north-east Atlantic Ocean |
| **2015** | Neat, F., Pinto, C., Burrett, I., et al. | Site fidelity, survival and conservation options for the threatened flapper skate (*Dipturus cf. intermedia*) |
| **2015** | Silva, J., Ellis, J. | Bycatch and discarding patterns of dogfish and sharks taken in English and Welsh commercial fisheries |
| **2016** | Brevé, N., Winter, H., Van Overzee, H., et al. | Seasonal migration of the starry smooth-hound shark *Mustelus asterias* as revealed from tag-recapture data of an angler-led tagging programme |
| **2016** | Brevé, N., Winter, H., Wijmans, P., et al. | Sex differentiation in seasonal distribution of the starry smooth-hound *Mustelus asterias* |
| **2016** | Gore, M., Frey, P., Ormond, R., et al. | Use of photo-identification and mark-recapture methodology to assess basking shark (*Cetorhinus maximus*) populations |
| **2016** | Ponte, D., Bárcelos, L., Santos, C., et al. | Diet of *Dasyatis pastinaca* and *Myliobatis aquila* (Myliobatiformes) from the Azores, NE Atlantic |
| **2017** | Biais, G., Coupeau, Y., Séret, B., et al. | Return migration patterns of porbeagle shark (*Lamna nasus*) in the Northeast Atlantic: Implications for stock range and structure |
| **2017** | Couto, A., Queiroz, N., Relvas, P., et al. | Occurrence of basking shark *Cetorhinus maximus* in southern Portuguese waters: A two-decade survey |
| **2017** | Doherty, P., Baxter, J., Gell, F., et al. | Long-term satellite tracking reveals variable seasonal migration strategies of basking sharks in the north-east Atlantic |
| **2017** | Doherty, P., Baxter, J., Godley, B., et al. | Testing the boundaries: Seasonal residency and inter-annual site fidelity of basking sharks in a proposed Marine Protected Area |
| **2017** | Meyers, E., Tuya, F., Barker, J., et al. | Population structure, distribution and habitat use of the Critically Endangered angelshark, *Squatina squatina*, in the Canary Islands |
| **2018** | Benjamins, S., Dodd, J., Thorburn, J., et al. | Evaluating the potential of photo-identification as a monitoring tool for flapper skate (*Dipturus intermedius*) |
| **2018** | Hayes, E., Godley, B., Nimak-Wood, M., et al. | Basking shark breaching behaviour observations west of Shetland |
| **2019** | Gore, M., Abels, L., Wasik, S., et al. | Are close-following and breaching behaviours by basking sharks at aggregation sites related to courtship? |
| **2019** | Johnston, E., Mayo, P., Mensink, P., et al. | Serendipitous re-sighting of a basking shark *Cetorhinus maximus* reveals inter-annual connectivity between American and European coastal hotspots |
| **2019** | Sousa, I., Baeyaert, J., Gonçalves, J., et al. | Preliminary insights into the spatial ecology and movement patterns of a regionally critically endangered skate (*Rostroraja alba*) associated with a marine protected area |
| **2019** | Silva, J., Ellis, J. | Bycatch and discarding patterns of dogfish and sharks taken in English and Welsh commercial fisheries |
| **2020** | Biton-Porsmoguer, S. | Fisheries and Ecology of the Skates (Rajiformes: Rajidae) in the English Channel |
| **2020** | Bird, C., Burt, G., Hampton, N., et al. | Fifty years of tagging skates (Rajidae): Using mark-recapture data to evaluate stock units |
| **2020** | Brown-Vuillemin, S., Barreau, T., Caraguel, J., et al. | Trophic ecology and ontogenetic diet shift of the blue skate (*Dipturus cf. flossada*) |
| **2020** | Dolton, H., Gell, F., Hall, J., et al. | Assessing the importance of Isle of Man waters for the basking shark *Cetorhinus maximus* |
| **2020** | Frost, M., Neat, F., Stirling, D., et al. | Distribution and thermal niche of the common skate species complex in the north-east Atlantic |
| **2020** | Griffiths, C., Wright, S., Silva, J., et al. | Horizontal and vertical movements of starry smooth-hound *Mustelus asterias* in the northeast Atlantic |

| | | |
|---|---|---|
| **2020** | Jiménez-Alvarado, D., Meyers, E., Caro, M., et al. | Investigation of juvenile angelshark (*Squatina squatina*) habitat in the Canary Islands with recommended measures for protection and management |
| **2020** | Lieber, L., Hall, G., Hall, J., et al. | Spatio-temporal genetic tagging of a cosmopolitan planktivorous shark provides insight to gene flow, temporal variation and site-specific re-encounters |
| **2020** | Tuya, F., Asensio, M., Navarro, A. | “Urbanite” rays and sharks: Presence, habitat use and population structure in an urban semi-enclosed lagoon |
| **2021** | Leeb, K., Villegas-Ríos, D., Mucientes, G., et al. | Drivers of spatial behaviour of the endangered undulate skate, *Raja undulata* |
| **2021** | McAllister, M., Fraser, S., Henry, L.-A. | Population ecology and juvenile density hotspots of thornback ray (*Raja clavata*) around the Shetland Islands, Scotland |
| **2021** | Phillips, N., Garbett, A., Wise, D., et al. | Evidence of egg-laying grounds for critically endangered flapper skate (*Dipturus intermedius*) off Orkney, UK |
| **2021** | Rudd, J., Bartolomeu, T., Dolton, H., et al. | Basking shark sub-surface behaviour revealed by animal-towed cameras |
| **2021** | Santos, R., Medeiros-Leal, W., Novoa-Pabon, A., et al. | Biological knowledge of thornback ray (*Raja clavata*) from the Azores: Improving scientific information for the effectiveness of species-specific management measures |
| **2021** | Simpson, S., Humphries, N., Sims, D. | Habitat selection, fine-scale spatial partitioning and sexual segregation in Rajidae, determined using passive acoustic telemetry |
| **2021** | Thorburn, J., Wright, P., Lavender, E., et al. | Corrigendum: Seasonal and ontogenetic variation in depth use by a critically endangered benthic elasmobranch and its implications for spatial management |
| **2021** | Thorburn, J., Wright, P., Lavender, E., et al. | Seasonal and ontogenetic variation in depth use by a critically endangered benthic elasmobranch and its implications for spatial management |
| **2022** | Biton-Porsmoguer, S. | Diet strategies of starry smooth-hound *Mustelus asterias* and tope shark *Galeorhinus galeus* (Carcharhiniformes: Triakidae) in the Eastern English Channel: Implications for conservation |
| **2022** | Delaval, A., Frost, M., Bendall, V., et al. | Population and seascape genomics of a critically endangered benthic elasmobranch, the blue skate *Dipturus batis* |
| **2022** | Dodd, J., Baxter, J., Donnan, D., et al. | First report of an egg nursery for the critically endangered flapper skate *Dipturus intermedius* (Rajiformes: Rajidae) |
| **2022** | Johnston, E., Houghton, J., Mayo, P., et al. | Cool runnings: Behavioural plasticity and the realised thermal niche of basking sharks |
| **2022** | Lavender, E., Aleynik, D., Dodd, J., et al. | Movement patterns of a critically endangered elasmobranch (*Dipturus intermedius*) in a Marine Protected Area |
| **2022** | Sims, D., Berrow, S., O'Sullivan, K., et al. | Circles in the sea: annual courtship “torus” behaviour of basking sharks *Cetorhinus maximus* identified in the eastern North Atlantic Ocean |
| **2022** | Barker, J., Davies, J., Goralczyk, M., et al. | The distribution, ecology and predicted habitat use of the critically endangered angelshark (*Squatina squatina*) in coastal waters of Wales and the central Irish Sea |
| **2023** | Biton-Porsmoguer, S., Lloret, J. | Estimating the effects of recreational fisheries on sharks in the English Channel and adjacent seas using social networks |
| **2023** | Fontaine, P., Barreiros, J., Jaquemet, S. | Trophic ecology of three sympatric batoid species (*Dasyatis pastinaca*, *Raja clavata*, and *Raja maderensis*) from the Azores, NE Atlantic |
| **2023** | Kraft, S., Winkler, A.C., Abecasis, D. | Small coastal marine protected areas offer recurring, seasonal protection to the common stingray (*Dasyatis pastinaca*) |
| **2023** | Mead, L., Alvarado, D., Meyers, E., et al. | Spatiotemporal distribution and sexual segregation in the critically endangered angelshark *Squatina squatina* in Spain’s largest marine reserve |
| **2023** | Papadopoulo, K., Villegas-Ríos, D., Mucientes, G., et al. | Drivers of the spatial behaviour of the threatened thornback skate (*Raja clavata*) |
| **2023** | Thorburn, J., Cole, G., Naylor, A., et al. | Preliminary insight into the reproductive traits of the flapper skate *Dipturus intermedius* using in-field ultrasonography and circulating hormone concentrations |
| **2023** | Thorburn, J., Collins, P., Garbett, A., et al. | Assessing the potential of acoustic telemetry to underpin the regional management of basking sharks (*Cetorhinus maximus*) |

| | | |
|---|---|---|
| **2024** | Bortoluzzi, J., McNicholas, G., Jackson, A., et al. | Transboundary movements of porbeagle sharks support need for continued cooperative research and management approaches |
| **2024** | Carrasco-Puig, P., Miralles, L., Colmenero, A., et al. | Occurrence of juvenile porbeagle sharks (*Lamna nasus*) in the northern coast of Spain |
| **2024** | Daban, P., Hillinger, A., Mucientes, G., et al. | Movement ecology determines isotopic niche width in the undulate skate *Raja undulata* |
| **2024** | Jung, A., Ory, A., Abaut, P., et al. | First use of free-diving photo-identification of porbeagle shark (*Lamna nasus*) off the Brittany coast, France |
| **2024** | Kraft, S., Winkler, A.C., Abecasis, D., et al. | Long-term co-occurrence and gregariousness in the migratory common stingray using network analysis |
| **2024** | Kraft, S., Winkler, A.C., Abecasis, D. | Seasonal movement dynamics of the commercially important thornback ray (*Raja clavata*) in a coastal marine protected area |
| **2024** | Kraft, S., Winkler, A.C., Abecasis, D. | Horizontal and vertical movements of the critically endangered *Rostroraja alba* in a coastal marine protected area |
| **2024** | Rodríguez-García, C., Gonçalves Neto, J., García-Romero, C., et al. | Feeding habits of two shark species: velvet belly *Etmopterus spinax* and blackmouth catshark *Galeus melastomus*, present in fishing discards in the Gulf of Cádiz |
| **2024** | Schwanck, T., Vizer, L., Thorburn, J., et al. | Mitochondrial haplotypes reveal low diversity and restricted connectivity of the critically endangered batoid population in a Marine Protected Area |

#### Table S6. Size at birth, and size at maturity

| *Species* | *Size at Birth / Hatching* | *YOY / First-Cohort Cut-off* | *Size at Maturity* |
|---|---|---|---|
| ***Amblyraja radiata*** | *8–12 cm*[1] | *≤14 cm*[2] | *36–88 cm*[1] |
| ***Cetorhinus maximus*** | *150–200 cm*[1] | *≤240 cm*[3,4] | *Females >800 cm, Males 750–800 cm*[1] |
| ***Dipturus batis*** | *21–29 cm*[1] | — | *115–125 cm*[1] |
| ***Dipturus intermedius*** | *21–29 cm*[1] | — | *Females 197 cm, Males 185 cm*[1] |
| ***Dipturus nidarosiensis*** | *24–28 cm*[1] | — | *~120 cm*[7] |
| ***Galeus melastomus*** | *7.5–9 cm*[5,6] | — | *Females 39–45 cm, Males 34–42 cm*[1] |
| ***Lamna nasus*** | *60–80 cm*[1] | *≤98 cm*[8] | *Females 245 cm, Males 195 cm*[1] |
| ***Leucoraja circularis*** | — | — | *Males 66 cm, Females >76 cm*[1] |
| ***Leucoraja fullonica*** | *15–20 cm*[1] | — | *Females >82 cm, Males 75–81 cm*[1] |
| ***Leucoraja naevus*** | *9–12 cm*[1] | *≤15 cm*[2] | *Females 53–60 cm, Males 50–57 cm*[1] |
| ***Mustelus asterias*** | *28–32 cm*[1] | *≤50 cm*[9] | *Females 83–96 cm, Males 72–85 cm*[1] |
| ***Raja brachyura*** | *16–18 cm*[1] | *≤19 cm*[2] | *80–90 cm*[1] |
| ***Raja clavata*** | *10–13 cm*[1] | *≤15 cm*[2] | *Females 60–85 cm, Males 60–77 cm*[1] |
| ***Raja microocellata*** | *10–13 cm*[1] | *≤22 cm*[2] | *~58 cm*[1] |
| ***Raja montagui*** | *8–10 cm*[1] | *≤16 cm*[2] | *Females 49–64 cm, Males 40–50 cm*[1] |
| ***Raja undulata*** | *~14 cm*[1] | *≤19 cm (Females)*[2] | *Females 76–84 cm, Males 74–78 cm*[1] |
| ***Rostroraja alba*** | *~30 cm*[1] | — | *Females 130–195 cm, Males 120–170 cm*[1] |
| ***Squatina squatina*** | *24–30 cm*[1] | — | *Females 126–167 cm, Males 80–132 cm*[1] |

This table was primarily populated using values from Ebert & Dando (2021) and, gaps were filled using data from Ellis *et al.,* (2024), particularly for size at 1st cohort. However, in some cases this value was smaller than the upper limit for size at birth (e.g. *D. batis; D. intermedius*), in which case individuals of this size were classed as neonates. Remaining gaps were populated by consulting biological data from the most recent species-specific IUCN Red List of Threatened Species assessment pages (Pauly, 1978; 2002; Costa et al., 2005; Capapé et al., 2007). When no length data were specified, or no lower bound was given (e.g. < 40 cm) individuals were classed as immature (potentially containing any sized immature individuals).

[1] Ebert, D.A. & Dando, M. (2021) Field guide to sharks, rays, and chimaeras of Europe and the Mediterranean. Princeton University Press, Princeton.
[2] Ellis, J.R., Gordon, C.A., Allen, H.L., Silva, J.F., Bird, C., Johnston, G., O'Connor, B., McCully Phillips, S.R. & Hood, A. (2024) The distribution of the juvenile stages and eggcases of skates (Rajidae) around the British Isles. Aquatic Conservation: Marine and Freshwater Ecosystems.
[3] Pauly, D. (1978) A critique of some literature on the growth, reproduction and mortality of the lamnid shark Cetorhinus maximus (Gunnerus). ICES Pelagic Fish Committee paper CM 1978/H:17.
[4] Pauly, D. (2002) Growth and mortality of the basking shark *Cetorhinus maximus* and their implications for management of

whale sharks *Rhincodon typus* . In: SL Fowler, TM Reed and FA Dipper (eds), *Elasmobranch Biodiversity, Conservation and Management. Proceedings of the International Seminar and Workshop, Sabah, Malaysia, July 1997* , pp. 199–208. IUCN SSC Shark Specialist Group. IUCN, Gland, Switzerland and Cambridge, UK.
[5]Costa M.E., Erzini, K. and Borges, T.C. (2005) Reproductive biology of the blackmouth catshark, *Galeus melastomus* (Chondrichthyes: Scyliorhinidae) off the south coast of Portugal. *Journal of the Marine Biological Association of the United Kingdom* 85(5): 1173-1183.
[6]Capapé C, Guélorget O, Vergne Y, Reynaud C. Reproductive biology of the blackmouth catshark, *Galeus melastomus* (Chondrichthyes: Scyliorhinidae) off the Languedocian coast (southern France, northern Mediterranean). *Journal of the Marine Biological Association of the United Kingdom* 88(2):415-421.
[7]Follesa, M.C., Cannas, R., Cabiddu, S., Cau, A., Mulas, A., Porcu, C. and Cau, A. (2012) Preliminary observations of the reproductive biology and diet for the Norwegian skate *Dipturus nidarosiensis* (Rajidae) from the central western Mediterranean Sea. *Cybium* 36: 473–477.
[8]Francis, M., Natantson, L.J. and Campana, S.E. (2009) Chapter 9, The Biology and Ecology of the Porbeagle Shark, *Lamna Nasus.*In *Sharks of the Open Ocean, Biology, Fisheries and Conservation* (eds M.D. Camhi, E.K. Pikitch and E.A. Babcock). Blackwell Publishing, Oxford, UK.
[9]Farrell, E.D., Mariani, S. & Clarke, M.W. (2010) Reproductive biology of the starry smooth-hound shark *Mustelus asterias*: geographic variation and implications for sustainable exploitation. Journal of Fish Biology 77, 1505–1525.

**Table S7. Risk of bias calculation example (viviparous gestation areas) and abbreviation meanings**

| Reference | Location | Species | Mult. Ind. | Effort | Resp. Var. | Evidence | Repr. | Obs. Type | Interp. | Score | % | Bias | Evidence Cap |
|---|---|---|---|---|---|---|---|---|---|---|---|---|---|
| Silva & Ellis, 2019 | Bristol Channel (ICES 30E5, 31E5–31E6) | *Mustelus asterias* | NR | H | O | E | U | DO | NA | 3.5 | 58.3 | Med | Large females |
| Mead et al., 2023 | La Graciosa Reserve, Canary Islands | *Squatina squatina* | Y | H | M | E | L | BI | L | 4 | 57.1 | Med | Females |
| McCully-Phillips & Ellis, 2015 | W. English Channel, French Coast | *Mustelus asterias* | N | H | O | E | U | DO | NA | 3.5 | 58.3 | Med | Pregnant females |
| Kraft et al., 2023 | Sado Estuary, Portugal | *Dasyatis pastinaca* | Y | H | M | E | H | BI | L | 5 | 71.4 | Low | Mature females |
| Jung et al., 2024 | Tregor Area, France | *Lamna nasus* | Y | H | A | E | H | DO | NA | 5.5 | 91.7 | Low | Pregnant females |

| Column | Meaning |
|---|---|
| **Mult. Ind.** | Multiple individuals observed (Y = Yes, N = No, NR = Not Reported) |
| **Resp. Var.** | Response variable (O = Occurrence, M = Movement, A = Aggregation) |
| **Evidence** | Anecdotal or Empirical (E = Empirical, A = Anecdotal) |
| **Repr.** | Representation of life history stage in sample (U = Unsure, L = Low, H = High) |
| **Obs. Type** | Observation type (DO = Direct Observation, BI = Behavioural interpretation) |
| **Interp.** | Strength of behavioural interpretation (L = Low, H = High) |
| **Score** | Total score |
| **%** | Percentage of highest possible score |
| **Bias** | Overall risk of bias |
| **Evidence Cap** | Is a cap placed on overall score introduced due to evidence available (Green = No Cap, Red = Cap) |

**Table S8. All potential critical areas identified from the literature, and associated requirements (R1 – R3) met and risk of bias score**

*In separate pdf document.*

**Table S9. Included acoustic telemetry studies with sufficient sampling to identify "known unimportant areas" (Q4; Figure12).**

| Year | Author | Title |
|---|---|---|

| | | |
|---|---|---|
| **2021** | Leeb, Villegas-Ríos, Mucientes, et al. | Drivers of spatial behaviour of the endangered undulate skate, *Raja undulata* |
| **2021** | Simpson, Humphries, Sims | Habitat selection, fine-scale spatial partitioning and sexual segregation in Rajidae, determined using passive acoustic telemetry |
| **2023** | Kraft, Winkler, Abecasis | Small coastal marine protected areas offer recurring, seasonal protection to the common stingray (*Dasyatis pastinaca*) |
| **2023** | Mead, Alvarado, et al. | Spatiotemporal distribution and sexual segregation in the critically endangered angelshark *Squatina squatina* in Spain's largest marine reserve |
| **2024** | Daban, Hillinger, Mucientes, et al. | Movement ecology determines isotopic niche width in the undulate skate *Raja undulata* |
| **2024** | Kraft, Winkler, Abecasis, Mourier | Long-term co-occurrence and gregariousness in the migratory common stingray using network analysis |
| **2024** | Kraft, Winkler, Abecasis | Seasonal movement dynamics of the commercially important thornback ray (*Raja clavata*) in a coastal marine protected area |
| **2024** | Kraft, Winkler, Abecasis | Horizontal and vertical movements of the critically endangered *Rostroraja alba* in a coastal marine protected area |
| **2024** | Thorburn, Collins, Garbett, et al. | Assessing the potential of acoustic telemetry to underpin the regional management of basking sharks (*Cetorhinus maximus*) |

**Table S10. Potential high use areas with undefined function, not captured by any *Strong* or *Moderate* evidence areas in the primary analysis**

| Reference | Location (s) | Species |
|---|---|---|
| **Tuya et al., 2020** | Las Canteras Beach, Gran Canaria | *Squatina squatina* |
| **Thorburn et al., 2024** | Malin-Islay Front, Ireland/Scotland | *Cetorhinus maximus* |
| **Simpson et al., 2021** | Coastal waters off Bantham, England | *Raja brachyura* |
| **Simpson et al., 2021** | Coastal waters off Bigbury-on-sea, England | *Raja clavata* |
| **Simpson et al., 2021** | Coastal waters off Wembury, England | *Raja montagui* |
| **Simpson et al., 2021** | Plymouth Sound, England | *Raja microocellata* |
| **Silva & Ellis, 2019** | Celtic Seas Ecoregion | *Mustelus asterias* |
| **Silva & Ellis, 2019** | Celtic Seas Ecoregion | *Galeus melastomus* |
| **Silva & Ellis, 2019** | Western English Channel | *Alopias vulpinus* |
| **Silva & Ellis, 2019** | Celtic Seas Ecoregion | *Scyliorhinus stellaris* |
| **Shephard et al., 2021** | Tralee Bay | *Squatina squatina* |
| **Shephard et al., 2021** | Clew Bay | *Squatina squatina* |
| **Neat et al., 2015** | Rockall Trough | *Dipturus nidarioensis* |
| **Neat et al., 2015** | Rockall Trough | *Galeus melsatomus* |
| **Lieber et al., 2020** | Ireland | *Cetorhinus maximus* |
| **Leeb at al., 2021** | Ria de Vigo, Spain | *Raja undulata* |
| **Kraft et al., 2023** | Setúbal peninsula, Portugal | *Dasyatis pastinaca* |
| **Hiddink et al., 2019** | Cardigan Bay, Wales | *Squatina squatina* |
| **Griffiths et al., 2020** | Northern Bay of Biscay | *Mustelus asterias* |
| **Griffiths et al., 2020** | Celtic Sea | *Mustelus asterias* |
| **Frost et al., 2020** | Rockall Trough | *Dipturus batis* |
| **Frost et al., 2020** | Western Scotland (Especially West of Tiree) | *Dipturus intermedius* |
| **Dolton et al., 2020** | Moroccan EEZ | *Cetorhinus maximus* |
| **Delaval et al., 2023** | Isles of Scilly, UK | *Dipturus batis* |
| **Daban et al., 2024** | Cies Islands, Galicia | *Raja undulata* |

| | | |
|---|---|---|
| **Bortoluzzi et al., 2024** | North of Ireland | *Lamna nasus* |
| **Bortoluzzi et al., 2024** | Celtic Deep | *Lamna nasus* |
| **Bortoluzzi et al., 2024** | Irish Sea | *Lamna nasus* |
| **Bortoluzzi et al., 2024** | North & East Coast of UK | *Lamna nasus* |
| **Biton-Porsmoguer et al., 2023** | Eastern English Channel | *Raja clavata* |
| **Biton-Porsmoguer et al., 2023** | Estuaries Picards & the Opal Sea | *Raja clavata* |
| **Biton-Porsmoguer et al., 2023** | North Sea/Detroit Pas de Calais | *Raja clavata* |
| **Biton-Porsmoguer et al., 2023** | Estuaries Picards & the Opal Sea | *Raja montagui* |
| **Biton-Porsmoguer et al., 2023** | North Sea/Detroit Pas de Calais | *Raja montagui* |
| **Bird et al., 2020** | Outer Thames, UK | *Raja clavata* |
| **Bird et al., 2020** | 6.a (UK) | *Raja brachyura* |
| **Bird et al., 2020** | 7.e(UK) | *Raja brachyura* |
| **Bird et al., 2020** | 4.c(UK) | *Raja montagui* |
| **Bird et al., 2020** | 6.a (UK) | *Leucoraja naevus* |
| **Bird et al., 2020** | 7.e(UK) | *Raja undulata* |
| **Bird et al., 2020** | 7.h (UK) | *Dipturus spp.* |
| **Bird et al., 2020** | 4.c (UK) | *Amblyraja radiata* |
| **Bird et al., 2020** | 7.h (UK) | *Leucoraja fullonica* |
| **Barker et al., 2022** | Welsh waters | *Squatina squatina* |
| **Serra-Pereira et al., 2014** | Off Santa Cruz, Portugal | *Raja brachyura* |

### Supplementary Figures:

#### Figure S1. Risk of bias tool

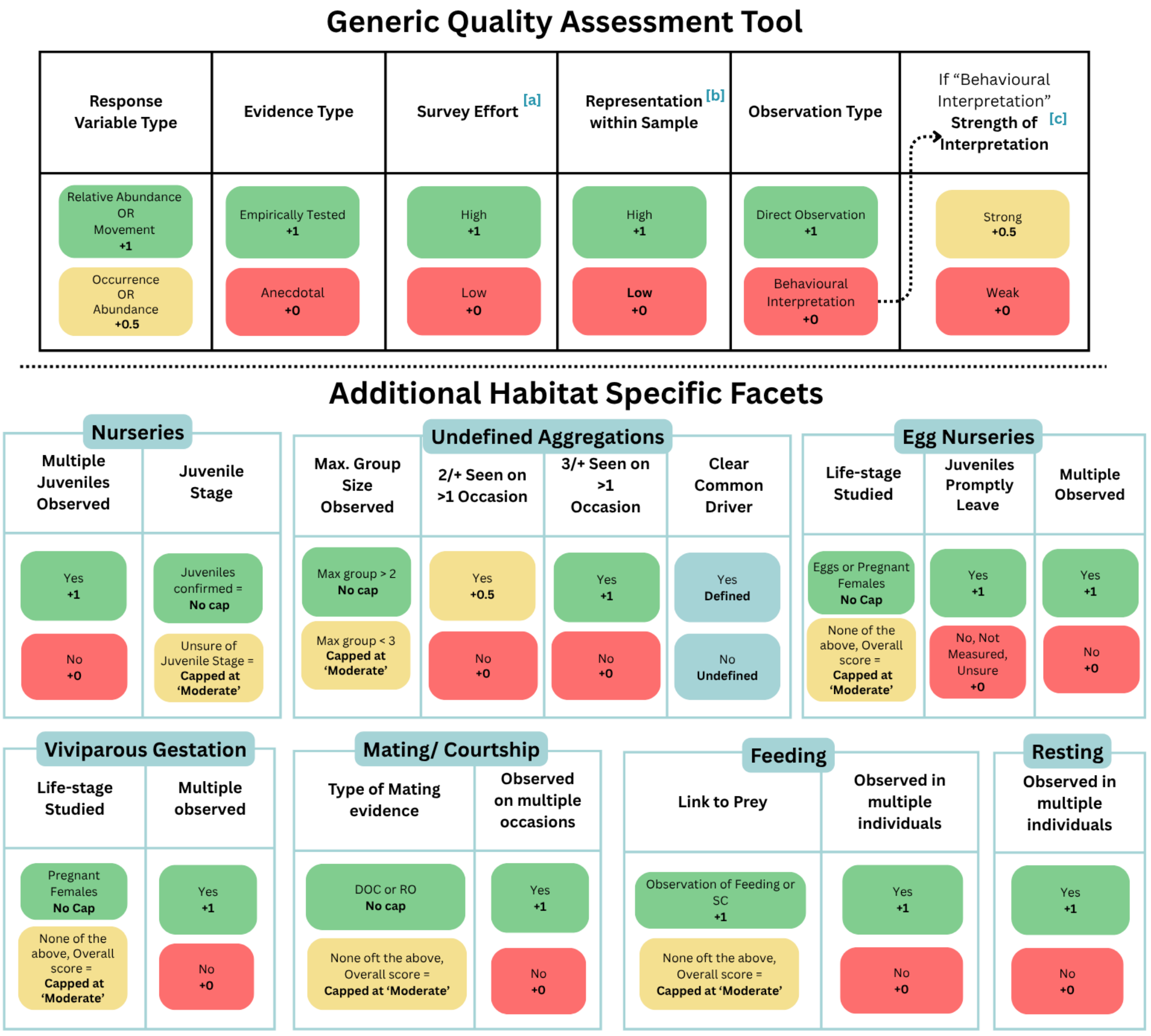


[a] < ~ 30 samples per area or individuals tagged = Low
[b] Behaviour of life stage is represented by < ~ 25 % of sample = Low
[c] Behavioural interpretation is "weak" if the behaviour could be feasibly explained by an alternative driver

**Figure S2. Temporal trend in publications discussing each critical area type.**

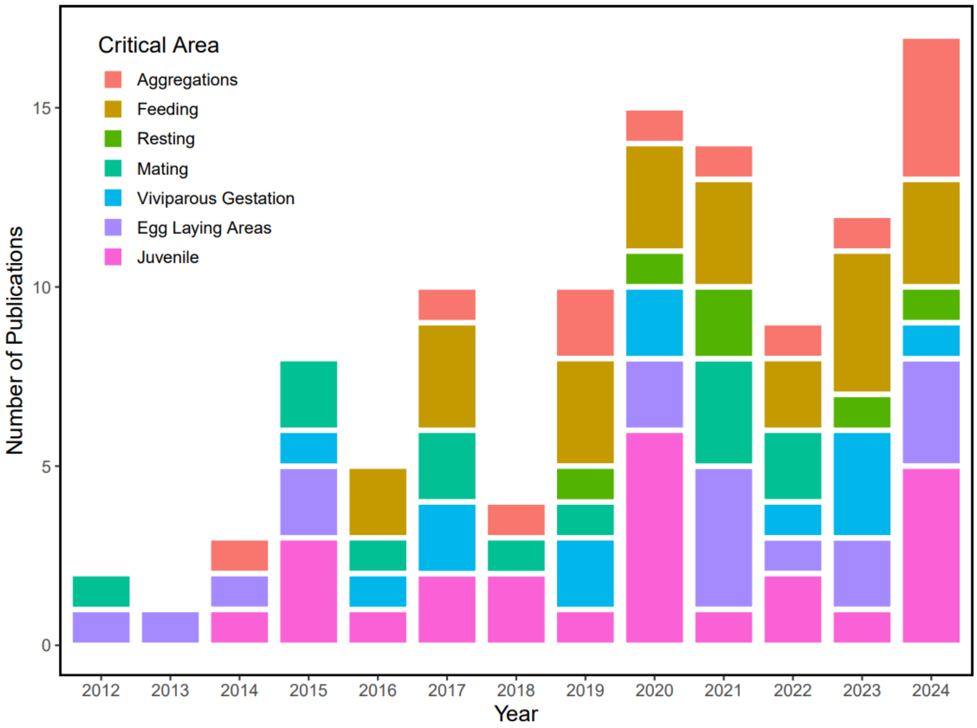
