## Supplementary material for "Tracks, Maps and Gaps: A Testable Research Definition for Critical Chondrichthyan Areas, European Atlantic Insights": Table S8

Table S8. All areas examined and the strength of associated evidence based on the critical area definition requirements (R1-R3) met.

| Reference(s) | Location | Species | R1 | R2 (Species) | R2 (Individual) | R3 (Species) | R3 (Individual) | Strength of Evidence | Quality Assessment | Evidence Cap* | Overall Evidence | Critical Area |
|---|---|---|---|---|---|---|---|---|---|---|---|---|
| **Aggregations** | | | | | | | | | | **Maximum Group Size Observed** | | |
| Thorburn et al., 2024 | Achill Island, Ireland | *Cetorhinus maximus* | ✓ | ✓ | ✓ | ✓ | ✓ | Strong | High | 12 | Strong | Aggregations |
| Biton-Porsmoguer & Lloret, 2023 | The Roadstead of Brest, France | *Mustelus asterias* | ✗ | ✓ | ✗ | ✓ | ✗ | Strong | High | 4 | Strong | Aggregations |
| Kraft et al., 2024b | Professor Luiz Saldanha Marine Park, Portugal | *Dasyatis pastinaca* | ✓ | ✓ | ✓ | ✗ | ✗ | Strong | High | 9 | Strong | Aggregations |
| Gore et al., 2019 | Gunna Sound & Western Coll | *Cetorhinus maximus* | ✓ | ✓ | ✗ | ✗ | ✗ | Strong | High | 12 | Strong | Aggregations |
| Thorburn et al., 2024 | South Barra, Coll & Small Isles, Scotland | *Cetorhinus maximus* | ✗ | ✓ | ✓ | ✓ | ✓ | Strong | High | 2 | Moderate | Aggregations |
| Thorburn et al., 2024 | Malin head, Ireland | *Cetorhinus maximus* | ✗ | ✓ | ✓ | ✗ | ✗ | Moderate | High | 3 | Moderate | Aggregations |
| Lieber et al., 2020 | Isle of Man Waters | *Cetorhinus maximus* | ✗ | ✗ | ✗ | ✓ | ✓ | Moderate | High | 8 | Moderate | Aggregations |
| Lieber et al., 2020 | Irish Waters | *Cetorhinus maximus* | ✗ | ✗ | ✗ | ✓ | ✓ | Moderate | High | 14 | Moderate | Aggregations |
| Lieber et al., 2020 | The Moray Firth | *Cetorhinus maximus* | ✗ | ✗ | ✗ | ✓ | ✓ | Moderate | High | 6 | Moderate | Aggregations |
| Lieber et al., 2020 | West Coast of Scotland | *Cetorhinus maximus* | ✗ | ✗ | ✗ | ✓ | ✓ | Moderate | High | 10 | Moderate | Aggregations |
| Biton-Porsmoguer & Lloret, 2023 | The Trieux Estuary, France | *Mustelus asterias* | ✗ | ✓ | ✗ | ✗ | ✗ | Weak | High | 5 | Weak-Cannot Assess | Aggregations |
| Jung et al., 2024 | Tregor Area, France | *Lamna nasus* | ✗ | ✗ | ✗ | ✗ | ✗ | Weak | Low | Not Reported | Weak-Cannot Assess | Aggregations |
| Thorburn et al., 2024 | Front Between Malin Head & Isle of Islay | *Cetorhinus maximus* | ✗ | ✗ | ✗ | ✗ | ✗ | Weak | High | 3 | Weak-Cannot Assess | Aggregations |
| Lavender et al., 2022 | Off Kerrera and Insh, Loch Sunart to the Sound of Jura MPA, Scotland | *Dipturus intermedius* | ✗ | ✗ | ✗ | ✗ | ✗ | Weak | High | 9 | Weak-Cannot Assess | Aggregations |
| Thorburn et al., 2024 | Tory Island, Ireland | *Cetorhinus maximus* | ✗ | ✗ | ✗ | ✗ | ✗ | Weak | High | 2 | Weak-Cannot Assess | Aggregations |
| Biton-Porsmoguer & Lloret, 2023 | The Thames Estuary, England | *Mustelus asterias* | ✗ | ✗ | ✗ | ✗ | ✗ | Weak | Moderate | 5 | Weak-Cannot Assess | Aggregations |
| Biton-Porsmoguer & Lloret, 2023 | Bristol Channel, England | *Mustelus asterias* | ✗ | ✗ | ✗ | ✗ | ✗ | Weak | Moderate | 3 | Weak-Cannot Assess | Aggregations |
| Biton-Porsmoguer & Lloret, 2023 | South Devon, England | *Mustelus asterias* | ✗ | ✗ | ✗ | ✗ | ✗ | Weak | Moderate | 5 | Weak-Cannot Assess | Aggregations |
| Biton-Porsmoguer & Lloret, 2023 | Normandy, France | *Mustelus asterias* | ✗ | ✗ | ✗ | ✗ | ✗ | Weak | Moderate | 2 | Weak-Cannot Assess | Aggregations |
| Biton-Porsmoguer & Lloret, 2023 | Norfolk coast | *Mustelus asterias* | ✗ | ✗ | ✗ | ✗ | ✗ | Weak | Moderate | 2 | Weak-Cannot Assess | Aggregations |

*This column highlights (in grey) when a cap was placed on the overall score based on the rules outlined in the risk of bias assessment.

| Reference(s) | Location | Species | R1 | R2 (Species) | R2 (Individual) | R3 (Species) | R3 (Individual) | Strength of Evidence | Quality Assessment | Evidence Cap* | Overall Evidence | Critical Area |
|---|---|---|---|---|---|---|---|---|---|---|---|---|
| Biton-Porsmoguer & Lloret, 2023 | Langney Point Eastbourne | *Mustelus asterias* | ✗ | ✗ | ✗ | ✗ | ✗ | Weak | Moderate | 2 | Weak-Cannot Assess | Aggregations |
| Hayes et al., 2018 | West of Shetland, Scotland | *Cetorhinus maximus* | ✓ | ✗ | ✗ | ✗ | ✗ | Weak | High | 3 | Weak-Cannot Assess | Aggregations |
| Serra-Pereira et al., 2014 | Peniche and Cabo da Roca, Portugal | *Rajidae* | ✗ | ✗ | ✗ | ✗ | ✗ | Weak | Low | Definition based on Diversity | Weak-Cannot Assess | Aggregations |
| Meyers et al., 2017 | Waters around the Canary Islands | *Squatina squatina* | ✗ | ✗ | ✗ | ✗ | ✗ | Weak | Moderate | Not Reported | Weak-Cannot Assess | Aggregations |
| Leeb at al., 2021 | Ria de Vigo, Spain | *Raja undulata* | ✗ | ✗ | ✗ | ✗ | ✗ | Weak | Moderate | Not Reported | Weak-Cannot Assess | Aggregations |
| Daban et al., 2024 | Cies Islands, Galicia | *Raja undulata* | ✗ | ✗ | ✗ | ✗ | ✗ | Weak | Moderate | Not Reported | Weak-Cannot Assess | Aggregations |
| Cameron et al., 2019 | Irish Waters | *Lamna nasus* | ✗ | ✗ | ✗ | ✗ | ✗ | Weak | Moderate | Not Reported | Weak-Cannot Assess | Aggregations |
| **Egg Laying Areas** | | | | | | | | | | **Type of Egg laying Evidence** | | |
| Dodd et al., 2022 | Red Rocks and Longay MPA, Scotland | *Dipturus intermedius* | ✓ | ✓ | ✓ | ✓ | ✗ | Strong | High | In-situ Eggs | Strong | Egg Laying Areas |
| Henry et al., 2013 | Mingulay Reef, the Minch, Scotland | *Galeus melastomus* | ✗ | ✓ | ✓ | ✓ | ✗ | Strong | High | In-situ Eggs | Strong | Egg Laying Areas |
| Phillips et al., 2021 | Foot of Shapsinay site, Scotland | *Dipturus intermedius* | ✗ | ✓ | ✓ | ✓ | ✗ | Strong | High | In-situ Eggs | Strong | Egg Laying Areas |
| Phillips et al., 2021 | The Galt, Scotland | *Dipturus intermedius* | ✗ | ✓ | ✓ | ✓ | ✗ | Strong | High | In-situ Eggs | Strong | Egg Laying Areas |
| Schwanck et al., 2024, Thorburn et al., 2021, Thorburn et al., 2023 | Loch Sunart to the Sound of Jura Marine Protected Area, Scotland | *Dipturus intermedius* | ✗ | ✓ | ✓ | ✓ | ✓ | Strong | Moderate | In-situ Eggs/ Pregnant Females/ Mature Females | Strong | Egg Laying Areas |
| Ellis et al., 2024 | North Sea | *Amblyraja radiata* | ✗ | ✓ | ✓ | ✗ | ✗ | Moderate | High | In-situ Eggs | Moderate | Egg Laying Areas |
| Ellis et al., 2024 | Celtic Sea | *Dipturus batis* | ✗ | ✓ | ✓ | ✗ | ✗ | Moderate | High | In-situ Eggs | Moderate | Egg Laying Areas |
| Ellis et al., 2024 | Celtic Sea | *Leucoraja naevus* | ✗ | ✓ | ✓ | ✗ | ✗ | Moderate | High | In-situ Eggs | Moderate | Egg Laying Areas |
| Ellis et al., 2024 | North Sea | *Leucoraja naevus* | ✗ | ✓ | ✓ | ✗ | ✗ | Moderate | High | In-situ Eggs | Moderate | Egg Laying Areas |
| Ellis et al., 2024 | Bristol Channel | *Raja brachyura* | ✗ | ✓ | ✓ | ✗ | ✗ | Moderate | High | In-situ Eggs | Moderate | Egg Laying Areas |
| Ellis et al., 2024 | Coastal waters near Plougrescant, France | *Raja brachyura* | ✗ | ✓ | ✓ | ✗ | ✗ | Moderate | Moderate | In-situ Eggs | Moderate | Egg Laying Areas |
| Ellis et al., 2024 | Bristol Channel | *Raja clavata* | ✗ | ✓ | ✓ | ✗ | ✗ | Moderate | High | In-situ Eggs/ Stranded Eggcases | Moderate | Egg Laying Areas |
| Ellis et al., 2024 | Lyme Bay | *Raja clavata* | ✗ | ✓ | ✓ | ✗ | ✗ | Moderate | High | In-situ Eggs | Moderate | Egg Laying Areas |
| Ellis et al., 2024 | Outer Thames and Netherlands Coast | *Raja clavata* | ✗ | ✓ | ✓ | ✗ | ✗ | Moderate | High | In-situ Eggs | Moderate | Egg Laying Areas |
| Ellis et al., 2024 | The Wash | *Raja clavata* | ✗ | ✓ | ✓ | ✗ | ✗ | Moderate | High | In-situ Eggs | Moderate | Egg Laying Areas |

*This column highlights (in grey) when a cap was placed on the overall score based on the rules outlined in the risk of bias assessment.

| Reference(s) | Location | Species | R1 | R2 (Species) | R2 (Individual) | R3 (Species) | R3 (Individual) | Strength of Evidence | Quality Assessment | Evidence Cap* | Overall Evidence | Critical Area |
|---|---|---|---|---|---|---|---|---|---|---|---|---|
| Ellis et al., 2024 | Bristol Channel | *Raja montagui* | ✗ | ✓ | ✓ | ✗ | ✗ | Moderate | High | In-situ Eggs | Moderate | Egg Laying Areas |
| Ellis et al., 2024 | Netherlands Coast/ Eastern English Channel. | *Raja montagui* | ✗ | ✓ | ✓ | ✗ | ✗ | Moderate | Moderate | In-situ Eggs | Moderate | Egg Laying Areas |
| Ellis et al., 2024 | North Cornwall | *Raja montagui* | ✗ | ✓ | ✓ | ✗ | ✗ | Moderate | Moderate | In-situ Eggs | Moderate | Egg Laying Areas |
| Ellis et al., 2024 | South Eastern Coasts of Ireland | *Raja montagui* | ✗ | ✓ | ✓ | ✗ | ✗ | Moderate | High | In-situ Eggs | Moderate | Egg Laying Areas |
| Ellis et al., 2024 | The Wash | *Raja montagui* | ✗ | ✓ | ✓ | ✗ | ✗ | Moderate | Moderate | In-situ Eggs | Moderate | Egg Laying Areas |
| Ellis et al., 2024 | Western English Channel | *Raja montagui* | ✗ | ✓ | ✓ | ✗ | ✗ | Moderate | Moderate | In-situ Eggs | Moderate | Egg Laying Areas |
| Ellis et al., 2024 | Flamanville coast, France | *Raja undulata* | ✗ | ✓ | ✓ | ✗ | ✗ | Moderate | Moderate | In-situ Eggs | Moderate | Egg Laying Areas |
| Henry et al., 2013 | Banana Reef, the Minch, Scotland | *Galeus melastomus* | ✗ | ✓ | ✓ | ✗ | ✗ | Moderate | High | In-situ Eggs | Moderate | Egg Laying Areas |
| Maia et al., 2012 | Inshore Portugese Waters | *Leucoraja naevus* | ✗ | ✓ | ✓ | ✗ | ✗ | Moderate | High | In-situ Eggs | Moderate | Egg Laying Areas |
| McAllister at al., 2024 | NE Shetland Islands , Scotland | *Raja clavata* | ✓ | ✗ | ✗ | ✓ | ✗ | Strong | High | Mature Females | Moderate | Egg Laying Areas |
| Phillips et al., 2021 | The Grinds, Scotland | *Dipturus intermedius* | ✗ | ✓ | ✓ | ✗ | ✗ | Moderate | Moderate | In-situ Eggs | Moderate | Egg Laying Areas |
| Serra-Pereira et al., 2015 | Portugal, Coastal Waters | *Raja undulata* | ✗ | ✓ | ✓ | ✗ | ✗ | Moderate | High | Pregnant Females/ In-situ Eggs | Moderate | Egg Laying Areas |
| Bird et al., 2020 | Western Scotland & Irish Sea | *Raja brachyura* | ✗ | ✗ | ✗ | ✗ | ✗ | Weak | High | Mature Adults | Weak-Cannot Assess | Egg Laying Areas |
| Biton-Porsmoguer et al., 2020 | Littoral seino-marin, France | *Raja clavata* | ✗ | ✗ | ✗ | ✗ | ✗ | Weak | Moderate | Pregnant Females | Weak-Cannot Assess | Egg Laying Areas |
| Biton-Porsmoguer et al., 2020 | Boulogne-sur-Mer and Dieppe, France | *Raja montagui* | ✗ | ✗ | ✗ | ✗ | ✗ | Weak | Moderate | Pregnant Females | Weak-Cannot Assess | Egg Laying Areas |
| Biton-Porsmoguer et al., 2020 | East of Cherbourg, France | *Raja montagui* | ✗ | ✗ | ✗ | ✗ | ✗ | Weak | Moderate | Pregnant Females | Weak-Cannot Assess | Egg Laying Areas |
| Biton-Porsmoguer et al., 2020 | Littoral seino-marin, France | *Raja montagui* | ✗ | ✗ | ✗ | ✗ | ✗ | Weak | Moderate | Pregnant Females | Weak-Cannot Assess | Egg Laying Areas |
| Ellis et al., 2024 | Eastern Seaboard of Scotland | *Amblyraja radiata* | ✗ | ✗ | ✗ | ✗ | ✗ | Weak | High | Stranded Eggcases | Weak-Cannot Assess | Egg Laying Areas |
| Ellis et al., 2024 | Eastern Seaboard of England | *Amblyraja radiata* | ✗ | ✗ | ✗ | ✗ | ✗ | Weak | High | Stranded Eggcases | Weak-Cannot Assess | Egg Laying Areas |
| Ellis et al., 2024 | Cornwall | *Dipturus batis* | ✗ | ✗ | ✗ | ✗ | ✗ | Weak | High | Stranded Eggcases | Weak-Cannot Assess | Egg Laying Areas |
| Ellis et al., 2024 | Mid Wales | *Dipturus batis* | ✗ | ✗ | ✗ | ✗ | ✗ | Weak | Moderate | Stranded Eggcases | Weak-Cannot Assess | Egg Laying Areas |
| Ellis et al., 2024 | Northern Ireland | *Dipturus batis* | ✗ | ✗ | ✗ | ✗ | ✗ | Weak | High | Stranded Eggcases | Weak-Cannot Assess | Egg Laying Areas |
| Ellis et al., 2024 | Orkney | *Dipturus batis* | ✗ | ✗ | ✗ | ✗ | ✗ | Weak | High | Stranded Eggcases | Weak-Cannot Assess | Egg Laying Areas |
| Ellis et al., 2024 | South Wales | *Dipturus batis* | ✗ | ✗ | ✗ | ✗ | ✗ | Weak | High | Stranded Eggcases | Weak-Cannot Assess | Egg Laying Areas |

*This column highlights (in grey) when a cap was placed on the overall score based on the rules outlined in the risk of bias assessment.

| Reference(s) | Location | Species | R1 | R2 (Species) | R2 (Individual) | R3 (Species) | R3 (Individual) | Strength of Evidence | Quality Assessment | Evidence Cap* | Overall Evidence | Critical Area |
|---|---|---|---|---|---|---|---|---|---|---|---|---|
| Ellis et al., 2024 | The Highlands | *Dipturus batis* | ✗ | ✗ | ✗ | ✗ | ✗ | Weak | Moderate | Stranded Eggcases | Weak-Cannot Assess | Egg Laying Areas |
| Ellis et al., 2024 | Cardigan Bay | *Dipturus intermedius* | ✗ | ✗ | ✗ | ✗ | ✗ | Weak | Moderate | Stranded Eggcases | Weak-Cannot Assess | Egg Laying Areas |
| Ellis et al., 2024 | Northen Scotland | *Dipturus intermedius* | ✗ | ✗ | ✗ | ✗ | ✗ | Weak | High | Stranded Eggcases | Weak-Cannot Assess | Egg Laying Areas |
| Ellis et al., 2024 | Northumberland | *Dipturus intermedius* | ✗ | ✗ | ✗ | ✗ | ✗ | Weak | Moderate | Stranded Eggcases | Weak-Cannot Assess | Egg Laying Areas |
| Ellis et al., 2024 | Orkney | *Dipturus intermedius* | ✗ | ✗ | ✗ | ✗ | ✗ | Weak | High | Stranded Eggcases | Weak-Cannot Assess | Egg Laying Areas |
| Ellis et al., 2024 | West Coast of Ireland | *Dipturus intermedius* | ✗ | ✗ | ✗ | ✗ | ✗ | Weak | High | Stranded Eggcases | Weak-Cannot Assess | Egg Laying Areas |
| Ellis et al., 2024 | Celtic Sea | *Leucoraja fullonica* | ✗ | ✗ | ✗ | ✗ | ✗ | Weak | Moderate | Pregnant Female | Weak-Cannot Assess | Egg Laying Areas |
| Ellis et al., 2024 | Eastern Irish Sea | *Leucoraja naevus* | ✗ | ✗ | ✗ | ✗ | ✗ | Weak | High | Stranded Eggcases | Weak-Cannot Assess | Egg Laying Areas |
| Ellis et al., 2024 | Northen Scotland | *Leucoraja naevus* | ✗ | ✗ | ✗ | ✗ | ✗ | Weak | High | Stranded Eggcases | Weak-Cannot Assess | Egg Laying Areas |
| Ellis et al., 2024 | Southwest England | *Leucoraja naevus* | ✗ | ✗ | ✗ | ✗ | ✗ | Weak | High | Stranded Eggcases | Weak-Cannot Assess | Egg Laying Areas |
| Ellis et al., 2024 | Ireland | *Raja brachyura* | ✗ | ✗ | ✗ | ✗ | ✗ | Weak | High | Stranded Eggcases | Weak-Cannot Assess | Egg Laying Areas |
| Ellis et al., 2024 | Northern Ireland | *Raja brachyura* | ✗ | ✗ | ✗ | ✗ | ✗ | Weak | High | Stranded Eggcases | Weak-Cannot Assess | Egg Laying Areas |
| Ellis et al., 2024 | Scotland | *Raja brachyura* | ✗ | ✗ | ✗ | ✗ | ✗ | Weak | High | Stranded Eggcases | Weak-Cannot Assess | Egg Laying Areas |
| Ellis et al., 2024 | Southern Coasts of England | *Raja brachyura* | ✗ | ✗ | ✗ | ✗ | ✗ | Weak | High | Stranded Eggcases | Weak-Cannot Assess | Egg Laying Areas |
| Ellis et al., 2024 | Southern Coasts of Wales | *Raja brachyura* | ✗ | ✗ | ✗ | ✗ | ✗ | Weak | High | Stranded Eggcases | Weak-Cannot Assess | Egg Laying Areas |
| Ellis et al., 2024 | Areas bordering the Irish Sea | *Raja clavata* | ✗ | ✗ | ✗ | ✗ | ✗ | Weak | High | Stranded Eggcases | Weak-Cannot Assess | Egg Laying Areas |
| Ellis et al., 2024 | Southern & Eastern Coasts of England | *Raja clavata* | ✗ | ✗ | ✗ | ✗ | ✗ | Weak | High | Stranded Eggcases | Weak-Cannot Assess | Egg Laying Areas |
| Ellis et al., 2024 | Dumfries & Galloway | *Raja microocellata* | ✗ | ✗ | ✗ | ✗ | ✗ | Weak | High | Stranded Eggcases | Weak-Cannot Assess | Egg Laying Areas |
| Ellis et al., 2024 | South Eastern Coasts of Ireland | *Raja microocellata* | ✗ | ✗ | ✗ | ✗ | ✗ | Weak | High | Stranded Eggcases | Weak-Cannot Assess | Egg Laying Area |
| Ellis et al., 2024 | Southern & Western Coasts of England. | *Raja microocellata* | ✗ | ✗ | ✗ | ✗ | ✗ | Weak | High | Stranded Eggcases | Weak-Cannot Assess | Egg Laying Areas |
| Ellis et al., 2024 | Welsh Coasts | *Raja microocellata* | ✗ | ✗ | ✗ | ✗ | ✗ | Weak | High | Stranded Eggcases | Weak-Cannot Assess | Egg Laying Areas |

*This column highlights (in grey) when a cap was placed on the overall score based on the rules outlined in the risk of bias assessment.

| Reference(s) | Location | Species | R1 | R2 (Species) | R2 (Individual) | R3 (Species) | R3 (Individual) | Strength of Evidence | Quality Assessment | Evidence Cap* | Overall Evidence | Critical Area |
|---|---|---|---|---|---|---|---|---|---|---|---|---|
| Ellis et al., 2024 | Yorkshire | *Raja microocellata* | ✗ | ✗ | ✗ | ✗ | ✗ | Weak | High | Stranded Eggcases | Weak-Cannot Assess | Egg Laying Areas |
| Ellis et al., 2024 | UK coastline (excluding NE England & Scotland) | *Raja montagui* | ✗ | ✗ | ✗ | ✗ | ✗ | Weak | High | Stranded Eggcases | Weak-Cannot Assess | Egg Laying Areas |
| Ellis et al., 2024 | Southern Coast of England & Channel Islands | *Raja undulata* | ✗ | ✗ | ✗ | ✗ | ✗ | Weak | High | Stranded Eggcases | Weak-Cannot Assess | Egg Laying Areas |
| Ellis et al., 2024 | Southern North Sea | *Raja undulata* | ✗ | ✗ | ✗ | ✗ | ✗ | Weak | High | Stranded Eggcases | Weak-Cannot Assess | Egg Laying Areas |
| Ellis et al., 2024 | Western Ireland | *Raja undulata* | ✗ | ✗ | ✗ | ✗ | ✗ | Weak | High | Stranded Eggcases | Weak-Cannot Assess | Egg Laying Areas |
| Ellis et al., 2024 | Western Wales | *Raja undulata* | ✗ | ✗ | ✗ | ✗ | ✗ | Weak | High | Stranded Eggcases | Weak-Cannot Assess | Egg Laying Areas |
| Ellis et al., 2024 | Braunton Burrows, North Devon | *Rostroraja alba* | ✗ | ✗ | ✗ | ✗ | ✗ | Weak | High | Stranded Eggcases | Weak-Cannot Assess | Egg Laying Areas |
| Ellis et al., 2024 | Fair Isle, SW Shetland | *Rostroraja alba* | ✗ | ✗ | ✗ | ✗ | ✗ | Weak | High | Stranded Eggcases | Weak-Cannot Assess | Egg Laying Areas |
| Ellis et al., 2024 | Gunwalloe, SW Cornwall | *Rostroraja alba* | ✗ | ✗ | ✗ | ✗ | ✗ | Weak | High | Stranded Eggcases | Weak-Cannot Assess | Egg Laying Areas |
| Ellis et al., 2024 | Northwest Wales | *Rostroraja alba* | ✗ | ✗ | ✗ | ✗ | ✗ | Weak | High | Stranded Eggcases | Weak-Cannot Assess | Egg Laying Areas |
| Ellis et al., 2024 | Peterhead, NE Scotland | *Rostroraja alba* | ✗ | ✗ | ✗ | ✗ | ✗ | Weak | High | Stranded Eggcases | Weak-Cannot Assess | Egg Laying Areas |
| Maia et al., 2015 | Peniche region, Portugal | *Leucoraja naevus* | ✗ | ✗ | ✗ | ✗ | ✗ | Weak | High | In-situ Eggs | Weak-Cannot Assess | Egg Laying Areas |
| Maia et al., 2015 | Peniche region, Portugal | *Raja brachyura* | ✗ | ✗ | ✗ | ✗ | ✗ | Weak | High | In-situ Eggs | Weak-Cannot Assess | Egg Laying Areas |
| Maia et al., 2015 | Peniche region, Portugal | *Raja clavata* | ✗ | ✗ | ✗ | ✗ | ✗ | Weak | High | In-situ Eggs | Weak-Cannot Assess | Egg Laying Areas |
| Maia et al., 2015 | Peniche region, Portugal | *Raja microocellata* | ✗ | ✗ | ✗ | ✗ | ✗ | Weak | Moderate | In-situ Eggs | Weak-Cannot Assess | Egg Laying Areas |
| Maia et al., 2015 | Peniche region, Portugal | *Raja montagui* | ✗ | ✗ | ✗ | ✗ | ✗ | Weak | High | In-situ Eggs | Weak-Cannot Assess | Egg Laying Areas |
| Maia et al., 2015 | Peniche region, Portugal | *Raja undulata* | ✗ | ✗ | ✗ | ✗ | ✗ | Weak | High | In-situ Eggs | Weak-Cannot Assess | Egg Laying Areas |
| Papadopoulo et al., 2023 | Inshore waters near Cies Island, Spain | *Raja clavata* | ✗ | ✗ | ✗ | ✗ | ✗ | Weak | Moderate | Mature Adults | Weak-Cannot Assess | Egg Laying Areas |
| Santos et al., 2021 | Azores | *Raja clavata* | ✗ | ✗ | ✗ | ✗ | ✗ | Weak | High | Reproductively Active Females | Weak-Cannot Assess | Egg Laying Area |
| Serra-Pereira et al., 2014 | Mar da Ericeira, Portugal | *Leucoraja naevus* | ✗ | ✗ | ✗ | ✗ | ✗ | Weak | High | Reproductively Active Adults | Weak-Cannot Assess | Egg Laying Areas |
| Serra-Pereira et al., 2014 | Areia Branca, Portugal | *Raja brachyura* | ✗ | ✗ | ✗ | ✗ | ✗ | Weak | High | Reproductively Active Adults | Weak-Cannot Assess | Egg Laying Areas |

*This column highlights (in grey) when a cap was placed on the overall score based on the rules outlined in the risk of bias assessment.

| Reference(s) | Location | Species | R1 | R2 (Species) | R2 (Individual) | R3 (Species) | R3 (Individual) | Strength of Evidence | Quality Assessment | Evidence Cap* | Overall Evidence | Critical Area |
|---|---|---|---|---|---|---|---|---|---|---|---|---|
| Serra-Pereira et al., 2014 | Mar do Cachimbo, Portugal | *Raja brachyura* | ✗ | ✗ | ✗ | ✗ | ✗ | Weak | High | Reproductively Active Adults | Weak-Cannot Assess | Egg Laying Areas |
| Serra-Pereira et al., 2014 | Santa Cruz, Portugal | *Raja brachyura* | ✗ | ✗ | ✗ | ✗ | ✗ | Weak | High | Reproductively Active Adults | Weak-Cannot Assess | Egg Laying Areas |
| Serra-Pereira et al., 2014 | Mar do NW da Roca, Portugal | *Raja clavata* | ✗ | ✗ | ✗ | ✗ | ✗ | Weak | High | Reproductively Active Adults | Weak-Cannot Assess | Egg Laying Areas |
| Serra-Pereira et al., 2014 | Santa Cruz, Portugal | *Raja clavata* | ✗ | ✗ | ✗ | ✗ | ✗ | Weak | High | Reproductively Active Adults | Weak-Cannot Assess | Egg Laying Areas |
| Serra-Pereira et al., 2014 | Berlengas, Portugal | *Raja microocellata* | ✗ | ✗ | ✗ | ✗ | ✗ | Weak | High | Reproductively Active Adults | Weak-Cannot Assess | Egg Laying Areas |
| Simpson et al., 2021 | Coastal waters off Mothecombe & Bigbury-on-sea (Southwest UK) | *Raja brachyura* | ✓ | ✗ | ✗ | ✗ | ✗ | Weak | Low | Mature Adults | Weak-Cannot Assess | Egg Laying Areas |
| **Feeding** | | | | | | | | | | **Type of Feeding Evidence.** | | |
| Gore et al., 2019, Gore et al., 2016 | Inner hebrides, Scotland | *Cetorhinus maximus* | ✗ | ✓ | ✓ | ✓ | ✓ | Strong | High | Feeding Observation/ Surface Swimming | Strong | Feeding |
| Couto et al., 2017 | Southern Portugal | *Cetorhinus maximus* | ✗ | ✓ | ✗ | ✓ | ✗ | Strong | High | Predator-Prey Overlap/ Surface Swimming | Strong | Feeding |
| Kraft et al., 2023 | Sado Estuary, Portugal | *Dasyatis pastinaca* | ✗ | ✗ | ✗ | ✓ | ✓ | Moderate | High | Migration | Moderate | Feeding |
| Biais et al., 2017 | Coastal shelf, Bay of Biscay | *Lamna nasus* | ✗ | ✓ | ✓ | ✓ | ✓ | Strong | High | Predator-Prey Overlap | Moderate | Feeding |
| Kraft et al., 2024a, Sousa et al., 2019 | Professor Luiz Saldanha Marine Park, Portugal | *Rostroraja alba* | ✗ | ✓ | ✓ | ✓ | ✗ | Strong | High | Activity Levels | Moderate | Feeding |
| Rudd et al., 2021 | Coll & Tiree Waters, Scotland | *Cetorhinus maximus* | ✗ | ✗ | ✗ | ✗ | ✗ | Weak | High | Feeding Observation | Weak-Cannot Assess | Feeding |
| Doherty et al., 2017a | Sea of the Hebrides MPA, Scotland | *Cetorhinus maximus* | ✗ | ✗ | ✗ | ✗ | ✗ | Weak | High | Feeding Observation | Weak-Cannot Assess | Feeding |
| Ponte et al., 2016, Fontaine et al., 2023 | Terceira Island, Azores | *Dasyatis pastinaca* | ✗ | ✗ | ✗ | ✗ | ✗ | Weak | Moderate | Stomach Contents | Weak-Cannot Assess | Feeding |
| Brown-Vuillemin et al., 2020 | Celtic Sea | *Dipturus batis* | ✗ | ✗ | ✗ | ✓ | ✗ | Weak | High | Stomach Contents | Weak-Cannot Assess | Feeding |
| Thorburn et al., 2021 | Deep trench system, Loch Sunart to the Sound of Jura MPA, Scotland | *Dipturus intermedius* | ✗ | ✗ | ✗ | ✗ | ✗ | Weak | High | Feeding Observation | Weak-Cannot Assess | Feeding |
| Rodriguez-Garcia al., 2024 | Gulf of Cadiz, Spain | *Galeus melastomus* | ✗ | ✗ | ✗ | ✗ | ✗ | Weak | High | Stomach Contents | Weak-Cannot Assess | Feeding |
| Carrasco-Puig et al., 2024 | Bay of biscay (NW Spain) Asturias, Spain | *Lamna nasus* | ✗ | ✗ | ✗ | ✗ | ✗ | Weak | Low | No Clear Evidence | Weak-Cannot Assess | Feeding |
| Cameron et al., 2019 | Northward movement from Irish waters | *Lamna nasus* | ✗ | ✗ | ✗ | ✗ | ✗ | Weak | High | Predator-Prey Overlap | Weak-Cannot Assess | Feeding |

*This column highlights (in grey) when a cap was placed on the overall score based on the rules outlined in the risk of bias assessment.

| Reference(s) | Location | Species | R1 | R2 (Species) | R2 (Individual) | R3 (Species) | R3 (Individual) | Strength of Evidence | Quality Assessment | Evidence Cap* | Overall Evidence | Critical Area |
|---|---|---|---|---|---|---|---|---|---|---|---|---|
| Biton-Porsmoguer et al., 2022 | Bay of the Seine, France | *Mustelus asterias* | ✗ | ✓ | ✗ | ✗ | ✗ | Weak | High | Stomach Contents | Weak-Cannot Assess | Feeding |
| Simpson et al., 2021 | Plymouth coastal waters | *Raja brachyura* | ✗ | ✗ | ✗ | ✗ | ✗ | Weak | Moderate | Habitat Preference | Weak-Cannot Assess | Feeding |
| Biton-Porsmoguer et al., 2020 | Littoral seinomarin | *Raja clavata* | ✓ | ✗ | ✗ | ✗ | ✗ | Weak | Low | Stomach Contents | Weak-Cannot Assess | Feeding |
| Biton-Porsmoguer et al., 2020 | Bay of the Seine, France | *Raja clavata* | ✓ | ✗ | ✗ | ✗ | ✗ | Weak | Low | Stomach Contents | Weak-Cannot Assess | Feeding |
| Biton-Porsmoguer et al., 2020 | Bay of the Somme, France | *Raja clavata* | ✓ | ✗ | ✗ | ✗ | ✗ | Weak | Low | Stomach Contents | Weak-Cannot Assess | Feeding |
| Biton-Porsmoguer et al., 2020 | Normand-Breton Gulf | *Raja clavata* | ✓ | ✗ | ✗ | ✗ | ✗ | Weak | Low | Stomach Contents | Weak-Cannot Assess | Feeding |
| Simpson et al., 2021 | Plymouth coastal waters | *Raja clavata* | ✗ | ✗ | ✗ | ✗ | ✗ | Weak | Moderate | Habitat Preference | Weak-Cannot Assess | Feeding |
| Fontaine et al., 2023 | Terceira Island, Azores | *Raja clavata* | ✗ | ✗ | ✗ | ✗ | ✗ | Weak | High | Stomach Contents | Weak-Cannot Assess | Feeding |
| Simpson et al., 2021 | Plymouth coastal waters | *Raja microocellata* | ✗ | ✗ | ✗ | ✗ | ✗ | Weak | Moderate | Habitat Preference | Weak-Cannot Assess | Feeding |
| Biton-Porsmoguer et al., 2020 | Littoral seinomarin | *Raja montagui* | ✓ | ✗ | ✗ | ✗ | ✗ | Weak | Low | Stomach Contents | Weak-Cannot Assess | Feeding |
| Biton-Porsmoguer et al., 2020 | Bay of the Seine, France | *Raja montagui* | ✓ | ✗ | ✗ | ✗ | ✗ | Weak | Low | Stomach Contents | Weak-Cannot Assess | Feeding |
| Biton-Porsmoguer et al., 2020 | Bay of the Somme, France | *Raja montagui* | ✓ | ✗ | ✗ | ✗ | ✗ | Weak | Low | Stomach Contents | Weak-Cannot Assess | Feeding |
| Biton-Porsmoguer et al., 2020 | Normand-Breton Gulf | *Raja montagui* | ✓ | ✗ | ✗ | ✗ | ✗ | Weak | Low | Stomach Contents | Weak-Cannot Assess | Feeding |
| Simpson et al., 2021 | Plymouth coastal waters | *Raja montagui* | ✗ | ✗ | ✗ | ✗ | ✗ | Weak | Moderate | Habitat Preference | Weak-Cannot Assess | Feeding |
| Barker et al., 2022 | Cardigan bay, Wales | *Squatina squatina* | ✗ | ✗ | ✗ | ✗ | ✗ | Weak | High | Predator-Prey Overlap | Weak-Cannot Assess | Feeding |
| Mead et al., 2023 | La Graciosa Marine Reserve, Canary Islands | *Squatina squatina* | ✗ | ✗ | ✗ | ✗ | ✗ | Weak | Moderate | Activity Levels | Weak-Cannot Assess | Feeding |
| Tuya et al., 2020 | Las Canteras Beach, Gran Canaria | *Squatina squatina* | ✗ | ✗ | ✗ | ✗ | ✗ | Weak | Moderate | Abundance | Weak-Cannot Assess | Feeding |
| Sousa et al., 2019 | Professor Luiz Saldanha Marine Park, Portugal | *Rostroraja alba* | ✗ | ✗ | ✗ | ✗ | ✗ | Weak | Low | Activity Levels | Weak-Cannot Assess | Feeding |
| **Immature Areas** | | | | | | | | | | **Immature Life Stage** | | |
| Ellis et al., 2024 | Between the Dogger Bank and Norwegian Deeps | *Amblyraja radiata* | ✓ | ✗ | ✗ | ✓ | ✗ | Strong | High | 1st Cohort | Strong | Immature Areas |
| Ellis et al., 2024 | Due East of Shetland | *Amblyraja radiata* | ✓ | ✗ | ✗ | ✓ | ✗ | Strong | High | 1st Cohort | Strong | Immature Areas |
| Cameron et al., 2019 | Irish waters | *Lamna nasus* | ✗ | ✓ | ✗ | ✓ | ✓ | Strong | High | Neonate | Strong | Immature Areas |

*This column highlights (in grey) when a cap was placed on the overall score based on the rules outlined in the risk of bias assessment.

| Reference(s) | Location | Species | R1 | R2 (Species) | R2 (Individual) | R3 (Species) | R3 (Individual) | Strength of Evidence | Quality Assessment | Evidence Cap* | Overall Evidence | Critical Area |
|---|---|---|---|---|---|---|---|---|---|---|---|---|
| Jung et al., 2024 | Tregor Area, France | *Lamna nasus* | ✗ | ✓ | ✓ | ✓ | ✗ | Strong | High | Juveniles | Strong | Immature Areas |
| Ellis et al., 2024 | Central Irish Sea & St George's channel | *Leucoraja naevus* | ✓ | ✗ | ✗ | ✓ | ✗ | Strong | High | 1st Cohort | Strong | Immature Areas |
| Ellis et al., 2024 | Northern North Sea | *Leucoraja naevus* | ✓ | ✗ | ✗ | ✓ | ✗ | Strong | High | Juveniles | Strong | Immature Areas |
| Ellis et al., 2024 | Between Dublin Bay and Wicklow, Irish Sea | *Raja brachyura* | ✓ | ✗ | ✗ | ✓ | ✗ | Strong | High | 1st Cohort | Strong | Immature Areas |
| Ellis et al., 2024 | Cardigan Bay, Irish Sea | *Raja brachyura* | ✓ | ✗ | ✗ | ✓ | ✗ | Strong | High | Juveniles | Strong | Immature Areas |
| Ellis et al., 2024 | East of Cherbourg Peninsula, France | *Raja brachyura* | ✓ | ✗ | ✗ | ✓ | ✗ | Strong | High | Juveniles | Strong | Immature Areas |
| Ellis et al., 2024 | Luce Bay, Irish Sea | *Raja brachyura* | ✓ | ✗ | ✗ | ✓ | ✗ | Strong | High | 1st Cohort | Strong | Immature Areas |
| Ellis et al., 2024 | South wales, Bristol Channel | *Raja brachyura* | ✓ | ✗ | ✗ | ✓ | ✗ | Strong | High | 1st Cohort | Strong | Immature Areas |
| Ellis et al., 2024 | Southeastern English Coast (Incl. due East of Kent) | *Raja brachyura* | ✓ | ✗ | ✗ | ✓ | ✗ | Strong | High | 1st Cohort | Strong | Immature Areas |
| Ellis et al., 2024 | Baie de Seine, Eastern English Channel | *Raja clavata* | ✓ | ✗ | ✗ | ✓ | ✗ | Strong | High | 1st Cohort | Strong | Immature Areas |
| Ellis et al., 2024 | Baie de Somme, France | *Raja clavata* | ✓ | ✗ | ✗ | ✓ | ✗ | Strong | High | 1st Cohort | Strong | Immature Areas |
| Ellis et al., 2024 | Bournemouth Bay, Eastern English Channel | *Raja clavata* | ✓ | ✗ | ✗ | ✓ | ✗ | Strong | High | 1st Cohort | Strong | Immature Areas |
| Ellis et al., 2024 | Cardigan Bay, Irish Sea | *Raja clavata* | ✓ | ✗ | ✗ | ✓ | ✗ | Strong | High | 1st Cohort | Strong | Immature Areas |
| Ellis et al., 2024 | Carmarthen Bay, Bristol Channel | *Raja clavata* | ✓ | ✗ | ✗ | ✓ | ✗ | Strong | High | 1st Cohort | Strong | Immature Areas |
| Ellis et al., 2024 | Liverpool Bay, Irish Sea | *Raja clavata* | ✓ | ✗ | ✗ | ✓ | ✗ | Strong | High | 1st Cohort | Strong | Immature Areas |
| Ellis et al., 2024 | Off Brighton, Eastern English Channel | *Raja clavata* | ✓ | ✗ | ✗ | ✓ | ✗ | Strong | High | 1st Cohort | Strong | Immature Areas |
| Ellis et al., 2024 | Outer Thames, Southern North Sea | *Raja clavata* | ✓ | ✗ | ✗ | ✓ | ✗ | Strong | High | 1st Cohort | Strong | Immature Areas |
| Ellis et al., 2024 | Pevensey Bay, Eastern English Channel | *Raja clavata* | ✓ | ✗ | ✗ | ✓ | ✗ | Strong | High | 1st Cohort | Strong | Immature Areas |
| McAllister at al., 2024 | Shallow waters, Shetland Islands , Scotland | *Raja clavata* | ✓ | ✗ | ✗ | ✓ | ✗ | Strong | High | Immature | Strong | Immature Areas |
| Ellis et al., 2024 | Solway Firth, Irish Sea | *Raja clavata* | ✓ | ✗ | ✗ | ✓ | ✗ | Strong | High | 1st Cohort | Strong | Immature Areas |
| Ellis et al., 2024 | Swansea Bay, Bristol channel | *Raja clavata* | ✓ | ✗ | ✗ | ✓ | ✗ | Strong | High | 1st Cohort | Strong | Immature Areas |
| Ellis et al., 2024 | Carmarthen Bay, Bristol Channel | *Raja microocellata* | ✓ | ✗ | ✗ | ✓ | ✗ | Strong | High | Juveniles | Strong | Immature Areas |
| Ellis et al., 2024 | Bristol Channel | *Raja montagui* | ✓ | ✗ | ✗ | ✓ | ✗ | Strong | High | 1st Cohort | Strong | Immature Areas |
| Ellis et al., 2024 | Caernarfon Bay, Irish Sea | *Raja montagui* | ✓ | ✗ | ✗ | ✓ | ✗ | Strong | High | 1st Cohort | Strong | Immature Areas |
| Ellis et al., 2024 | Cardigan Bay, Irish Sea | *Raja montagui* | ✓ | ✗ | ✗ | ✓ | ✗ | Strong | High | 1st Cohort | Strong | Immature Areas |

*This column highlights (in grey) when a cap was placed on the overall score based on the rules outlined in the risk of bias assessment.

| Reference(s) | Location | Species | R1 | R2 (Species) | R2 (Individual) | R3 (Species) | R3 (Individual) | Strength of Evidence | Quality Assessment | Evidence Cap* | Overall Evidence | Critical Area |
|---|---|---|---|---|---|---|---|---|---|---|---|---|
| Ellis et al., 2024 | near Dublin Bay, Irish Sea | *Raja montagui* | ✓ | ✗ | ✗ | ✓ | ✗ | Strong | High | 1st Cohort | Strong | Immature Areas |
| Ellis et al., 2024 | Baie de Somme, France | *Raja undulata* | ✓ | ✗ | ✗ | ✓ | ✗ | Strong | High | Juveniles | Strong | Immature Areas |
| Ellis et al., 2024 | East of Cherbourg Peninsula, France | *Raja undulata* | ✓ | ✗ | ✗ | ✓ | ✗ | Strong | High | Juveniles | Strong | Immature Areas |
| Kraft et al., 2024a | Professor Luiz Saldanha Marine Park, Portugal | *Rostroraja alba* | ✗ | ✓ | ✓ | ✓ | ✓ | Strong | High | Juveniles | Strong | Immature Areas |
| Jimenez-Alvarado et al., 2020 | Playa Chica, Lanzarote | *Squatina squatina* | ✓ | ✓ | ✗ | ✓ | ✗ | Strong | Moderate | Immature | Strong | Immature Areas |
| Jimenez-Alvarado et al., 2020 | Playa Flamingo, Lanzarote | *Squatina squatina* | ✓ | ✓ | ✗ | ✓ | ✗ | Strong | Moderate | Immature | Strong | Immature Areas |
| Jimenez-Alvarado et al., 2020 | Playa Honda, Lanzarote | *Squatina squatina* | ✗ | ✓ | ✗ | ✓ | ✗ | Strong | Moderate | Immature | Strong | Immature Areas |
| Jimenez-Alvarado et al., 2020 | Playa Mogan, Gran Canaria | *Squatina squatina* | ✓ | ✓ | ✗ | ✓ | ✗ | Strong | Moderate | Immature | Strong | Immature Areas |
| Jimenez-Alvarado et al., 2020 | Playa Poris de Abona, Tenerife | *Squatina squatina* | ✓ | ✓ | ✗ | ✗ | ✗ | Strong | Moderate | Immature | Strong | Immature Areas |
| Jimenez-Alvarado et al., 2020 | Playa de Melanara, Gran Canaria | *Squatina squatina* | ✓ | ✗ | ✗ | ✓ | ✗ | Strong | Moderate | Immature | Strong | Immature Areas |
| Jimenez-Alvarado et al., 2020 | Playa de Sardina del Norte, Gran Canaria | *Squatina squatina* | ✓ | ✗ | ✗ | ✓ | ✗ | Strong | Moderate | Immature | Strong | Immature Areas |
| Jimenez-Alvarado et al., 2020 | Playa de la concha, Fuerteventura | *Squatina squatina* | ✓ | ✗ | ✗ | ✓ | ✗ | Strong | Moderate | Immature | Strong | Immature Areas |
| Jimenez-Alvarado et al., 2020 | Playa de las Vistas, Tenerife | *Squatina squatina* | ✓ | ✓ | ✗ | ✓ | ✗ | Strong | Moderate | Immature | Strong | Immature Areas |
| Jimenez-Alvarado et al., 2020 | Playa del Cabron, Gran Canaria | *Squatina squatina* | ✓ | ✓ | ✗ | ✓ | ✗ | Strong | Moderate | Immature | Strong | Immature Areas |
| Jimenez-Alvarado et al., 2020 | Playa del Muelle, Gran Canaria | *Squatina squatina* | ✗ | ✓ | ✗ | ✓ | ✗ | Strong | Moderate | Immature | Strong | Immature Areas |
| Jimenez-Alvarado et al., 2020 | Playa del Muellito, Fuerteventura | *Squatina squatina* | ✗ | ✓ | ✗ | ✓ | ✗ | Strong | Moderate | Immature | Strong | Immature Areas |
| Jimenez-Alvarado et al., 2020 | Playa del Pozo, Fuerteventura | *Squatina squatina* | ✓ | ✓ | ✗ | ✗ | ✗ | Strong | Moderate | Immature | Strong | Immature Areas |
| Jimenez-Alvarado et al., 2020 | Playa del castillo, Fuerteventura | *Squatina squatina* | ✓ | ✓ | ✗ | ✓ | ✓ | Strong | Moderate | Immature | Strong | Immature Areas |
| Jimenez-Alvarado et al., 2020 | Playa del jablito, Fuerteventura | *Squatina squatina* | ✓ | ✓ | ✗ | ✓ | ✗ | Strong | Moderate | Immature | Strong | Immature Areas |
| Jimenez-Alvarado et al., 2020 | Playa del puertito, Fuerteventura | *Squatina squatina* | ✗ | ✓ | ✗ | ✓ | ✗ | Strong | Moderate | Immature | Strong | Immature Areas |
| Couto et al., 2017 | Southern Portugal | *Cetorhinus maximus* | ✗ | ✗ | ✗ | ✗ | ✗ | Weak | High | Neonate | Weak-Cannot Assess | Immature Areas |
| Hayes et al., 2018 | West of Shetland, Scotland | *Cetorhinus maximus* | ✗ | ✗ | ✗ | ✗ | ✗ | Weak | Moderate | Neonate | Weak-Cannot Assess | Immature Areas |
| Frost et al., 2020, Delaval et al., 2023, Brown-Vuillemin et al., 2020, Delaval et al., 2022 | Celtic Sea | *Dipturus batis* | ✗ | ✗ | ✗ | ✓ | ✗ | Weak | High | Juveniles | Weak-Cannot Assess | Immature Areas |
| Ellis et al., 2024 | Celtic Sea | *Dipturus batis* | ✗ | ✗ | ✗ | ✗ | ✗ | Weak | High | 1st Cohort | Weak-Cannot Assess | Immature Areas |
| Frost et al., 2020 | Far North, Scotland | *Dipturus batis* | ✓ | ✗ | ✗ | ✗ | ✗ | Weak | High | Young-of-the-year | Weak-Cannot Assess | Immature Areas |

*This column highlights (in grey) when a cap was placed on the overall score based on the rules outlined in the risk of bias assessment.

| Reference(s) | Location | Species | R1 | R2 (Species) | R2 (Individual) | R3 (Species) | R3 (Individual) | Strength of Evidence | Quality Assessment | Evidence Cap* | Overall Evidence | Critical Area |
|---|---|---|---|---|---|---|---|---|---|---|---|---|
| Delaval et al., 2022 | Faroe Bank | *Dipturus batis* | ✗ | ✗ | ✗ | ✗ | ✗ | Weak | Low | Juveniles | Weak-Cannot Assess | Immature Areas |
| Delaval et al., 2022 | Faroe Shelf | *Dipturus batis* | ✗ | ✗ | ✗ | ✗ | ✗ | Weak | Low | Juveniles | Weak-Cannot Assess | Immature Areas |
| Delaval et al., 2022 | North Scotland | *Dipturus batis* | ✗ | ✗ | ✗ | ✗ | ✗ | Weak | Moderate | Juveniles | Weak-Cannot Assess | Immature Areas |
| Delaval et al., 2022; Frost et al., 2020 | Rockall | *Dipturus batis* | ✓ | ✗ | ✗ | ✗ | ✗ | Weak | High | Neonate | Weak-Cannot Assess | Immature Areas |
| Delaval et al., 2022; Frost et al., 2020 | West Coast Scotland | *Dipturus batis* | ✓ | ✗ | ✗ | ✗ | ✗ | Weak | High | Neonate | Weak-Cannot Assess | Immature Areas |
| Ellis et al., 2024 | Western Ireland | *Dipturus batis* | ✗ | ✗ | ✗ | ✗ | ✗ | Weak | High | Neonate | Weak-Cannot Assess | Immature Areas |
| Ellis et al., 2024 | Western to Northern Scotland | *Dipturus batis* | ✗ | ✗ | ✗ | ✗ | ✗ | Weak | High | Neonate | Weak-Cannot Assess | Immature Areas |
| Ellis et al., 2024 | Celtic Sea | *Dipturus intermedius* | ✗ | ✗ | ✗ | ✗ | ✗ | Weak | High | Neonate | Weak-Cannot Assess | Immature Areas |
| Frost et al., 2020 | Far North, Scotland | *Dipturus intermedius* | ✓ | ✗ | ✗ | ✗ | ✗ | Weak | High | Neonate | Weak-Cannot Assess | Immature Areas |
| Benjamins et al., 2018 | Loch Sunart Sound of Jura MPA | *Dipturus intermedius* | ✗ | ✗ | ✗ | ✗ | ✗ | Weak | High | Juveniles | Weak-Cannot Assess | Immature Areas |
| Frost et al., 2020 | North Shelf, Scotland | *Dipturus intermedius* | ✓ | ✗ | ✗ | ✗ | ✗ | Weak | High | Neonate | Weak-Cannot Assess | Immature Areas |
| Frost et al., 2020 | South Shelf, Scotland | *Dipturus intermedius* | ✓ | ✗ | ✗ | ✗ | ✗ | Weak | High | Neonate | Weak-Cannot Assess | Immature Areas |
| Frost et al., 2020 | West coast, Scotland | *Dipturus intermedius* | ✗ | ✗ | ✗ | ✗ | ✗ | Weak | High | Neonate | Weak-Cannot Assess | Immature Areas |
| Ellis et al., 2024 | Western Ireland | *Dipturus intermedius* | ✗ | ✗ | ✗ | ✗ | ✗ | Weak | High | Neonate | Weak-Cannot Assess | Immature Areas |
| Ellis et al., 2024 | Western to Northern Scotland and Shetland | *Dipturus intermedius* | ✗ | ✗ | ✗ | ✗ | ✗ | Weak | High | Neonate | Weak-Cannot Assess | Immature Areas |
| Neat et al., 2015a | Rockall Trough, Continental slope (950m) | *Dipturus nidarosiensis* | ✓ | ✗ | ✗ | ✗ | ✗ | Weak | High | Juveniles | Weak-Cannot Assess | Immature Areas |
| Neat et al., 2015a | Southern Rockall Trough | *Galeus melastomus* | ✓ | ✗ | ✗ | ✗ | ✗ | Weak | High | Immature | Weak-Cannot Assess | Immature Areas |
| Carrasco-Puig et al., 2024 | Bay of biscay (NW Spain) Asturias, Spain | *Lamna nasus* | ✗ | ✓ | ✗ | ✗ | ✗ | Weak | Moderate | Young-of-the-year | Weak-Cannot Assess | Immature Areas |
| Serra-Pereira et al., 2014 | Mar do Canhimbo, Portugal | *Leucoraja naevus* | ✗ | ✗ | ✗ | ✗ | ✗ | Weak | High | Potentially Mature | Weak-Cannot Assess | Immature Areas |
| Ellis et al., 2024 | Northern Celtic Sea & Southwest of Cornwall | *Leucoraja naevus* | ✗ | ✗ | ✗ | ✗ | ✗ | Weak | High | 1st Cohort | Weak-Cannot Assess | Immature Areas |
| Ellis et al., 2024 | Scotland | *Leucoraja naevus* | ✓ | ✗ | ✗ | ✗ | ✗ | Weak | High | 1st Cohort | Weak-Cannot Assess | Immature Areas |

*This column highlights (in grey) when a cap was placed on the overall score based on the rules outlined in the risk of bias assessment.

| Reference(s) | Location | Species | R1 | R2 (Species) | R2 (Individual) | R3 (Species) | R3 (Individual) | Strength of Evidence | Quality Assessment | Evidence Cap* | Overall Evidence | Critical Area |
|---|---|---|---|---|---|---|---|---|---|---|---|---|
| Serra-Pereira et al., 2014 | Off Areia Branca, Portugal | *Leucoraja naevus* | ✗ | ✗ | ✗ | ✗ | ✗ | Weak | Moderate | Potentially Mature | Weak-Cannot Assess | Immature Areas |
| Ellis et al., 2024 | Celtic Sea | *Leucoraja fullonica* | ✗ | ✗ | ✗ | ✗ | ✗ | Weak | High | Juveniles | Weak-Cannot Assess | Immature Areas |
| Ellis et al., 2024 | Northwestern North Sea | *Leucoraja fullonica* | ✗ | ✗ | ✗ | ✗ | ✗ | Weak | High | Juveniles | Weak-Cannot Assess | Immature Areas |
| Ellis et al., 2024 | Southwestern Ireland | *Leucoraja fullonica* | ✗ | ✗ | ✗ | ✗ | ✗ | Weak | High | Juveniles | Weak-Cannot Assess | Immature Areas |
| Ellis et al., 2024 | West of Scotland (Deep waters) | *Leucoraja fullonica* | ✗ | ✗ | ✗ | ✗ | ✗ | Weak | High | Juveniles | Weak-Cannot Assess | Immature Areas |
| Serra-Pereira et al., 2014 | Berlenga, Portugal | *Leucoraja naevus* | ✗ | ✗ | ✗ | ✗ | ✗ | Weak | High | Potentially Mature | Weak-Cannot Assess | Immature Areas |
| Serra-Pereira et al., 2014 | Foz do Arelho, Portugal | *Leucoraja naevus* | ✗ | ✗ | ✗ | ✗ | ✗ | Weak | Moderate | Potentially Mature | Weak-Cannot Assess | Immature Areas |
| Serra-Pereira et al., 2014 | Off Areia Branca, Portugal | *Leucoraja naevus* | ✗ | ✗ | ✗ | ✗ | ✗ | Weak | High | Potentially Mature | Weak-Cannot Assess | Immature Areas |
| Serra-Pereira et al., 2014 | Off Santa Cruz, Portugal | *Leucoraja naevus* | ✗ | ✗ | ✗ | ✗ | ✗ | Weak | High | Potentially Mature | Weak-Cannot Assess | Immature Areas |
| Breve et al., 2020; McCully-Phillips & Ellis, 2015 | Southern North Sea & Eastern English Channel | *Mustelus asterias* | ✗ | ✓ | ✗ | ✗ | ✗ | Weak | High | Neonates | Weak-Cannot Assess | Immature Areas |
| McCully-Phillips & Ellis, 2015 | Irish Sea | *Mustelus asterias* | ✗ | ✗ | ✗ | ✗ | ✗ | Weak | High | Neonates | Weak-Cannot Assess | Immature Areas |
| McCully-Phillips & Ellis, 2015 | Western English Channel | *Mustelus asterias* | ✗ | ✗ | ✗ | ✗ | ✗ | Weak | High | Neonates | Weak-Cannot Assess | Immature Areas |
| Breve et al., 2016 | surge barrier Neeltje Jans, just outside the Oosterschelde, Dutch delta, Netherlands | *Mustelus asterias* | ✗ | ✗ | ✗ | ✗ | ✗ | Weak | High | Neonates | Weak-Cannot Assess | Immature Areas |
| Serra-Pereira et al., 2014 | Baleal, Portugal | *Raja brachyura* | ✗ | ✗ | ✗ | ✗ | ✗ | Weak | High | Immature | Weak-Cannot Assess | Immature Areas |
| Ellis et al., 2024 | Channel Islands | *Raja brachyura* | ✗ | ✗ | ✗ | ✗ | ✗ | Weak | High | 1st Cohort | Weak-Cannot Assess | Immature Areas |
| Serra-Pereira et al., 2014 | Foz do Arelho, Portugal | *Raja brachyura* | ✗ | ✗ | ✗ | ✗ | ✗ | Weak | High | Immature | Weak-Cannot Assess | Immature Areas |
| Ellis et al., 2024 | Inshore Western English Channel | *Raja brachyura* | ✗ | ✗ | ✗ | ✗ | ✗ | Weak | High | 1st Cohort | Weak-Cannot Assess | Immature Areas |
| Ellis et al., 2024 | Liverpool Bay, Irish Sea | *Raja brachyura* | ✗ | ✗ | ✗ | ✗ | ✗ | Weak | High | 1st Cohort | Weak-Cannot Assess | Immature Areas |
| Serra-Pereira et al., 2014 | Mar do Canhimbo, Portugal | *Raja brachyura* | ✗ | ✗ | ✗ | ✗ | ✗ | Weak | High | Immature | Weak-Cannot Assess | Immature Areas |
| Serra-Pereira et al., 2014 | Off Areia Branca, Portugal | *Raja brachyura* | ✗ | ✗ | ✗ | ✗ | ✗ | Weak | High | Immature | Weak-Cannot Assess | Immature Areas |
| Serra-Pereira et al., 2014 | Off Santa Cruz, Portugal | *Raja brachyura* | ✗ | ✗ | ✗ | ✗ | ✗ | Weak | High | Immature | Weak-Cannot Assess | Immature Areas |

*This column highlights (in grey) when a cap was placed on the overall score based on the rules outlined in the risk of bias assessment.

| Reference(s) | Location | Species | R1 | R2 (Species) | R2 (Individual) | R3 (Species) | R3 (Individual) | Strength of Evidence | Quality Assessment | Evidence Cap* | Overall Evidence | Critical Area |
|---|---|---|---|---|---|---|---|---|---|---|---|---|
| Ellis et al., 2024 | Orkney | *Raja brachyura* | ✗ | ✗ | ✗ | ✗ | ✗ | Weak | High | 1st Cohort | Weak-Cannot Assess | Immature Areas |
| Serra-Pereira et al., 2014 | Santa Cruz, Portugal | *Raja brachyura* | ✗ | ✗ | ✗ | ✗ | ✗ | Weak | High | Immature | Weak-Cannot Assess | Immature Areas |
| Ellis et al., 2024 | Southwestern Scotland | *Raja brachyura* | ✗ | ✗ | ✗ | ✗ | ✗ | Weak | High | 1st Cohort | Weak-Cannot Assess | Immature Areas |
| Ellis et al., 2024 | Western Ireland | *Raja brachyura* | ✗ | ✗ | ✗ | ✗ | ✗ | Weak | High | 1st Cohort | Weak-Cannot Assess | Immature Areas |
| Serra-Pereira et al., 2014 | Areia Branca, Portugal | *Raja clavata* | ✗ | ✗ | ✗ | ✗ | ✗ | Weak | High | Immature | Weak-Cannot Assess | Immature Areas |
| Santos et al., 2021 | Around the Azores (in habitats near Islands) | *Raja clavata* | ✓ | ✗ | ✗ | ✗ | ✗ | Weak | High | Juveniles | Weak-Cannot Assess | Immature Areas |
| Serra-Pereira et al., 2014 | Baleal, Portugal | *Raja clavata* | ✗ | ✗ | ✗ | ✗ | ✗ | Weak | High | Immature | Weak-Cannot Assess | Immature Areas |
| Serra-Pereira et al., 2014 | Berlenga, Portugal | *Raja clavata* | ✗ | ✗ | ✗ | ✗ | ✗ | Weak | High | Immature | Weak-Cannot Assess | Immature Areas |
| Serra-Pereira et al., 2014 | Foz do Arelho, Portugal | *Raja clavata* | ✗ | ✗ | ✗ | ✗ | ✗ | Weak | High | Immature | Weak-Cannot Assess | Immature Areas |
| Biton-Porsmoguer et al., 2020 | Littoral Seino-Marin, France | *Raja clavata* | ✗ | ✗ | ✗ | ✗ | ✗ | Weak | Moderate | Potentially Mature | Weak-Cannot Assess | Immature Areas |
| Ellis et al., 2024 | Lyme Bay, Western English Channel | *Raja clavata* | ✗ | ✗ | ✗ | ✗ | ✗ | Weak | High | 1st Cohort | Weak-Cannot Assess | Immature Areas |
| Serra-Pereira et al., 2014 | Ericeira, Portugal | *Raja clavata* | ✗ | ✗ | ✗ | ✗ | ✗ | Weak | High | Immature | Weak-Cannot Assess | Immature Areas |
| Serra-Pereira et al., 2014 | Mar do Canhimbo, Portugal | *Raja clavata* | ✗ | ✗ | ✗ | ✗ | ✗ | Weak | High | Immature | Weak-Cannot Assess | Immature Areas |
| Serra-Pereira et al., 2014 | NW Cabo da Roca, Portugal | *Raja clavata* | ✗ | ✗ | ✗ | ✗ | ✗ | Weak | High | Immature | Weak-Cannot Assess | Immature Areas |
| Serra-Pereira et al., 2014 | Off Areia Branca, Portugal | *Raja clavata* | ✗ | ✗ | ✗ | ✗ | ✗ | Weak | High | Immature | Weak-Cannot Assess | Immature Areas |
| Ellis et al., 2024 | Off Arklow, Irish Sea | *Raja clavata* | ✓ | ✗ | ✗ | ✗ | ✗ | Weak | High | 1st Cohort | Weak-Cannot Assess | Immature Areas |
| Serra-Pereira et al., 2014 | Off Santa Cruz, Portugal | *Raja clavata* | ✗ | ✗ | ✗ | ✗ | ✗ | Weak | High | Immature | Weak-Cannot Assess | Immature Areas |
| Serra-Pereira et al., 2014 | Santa Cruz, Portugal | *Raja clavata* | ✗ | ✗ | ✗ | ✗ | ✗ | Weak | High | Immature | Weak-Cannot Assess | Immature Areas |
| Ellis et al., 2024 | Southwestern Ireland | *Raja clavata* | ✗ | ✗ | ✗ | ✗ | ✗ | Weak | High | 1st Cohort | Weak-Cannot Assess | Immature Areas |
| Ellis et al., 2024 | Western Ireland | *Raja clavata* | ✗ | ✗ | ✗ | ✗ | ✗ | Weak | High | Juveniles | Weak-Cannot Assess | Immature Areas |
| Ellis et al., 2024 | Western Scotland | *Raja clavata* | ✗ | ✗ | ✗ | ✗ | ✗ | Weak | High | Juveniles | Weak-Cannot Assess | Immature Areas |

*This column highlights (in grey) when a cap was placed on the overall score based on the rules outlined in the risk of bias assessment.

| Reference(s) | Location | Species | R1 | R2 (Species) | R2 (Individual) | R3 (Species) | R3 (Individual) | Strength of Evidence | Quality Assessment | Evidence Cap* | Overall Evidence | Critical Area |
|---|---|---|---|---|---|---|---|---|---|---|---|---|
| Serra-Pereira et al., 2014 | Areia Branca, Portugal | *Raja microocellata* | ✗ | ✗ | ✗ | ✗ | ✗ | Weak | High | Immature | Weak-Cannot Assess | Immature Areas |
| Ellis et al., 2024 | Eastern English Channel | *Raja microocellata* | ✓ | ✗ | ✗ | ✗ | ✗ | Weak | High | Juveniles | Weak-Cannot Assess | Immature Areas |
| Serra-Pereira et al., 2014 | Foz do Arelho, Portugal | *Raja microocellata* | ✗ | ✗ | ✗ | ✗ | ✗ | Weak | High | Immature | Weak-Cannot Assess | Immature Areas |
| Serra-Pereira et al., 2014 | Off Areia Branca, Portugal | *Raja microocellata* | ✗ | ✗ | ✗ | ✗ | ✗ | Weak | Moderate | Immature | Weak-Cannot Assess | Immature Areas |
| Serra-Pereira et al., 2014 | Off Santa Cruz, Portugal | *Raja microocellata* | ✗ | ✗ | ✗ | ✗ | ✗ | Weak | High | Immature | Weak-Cannot Assess | Immature Areas |
| Serra-Pereira et al., 2014 | Santa Cruz, Portugal | *Raja microocellata* | ✗ | ✗ | ✗ | ✗ | ✗ | Weak | High | Immature | Weak-Cannot Assess | Immature Areas |
| Ellis et al., 2024 | Southern England | *Raja microocellata* | ✓ | ✗ | ✗ | ✗ | ✗ | Weak | High | Juveniles | Weak-Cannot Assess | Immature Areas |
| Serra-Pereira et al., 2014 | Baleal, Portugal | *Raja montagui* | ✗ | ✗ | ✗ | ✗ | ✗ | Weak | High | Potentially Mature | Weak-Cannot Assess | Immature Areas |
| Serra-Pereira et al., 2014 | Berlenga, Portugal | *Raja montagui* | ✗ | ✗ | ✗ | ✗ | ✗ | Weak | High | Potentially Mature | Weak-Cannot Assess | Immature Areas |
| Ellis et al., 2024 | Eas Anglia Coast | *Raja montagui* | ✗ | ✗ | ✗ | ✗ | ✗ | Weak | High | 1st Cohort | Weak-Cannot Assess | Immature Areas |
| Ellis et al., 2024 | Eastern English Channel/ Southern North Sea | *Raja montagui* | ✓ | ✗ | ✗ | ✗ | ✗ | Weak | High | 1st Cohort | Weak-Cannot Assess | Immature Areas |
| Serra-Pereira et al., 2014 | Foz do Arelho, Portugal | *Raja montagui* | ✗ | ✗ | ✗ | ✗ | ✗ | Weak | High | Potentially Mature | Weak-Cannot Assess | Immature Areas |
| Ellis et al., 2024 | Isles of Scilly to Lyme Bay, Southern England | *Raja montagui* | ✗ | ✗ | ✗ | ✗ | ✗ | Weak | High | 1st Cohort | Weak-Cannot Assess | Immature Areas |
| Biton-Porsmoguer et al., 2020 | Littoral Seino-Marin, France | *Raja montagui* | ✗ | ✗ | ✗ | ✗ | ✗ | Weak | Moderate | Juveniles | Weak-Cannot Assess | Immature Areas |
| Serra-Pereira et al., 2014 | Ericeira, Portugal | *Raja montagui* | ✗ | ✗ | ✗ | ✗ | ✗ | Weak | High | Potentially Mature | Weak-Cannot Assess | Immature Areas |
| Serra-Pereira et al., 2014 | Mar do Cachimbo, Portugal | *Raja montagui* | ✗ | ✗ | ✗ | ✗ | ✗ | Weak | High | Potentially Mature | Weak-Cannot Assess | Immature Areas |
| Ellis et al., 2024 | Northern Celtic Sea | *Raja montagui* | ✗ | ✗ | ✗ | ✗ | ✗ | Weak | High | Juveniles | Weak-Cannot Assess | Immature Areas |
| Serra-Pereira et al., 2014 | Off Areia Branca, Portugal | *Raja montagui* | ✗ | ✗ | ✗ | ✗ | ✗ | Weak | High | Potentially Mature | Weak-Cannot Assess | Immature Areas |
| Serra-Pereira et al., 2014 | Off Santa Cruz, Portugal | *Raja montagui* | ✗ | ✗ | ✗ | ✗ | ✗ | Weak | High | Potentially Mature | Weak-Cannot Assess | Immature Areas |
| Serra-Pereira et al., 2014 | Santa Cruz, Portugal | *Raja montagui* | ✗ | ✗ | ✗ | ✗ | ✗ | Weak | High | Potentially Mature | Weak-Cannot Assess | Immature Areas |
| Ellis et al., 2024 | West of Ireland | *Raja montagui* | ✗ | ✗ | ✗ | ✗ | ✗ | Weak | High | Juveniles | Weak-Cannot Assess | Immature Areas |

*This column highlights (in grey) when a cap was placed on the overall score based on the rules outlined in the risk of bias assessment.

| Reference(s) | Location | Species | R1 | R2 (Species) | R2 (Individual) | R3 (Species) | R3 (Individual) | Strength of Evidence | Quality Assessment | Evidence Cap* | Overall Evidence | Critical Area |
|---|---|---|---|---|---|---|---|---|---|---|---|---|
| Ellis et al., 2024 | West of Scotland | *Raja montagui* | ✗ | ✗ | ✗ | ✗ | ✗ | Weak | High | Juveniles | Weak-Cannot Assess | Immature Areas |
| Serra-Pereira et al., 2014 | Areia Branca, Portugal | *Raja undulata* | ✗ | ✗ | ✗ | ✗ | ✗ | Weak | High | Immature | Weak-Cannot Assess | Immature Areas |
| Serra-Pereira et al., 2014 | Baleal, Portugal | *Raja undulata* | ✗ | ✗ | ✗ | ✗ | ✗ | Weak | High | Immature | Weak-Cannot Assess | Immature Areas |
| Ellis et al., 2024 | Eastern English Channel | *Raja undulata* | ✓ | ✗ | ✗ | ✗ | ✗ | Weak | High | 1st Cohort | Weak-Cannot Assess | Immature Areas |
| Serra-Pereira et al., 2015 | Estuaries in Portugal | *Raja undulata* | ✗ | ✗ | ✗ | ✗ | ✗ | Weak | Low | Immature | Weak-Cannot Assess | Immature Areas |
| Serra-Pereira et al., 2014 | Foz do Arelho, Portugal | *Raja undulata* | ✗ | ✗ | ✗ | ✗ | ✗ | Weak | Moderate | Immature | Weak-Cannot Assess | Immature Areas |
| Serra-Pereira et al., 2014 | Mar do Cachimbo, Portugal | *Raja undulata* | ✗ | ✗ | ✗ | ✗ | ✗ | Weak | High | Immature | Weak-Cannot Assess | Immature Areas |
| Serra-Pereira et al., 2014 | Off Areia Branca, Portugal | *Raja undulata* | ✗ | ✗ | ✗ | ✗ | ✗ | Weak | High | Immature | Weak-Cannot Assess | Immature Areas |
| Serra-Pereira et al., 2014 | Off Santa Cruz, Portugal | *Raja undulata* | ✗ | ✗ | ✗ | ✗ | ✗ | Weak | High | Immature | Weak-Cannot Assess | Immature Areas |
| Serra-Pereira et al., 2014 | Santa Cruz, Portugal | *Raja undulata* | ✗ | ✗ | ✗ | ✗ | ✗ | Weak | High | Immature | Weak-Cannot Assess | Immature Areas |
| Ellis et al., 2024 | Southern England | *Raja undulata* | ✗ | ✗ | ✗ | ✗ | ✗ | Weak | High | 1st Cohort | Weak-Cannot Assess | Immature Areas |
| Ellis et al., 2024 | Western Ireland | *Rostroraja alba* | ✗ | ✗ | ✗ | ✗ | ✗ | Weak | High | Juveniles | Weak-Cannot Assess | Immature Areas |
| Jimenez-Alvarado et al., 2020 | El Balito, Tenerife | *Squatina squatina* | ✗ | ✗ | ✗ | ✗ | ✗ | Weak | Low | Immature | Weak-Cannot Assess | Immature Areas |
| Barker et al., 2022 | Irish Sea | *Squatina squatina* | ✗ | ✗ | ✗ | ✗ | ✗ | Weak | High | Juveniles | Weak-Cannot Assess | Immature Areas |
| Jimenez-Alvarado et al., 2020 | Playa Corralejo, Fuerteventura | *Squatina squatina* | ✓ | ✗ | ✗ | ✗ | ✗ | Weak | Moderate | Immature | Weak-Cannot Assess | Immature Areas |
| Jimenez-Alvarado et al., 2020 | Playa Ginginamar, Fuerteventura | *Squatina squatina* | ✗ | ✗ | ✗ | ✗ | ✗ | Weak | Moderate | Immature | Weak-Cannot Assess | Immature Areas |
| Jimenez-Alvarado et al., 2020 | Playa de Amadores, Gran Canaria | *Squatina squatina* | ✗ | ✗ | ✗ | ✗ | ✗ | Weak | Low | Immature | Weak-Cannot Assess | Immature Areas |
| Jimenez-Alvarado et al., 2020 | Playa de Anfi Del Mar, Gran Canaria | *Squatina squatina* | ✗ | ✗ | ✗ | ✗ | ✗ | Weak | Moderate | Immature | Weak-Cannot Assess | Immature Areas |
| Jimenez-Alvarado et al., 2020 | Playa de Atlantico, Lanzarote | *Squatina squatina* | ✗ | ✗ | ✗ | ✗ | ✗ | Weak | Low | Immature | Weak-Cannot Assess | Immature Areas |
| Jimenez-Alvarado et al., 2020 | Playa de Fanabe, Tenerife | *Squatina squatina* | ✗ | ✗ | ✗ | ✗ | ✗ | Weak | Low | Immature | Weak-Cannot Assess | Immature Areas |
| Jimenez-Alvarado et al., 2020 | Playa de Fariones, Lanzarote | *Squatina squatina* | ✓ | ✗ | ✗ | ✗ | ✗ | Weak | Moderate | Immature | Weak-Cannot Assess | Immature Areas |

*This column highlights (in grey) when a cap was placed on the overall score based on the rules outlined in the risk of bias assessment.

| Reference(s) | Location | Species | R1 | R2 (Species) | R2 (Individual) | R3 (Species) | R3 (Individual) | Strength of Evidence | Quality Assessment | Evidence Cap* | Overall Evidence | Critical Area |
|---|---|---|---|---|---|---|---|---|---|---|---|---|
| Jimenez-Alvarado et al., 2020 | Playa de Puerto Rico, Gran Canaria | *Squatina squatina* | ✓ | ✗ | ✗ | ✗ | ✗ | Weak | Low | Immature | Weak-Cannot Assess | Immature Areas |
| Jimenez-Alvarado et al., 2020 | Playa de San Juan, Tenerife | *Squatina squatina* | ✓ | ✗ | ✗ | ✗ | ✗ | Weak | Low | Immature | Weak-Cannot Assess | Immature Areas |
| Jimenez-Alvarado et al., 2020 | Playa de San Sebastian, La Graciosa | *Squatina squatina* | ✗ | ✗ | ✗ | ✗ | ✗ | Weak | Low | Immature | Weak-Cannot Assess | Immature Areas |
| Tuya et al., 2020, Jimenez-Alvarado et al., 2020 | Playa de la Canteras, Gran Canaria | *Squatina squatina* | ✗ | ✓ | ✗ | ✗ | ✗ | Weak | High | Juveniles | Weak-Cannot Assess | Immature Areas |
| Jimenez-Alvarado et al., 2020 | Playa de la Lajita, Fuerteventura | *Squatina squatina* | ✓ | ✗ | ✗ | ✗ | ✗ | Weak | Moderate | Immature | Weak-Cannot Assess | Immature Areas |
| Jimenez-Alvarado et al., 2020 | Playa de las Playitas, Fuerteventura | *Squatina squatina* | ✗ | ✗ | ✗ | ✗ | ✗ | Weak | Low | Immature | Weak-Cannot Assess | Immature Areas |
| Jimenez-Alvarado et al., 2020 | Playa de los Cristianos, Tenerife | *Squatina squatina* | ✗ | ✓ | ✗ | ✗ | ✗ | Weak | Moderate | Immature | Weak-Cannot Assess | Immature Areas |
| Jimenez-Alvarado et al., 2020 | Playa del Jabillo, Lanzarote | *Squatina squatina* | ✗ | ✗ | ✗ | ✗ | ✗ | Weak | Low | Immature | Weak-Cannot Assess | Immature Areas |
| Jimenez-Alvarado et al., 2020 | Playa del Reducto, Lanzarote | *Squatina squatina* | ✓ | ✗ | ✗ | ✗ | ✗ | Weak | Moderate | Immature | Weak-Cannot Assess | Immature Areas |
| Jimenez-Alvarado et al., 2020 | Puerto de Granadilla, Tenerife | *Squatina squatina* | ✗ | ✗ | ✗ | ✗ | ✗ | Weak | Moderate | Immature | Weak-Cannot Assess | Immature Areas |
| Jimenez-Alvarado et al., 2020 | Puerto de Santa Cruz, Tenerife | *Squatina squatina* | ✗ | ✗ | ✗ | ✗ | ✗ | Weak | Low | Immature | Weak-Cannot Assess | Immature Areas |
| Meyers et al., 2017 | Waters around Gran Canaria, Canary Islands | *Squatina squatina* | ✓ | ✗ | ✗ | ✗ | ✗ | Weak | High | Neonates | Weak-Cannot Assess | Immature Areas |
| Meyers et al., 2017 | Waters around Lanzarote, Canary Islands | *Squatina squatina* | ✓ | ✗ | ✗ | ✗ | ✗ | Weak | High | Neonates | Weak-Cannot Assess | Immature Areas |
| Meyers et al., 2017 | Waters around Tenerife, Canary Islands | *Squatina squatina* | ✓ | ✗ | ✗ | ✗ | ✗ | Weak | High | Neonates | Weak-Cannot Assess | Immature Areas |
| Barker et al., 2022 | Welsh Coastal Waters | *Squatina squatina* | ✗ | ✗ | ✗ | ✗ | ✗ | Weak | High | Immature | Weak-Cannot Assess | Immature Areas |
| Thorburn et al., 2024 | Achill Island, Ireland | *Cetorhinus maximus* | ✗ | ✗ | ✗ | ✗ | ✗ | Weak | High | Juveniles | Weak-Cannot Assess | Immature Areas |
| **Mating/Courtship** | | | | | | | | | | **Type of Mating Evidence** | | |
| Sims et al., 2022 | Coastal waters off County Clare & County Galway, Ireland | *Cetorhinus maximus* | ✗ | ✓ | ✓ | ✓ | ✗ | Strong | High | Direct Observation of Courtship | Strong | Mating/Courtship |
| Simpson et al., 2021 | Coastal waters off Mothecombe & Bigbury-on-sea (Southwest UK) | *Raja brachyura* | ✓ | ✓ | ✗ | ✗ | ✗ | Strong | High | Male-Female Overlap | Moderate | Mating/Courtship |
| Lavender et al., 2022 | Loch Sunart & Sound of Jura MPA, Scotland | *Dipturus intermedius* | ✗ | ✓ | ✓ | ✓ | ✓ | Strong | High | Mature Adult Overlap | Moderate | Mating/Courtship |
| Sims et al., 2022, Rudd et al., 2021 | Coll & Tiree waters, Scotland | *Cetorhinus maximus* | ✗ | ✗ | ✗ | ✓ | ✗ | Weak | Moderate | Direct Observation of Courtship | Weak-Cannot Assess | Mating/Courtship |

*This column highlights (in grey) when a cap was placed on the overall score based on the rules outlined in the risk of bias assessment.

| Reference(s) | Location | Species | R1 | R2 (Species) | R2 (Individual) | R3 (Species) | R3 (Individual) | Strength of Evidence | Quality Assessment | Evidence Cap* | Overall Evidence | Critical Area |
|---|---|---|---|---|---|---|---|---|---|---|---|---|
| Simpson et al., 2021 | Coastal waters off Mothecombe & Bigbury-on-sea (Southwest UK) | *Raja montagui* | ✓ | ✗ | ✗ | ✗ | ✗ | Weak | High | Male-Female Overlap | Weak-Cannot Assess | Mating/Courtship |
| Simpson et al., 2021 | Whitsand Bay, Southwest UK | *Raja microocellata* | ✓ | ✗ | ✗ | ✗ | ✗ | Weak | High | Male-Female Overlap | Weak-Cannot Assess | Mating/Courtship |
| Simpson et al., 2021 | Whitsand Bay, Southwest UK | *Raja clavata* | ✓ | ✗ | ✗ | ✗ | ✗ | Weak | High | Male-Female Overlap | Weak-Cannot Assess | Mating/Courtship |
| Serra-Pereira et al., 2015 | Portguese coastal waters | *Raja undulata* | ✗ | ✗ | ✗ | ✗ | ✗ | Weak | High | Reproductively Active Males | Weak-Cannot Assess | Mating/Courtship |
| Rudd et al., 2021 | Coll & Tiree waters, Scotland | *Cetorhinus maximus* | ✗ | ✗ | ✗ | ✓ | ✗ | Weak | High | Direct Observation of Courtship | Weak-Cannot Assess | Mating/Courtship |
| Meyers et al., 2017 | Waters around the Canary Islands | *Squatina squatina* | ✗ | ✗ | ✗ | ✗ | ✗ | Weak | High | Mating Scars/ Male-Female Overlap | Weak-Cannot Assess | Mating/Courtship |
| McCully-Phillips & Ellis, 2015 | English Channel | *Mustelus asterias* | ✗ | ✗ | ✗ | ✗ | ✗ | Weak | High | Reproductively Active Males | Weak-Cannot Assess | Mating/Courtship |
| Maia et al., 2012 | Inshore Portugese Waters | *Leucoraja naevus* | ✗ | ✗ | ✗ | ✗ | ✗ | Weak | High | Reproductively Active Adults | Weak-Cannot Assess | Mating/Courtship |
| Hayes et al., 2018 | West of Shetland, Scotland | *Cetorhinus maximus* | ✓ | ✗ | ✗ | ✗ | ✗ | Weak | High | Breaching | Weak-Cannot Assess | Mating/Courtship |
| Gore et al., 2019 | Inner hebrides, Scotland | *Cetorhinus maximus* | ✗ | ✗ | ✗ | ✗ | ✗ | Weak | High | Mating Scars | Weak-Cannot Assess | Mating/Courtship |
| Doherty et al., 2017a | Sea of the Hebrides MPA, Scotland | *Cetorhinus maximus* | ✗ | ✗ | ✗ | ✗ | ✗ | Weak | Low | Direct Observation of Courtship | Weak-Cannot Assess | Mating/Courtship |
| Breve et al., 2016 | Between Eastern English Channel and Bay of Biscay | *Mustelus asterias* | ✗ | ✗ | ✗ | ✗ | ✗ | Weak | Low | Unsure | Weak-Cannot Assess | Mating/Courtship |
| Santos et al., 2021 | Azores | *Raja clavata* | ✗ | ✗ | ✗ | ✗ | ✗ | Weak | High | Reproductively Active Adults | Weak-Cannot Assess | Mating/Courtship |
| **Resting** | | | | | | | | | | **Type of Resting Evidence** | | |
| Leeb et al., 2021 | Ria de Vigo, Spain | *Raja undulata* | ✗ | ✓ | ✓ | ✗ | ✗ | Moderate | Moderate | Activity Levels | Moderate | Resting |
| Sousa et al., 2019 | Professor Luiz Saldanha Marine Park, Portugal | *Rostroraja alba* | ✗ | ✗ | ✗ | ✗ | ✗ | Weak | Low | Activity Levels | Weak-Cannot Assess | Resting |
| Tuya et al., 2020 | Offshore, deeper areas adjacent to Las Canteras Beach | *Squatina squatina* | ✗ | ✗ | ✗ | ✗ | ✗ | Weak | High | Lower Occurrence | Weak-Cannot Assess | Resting |
| Simpson et al., 2021 | Plymouth coastal waters | *Raja brachyura* | ✗ | ✗ | ✗ | ✗ | ✗ | Weak | High | Habitat Preference | Weak-Cannot Assess | Resting |
| Simpson et al., 2021 | Plymouth coastal waters | *Raja clavata* | ✗ | ✗ | ✗ | ✗ | ✗ | Weak | High | Habitat Preference | Weak-Cannot Assess | Resting |
| Simpson et al., 2021 | Plymouth coastal waters | *Raja montagui* | ✗ | ✗ | ✗ | ✗ | ✗ | Weak | High | Habitat Preference | Weak-Cannot Assess | Resting |
| Simpson et al., 2021 | Plymouth coastal waters | *Raja microocellata* | ✗ | ✗ | ✗ | ✗ | ✗ | Weak | High | Habitat Preference | Weak-Cannot Assess | Resting |

*This column highlights (in grey) when a cap was placed on the overall score based on the rules outlined in the risk of bias assessment.

| Reference(s) | Location | Species | R1 | R2 (Species) | R2 (Individual) | R3 (Species) | R3 (Individual) | Strength of Evidence | Quality Assessment | Evidence Cap* | Overall Evidence | Critical Area |
|---|---|---|---|---|---|---|---|---|---|---|---|---|
| Mead et al., 2023 | La Graciosa Marine Reserve, Canary Islands | *Squatina squatina* | ✗ | ✗ | ✗ | ✗ | ✗ | Weak | Moderate | Activity Levels | Weak-Cannot Assess | Resting |
| Carrasco-Puig et al., 2024 | Bay of biscay (NW Spain) Asturias, Spain | *Lamna nasus* | ✗ | ✗ | ✗ | ✗ | ✗ | Weak | Low | No Clear Evidence | Weak-Cannot Assess | Resting |
| **Viviparous Gestation** | | | | | | | | | | **Type of Gestation Evidence** | | |
| Jung et al., 2024 | Tregor Area, France | *Lamna nasus* | ✗ | ✓ | ✓ | ✓ | ✓ | Strong | High | Pregnant Females | Strong | Viviparous Gestation |
| McCully-Phillips & Ellis, 2015, Griffiths et al., 2020, Breve et al., 2020, Silva & Ellis, 2019 | Eastern English Channel & Southern North Sea | *Mustelus asterias* | ✓ | ✓ | ✗ | ✓ | ✓ | Strong | High | Pregnant Females/ Females/ Mature Females/ Large Females | Strong | Viviparous Gestation |
| Kraft et al., 2023 | Sado Estuary, Portugal | *Dasyatis pastinaca* | ✗ | ✓ | ✓ | ✓ | ✓ | Strong | High | Mature Females | Moderate | Viviparous Gestation |
| Biais et al., 2017 | Bay of Biscay Shelf/ Shelf-edge | *Lamna nasus* | ✗ | ✓ | ✓ | ✓ | ✓ | Strong | High | Mature Females | Moderate | Viviparous Gestation |
| Biais et al., 2017 | Celtic Sea | *Lamna nasus* | ✗ | ✓ | ✓ | ✓ | ✓ | Strong | High | Mature Females | Moderate | Viviparous Gestation |
| Griffiths et al., 2020, Breve et al., 2020, Breve et al., 2016 | Bay of Biscay | *Mustelus asterias* | ✗ | ✗ | ✗ | ✓ | ✓ | Moderate | High | Females/ Mature Females | Moderate | Viviparous Gestation |
| Breve et al., 2016 | Eastern Tidal Baisn (Oosterschelde), Dutch Coast (Pupping) | *Mustelus asterias* | ✗ | ✗ | ✗ | ✓ | ✓ | Moderate | High | Mature Females | Moderate | Viviparous Gestation |
| Biton-Porsmoguer & Lloret, 2023 | The Roadstead of Brest, France | *Mustelus asterias* | ✗ | ✓ | ✗ | ✓ | ✗ | Strong | Moderate | Mature Females | Moderate | Viviparous Gestation |
| Mead et al., 2023 | La Graciosa Marine Reserve, Canary Islands | *Squatina squatina* | ✓ | ✓ | ✓ | ✓ | ✓ | Strong | High | Females | Moderate | Viviparous Gestation |
| Biais et al., 2017 | Western European Shelf | *Lamna nasus* | ✗ | ✗ | ✗ | ✓ | ✗ | Weak | Low | Neonates | Weak-Cannot Assess | Viviparous Gestation |
| Biton-Porsmoguer & Lloret, 2023, Silva & Ellis, 2019 | Bristol Channel | *Mustelus asterias* | ✓ | ✗ | ✗ | ✗ | ✗ | Weak | Moderate | Large Females/ Mature Females | Weak-Cannot Assess | Viviparous Gestation |
| Biton-Porsmoguer & Lloret, 2023 | Eastbourne | *Mustelus asterias* | ✗ | ✗ | ✗ | ✗ | ✗ | Weak | Moderate | Mature Females | Weak-Cannot Assess | Viviparous Gestation |
| McCully-Phillips & Ellis, 2015 | French Coast, Western English Channel | *Mustelus asterias* | ✗ | ✗ | ✗ | ✗ | ✗ | Weak | Moderate | Pregnant Females | Weak-Cannot Assess | Viviparous Gestation |
| Biton-Porsmoguer & Lloret, 2023 | Norfolk coast | *Mustelus asterias* | ✗ | ✗ | ✗ | ✗ | ✗ | Weak | Low | Mature Females | Weak-Cannot Assess | Viviparous Gestation |
| Biton-Porsmoguer & Lloret, 2023 | Normandy, France | *Mustelus asterias* | ✗ | ✗ | ✗ | ✓ | ✗ | Weak | Moderate | Mature Females | Weak-Cannot Assess | Viviparous Gestation |
| Biton-Porsmoguer & Lloret, 2023 | North Cornwall & Devon | *Mustelus asterias* | ✗ | ✗ | ✗ | ✗ | ✗ | Weak | Moderate | Mature Females | Weak-Cannot Assess | Viviparous Gestation |
| Silva & Ellis, 2019 | North of the Isles of Scilly and near Bann Shoal | *Mustelus asterias* | ✓ | ✗ | ✗ | ✗ | ✗ | Weak | Moderate | Large Females | Weak-Cannot Assess | Viviparous Gestation |
| Biton-Porsmoguer & Lloret, 2023 | Selsey Bill | *Mustelus asterias* | ✗ | ✗ | ✗ | ✗ | ✗ | Moderate | Low | Mature Females | Weak-Cannot Assess | Viviparous Gestation |

*This column highlights (in grey) when a cap was placed on the overall score based on the rules outlined in the risk of bias assessment.

| Reference(s) | Location | Species | R1 | R2 (Species) | R2 (Individual) | R3 (Species) | R3 (Individual) | Strength of Evidence | Quality Assessment | Evidence Cap* | Overall Evidence | Critical Area |
|---|---|---|---|---|---|---|---|---|---|---|---|---|
| Biton-Porsmoguer & Lloret, 2023 | South Devon | *Mustelus asterias* | ✗ | ✗ | ✗ | ✗ | ✗ | Weak | Moderate | Mature Females | Weak-Cannot Assess | Viviparous Gestation |
| Silva & Ellis, 2019 | The western Channel south of Selsey and Newhaven (in the western Channel:30E9, 30F0) | *Mustelus asterias* | ✓ | ✗ | ✗ | ✗ | ✗ | Weak | Moderate | Large Females | Weak-Cannot Assess | Viviparous Gestation |
| Silva & Ellis, 2019 | Near Eddystone (ICES rectangles: 29E5) | *Mustelus asterias* | ✓ | ✗ | ✗ | ✗ | ✗ | Weak | Moderate | Large Females | Weak-Cannot Assess | Viviparous Gestation |
| Silva & Ellis, 2019 | Northwest entrance of the western English Channel | *Mustelus asterias* | ✓ | ✗ | ✗ | ✗ | ✗ | Weak | Moderate | Large Females | Weak-Cannot Assess | Viviparous Gestation |
| Silva & Ellis, 2019 | The Banc des Langoustiers (ICES rectangles: 27E5) | *Mustelus asterias* | ✓ | ✗ | ✗ | ✗ | ✗ | Weak | Moderate | Large Females | Weak-Cannot Assess | Viviparous Gestation |
| Biton-Porsmoguer & Lloret, 2023 | The Thames Estuary in UK | *Mustelus asterias* | ✗ | ✗ | ✗ | ✗ | ✗ | Weak | Moderate | Mature Females | Weak-Cannot Assess | Viviparous Gestation |
| Biton-Porsmoguer & Lloret, 2023 | The Trieux Estuary, France | *Mustelus asterias* | ✗ | ✓ | ✗ | ✗ | ✗ | Weak | Moderate | Mature Females | Weak-Cannot Assess | Viviparous Gestation |
| Barker et al., 2022 | Irish Sea | *Squatina squatina* | ✗ | ✗ | ✗ | ✗ | ✗ | Weak | Moderate | Pregnant Females | Weak-Cannot Assess | Viviparous Gestation |
| Meyers et al., 2017 | Waters around the Canary Islands | *Squatina squatina* | ✗ | ✓ | ✗ | ✗ | ✗ | Weak | High | Pregnant Females | Weak-Cannot Assess | Viviparous Gestation |
| Barker et al., 2022 | Welsh coastal waters | *Squatina squatina* | ✗ | ✗ | ✗ | ✗ | ✗ | Weak | Moderate | Pregnant Females | Weak-Cannot Assess | Viviparous Gestation |

*This column highlights (in grey) when a cap was placed on the overall score based on the rules outlined in the risk of bias assessment.
